# Cell Settling, Migration, and Stochastic Cancer Expression Suggest Potassium Flux Per H^+^- Sensitive KCN Members Associated with Oncogenes and Histone/DNA Modulators Potentially Initiates pH-Reversal

**DOI:** 10.1101/2025.06.29.662144

**Authors:** Marie E. Beckner

## Abstract

Attraction of glioblastoma cells to potassium was suspected when cells clustered around dying cells and they migrated towards serum (high [K^+^]) and a positive potassium gradient. Potassium channel proteins (KCN family, 90 members) mediating altered transmembrane flux may provide K^+^ that releases H^+^ bound to inner membranes in cancer cells for cytosolic proton transfer (possibly Grotthuss) to extrusion sites. Cell settling and migration assay results led to collection of 70 studies, unbiased by authors for inclusion of KCN genes, that detected KCN differentially expressed genes (DEG). Of 53 KCN DEG found among 29 malignancies, 62.3% encoded H^+^-sensitive proteins. KCN DEG encoding H^+^-sensitive proteins were more prevalent in 50 studies involving one or more of 8 categories (7 oncogenes and histone/DNA modifiers) versus those with none, *p* = 0.0325. Pertinent genes for lactate outflow, etc. had relatively normal expressions. Brain tumors in REMBRANDT (database) showed altered expression of KCN genes encoding H^+^-sensitive proteins in glioblastomas versus less invasive oligodendrogliomas of patients on anti-seizure medications, with less *KCNJ16* /Kir5.1, *p* = 5.32 x 10^-8^. Altered H^+^-sensitive potassium flux via the KCN family downstream of oncogenes and histone/DNA modifiers putatively incites proton transfers for H^+^ release during pH reversal in cancer.

Graphical Abstract
Steps in K^+^/H^+^ Dynamics for pH Reversal in Cells. Step1. Potassium ions accumulate near cell membranes if (A) inward flow channels bring in more K^+^ than normal or (B) if the diffuse low outward flow channels are nonfunctional with compensation by channels in a few locations that are high flow requiring contributions of solvated potassium (K^+^-7H_2_O) from deeper cell regions. Step2. K^+^ reaches the inner cell membrane where H^+^ is bound diffusely to its negative charges. Step3. As K^+^-7H_2_O desolvates (for channel exit or membrane binding), the release of 7H_2_O may augment or form Grotthuss water wires near inner membrane surfaces. Step4. At the inner membrane K^+^ exchanges with H^+^. As H^+^ is released, it becomes H_3_O^+^ and diffuses into the cytosol. Step5. H_3_O^+^ enters a Grotthuss water wire that enables rapid proton transfer (conformational) to H^+^ exit sites. Step6. H^+^ leaves the cell with lactate or it exits via other H^+^ extruders. The cytosolic pH increases and the external pH decreases for pH reversal.

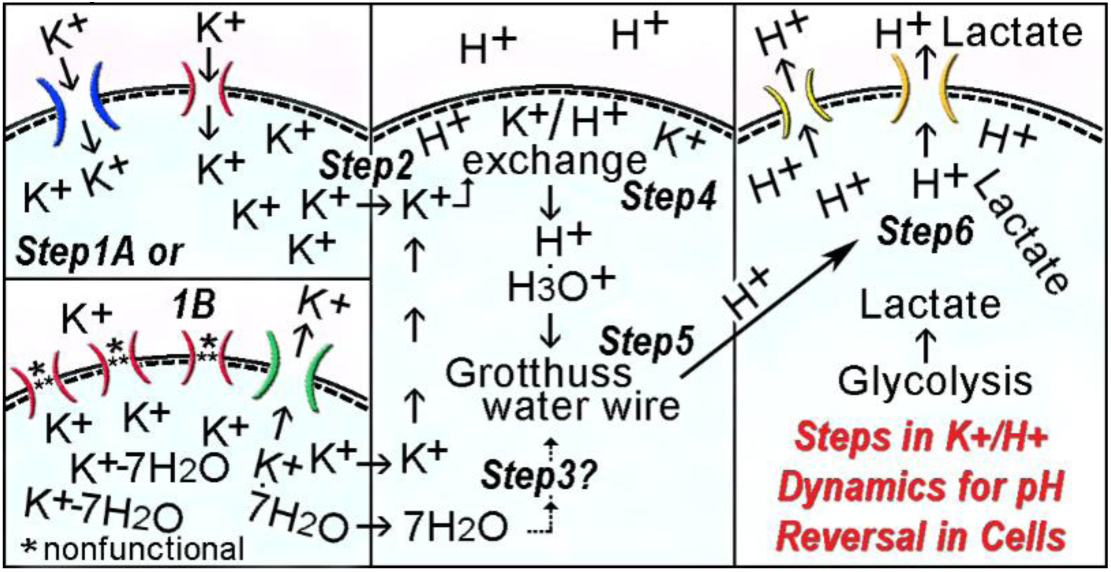

## 1. Introduction

### 1.1. General

Potassium flux is already known to maintain cell membrane potential in all cells, to assist volume changes in migrating cells, and permits neurons to fire action potentials as they release potassium that is buffered by glial cells. Potassium, as cytoplasmic monovalent cations, is now proposed to also exchange with fixed H^+^ on inner membranes to incite proton transfer and extrusion for shifts in pH that favor tumor cell survival and invasion. If so, then K^+^/H^+^ cation exchange may help to explain the attraction of glioblastoma cells (highest grade malignant glial tumors) to axons in CNS white matter tracts and peripheral nerve fibers [1,2] that release K^+^ via axonal action potentials as well as their growth factors. Experiments performed to study cell morphology and viability in a microaqueous environment suggested malignant cell attraction to K^+^. Glioblastoma cells tightly clustered around single dying cells, putatively releasing their cytoplasmic stores of K^+^. It had already been observed that glioblastoma cells migrated towards fetal bovine serum, known to have levels of 5-15 mM K^+^ [3]. This clustering behavior was similar to how glioblastoma cells rim necrotic foci (pseudopalisading) which is a typical histologic feature of these tumors [4–6]. Also, pseudopalisading glioblastoma cells have an enhanced migratory phenotype [6,7]. Additionally, in regard to reversal of pH, i.e. intracellular pH (pHi) is greater than extracellular pH (pHe), there are related metabolic features reported for benign astrocytic cells, such as (*i*) an increase of their intracellular pH with neuronal stimulation, possibly triggered by membrane K^+^ redistribution during depolarization [8,9]. Also, intracellular alkalinization detected in benign mammalian astrocytes has been attributed to their uptake of potassium released by repetitive neuronal firing of action potentials *in vivo* [10] as astrocytic cell membranes depolarize. Also, (*ii*) the rate of glycolysis (inhibited by lactic acid produced in hypoxia) is stimulated by changes in intracellular [H^+^] rather than [HCO_3_^-^] in astrocytes [11]. Cytoplasmic alkalinization in malignant and transformed cells occurring via efflux of protons resulting in extracellular acidosis has been attributed to mechanisms involving ion transporters, pumps, exchangers, etc. with the “active” ones requiring ATP [12–19]. Redistribution of K^+^ has been previously suggested as being a contributing factor in tumor cell alkalinization [12] and pH-sensitive ion channels, including those for K^+^, have been mentioned as “sensors” in descriptions of pH reversal [13,14,16,17]. Data in this study suggests the possibility of potassium flux as playing a role in glioblastoma cell behavior.

The large size of the KCN (potassium channel) family with its complex interactions among family members, an extensive complement of non-family regulators, functional reliance on contents of biological milieus, and a wide distribution across the genome serving as a relatively large target for genetic damage, were characteristics that supported further investigation of potential anomalous potassium flux in cancer through gathering data on this family in a stochastic manner. Retrieval of reported data unbiased for KCN gene expression in tumors allowed retrieval of data on differentially expressed genes (DEG) of KCN family members and associated non-KCN DEG in 70 studies of many types of cancer to provide an overall view of anomalous potassium flux in cancer. Also, a brain tumor database REMBRANDT [20,21], and a published whole genome analysis of 6 glioblastoma cell lines, Patil, et al. [22], were queried for data on KCN genes. The H^+^-sensitivity of a large proportion of the KCN family members detected and the high likelihood of K^+^ as a replacement cation for H^+^ suggest that potassium ion flux across plasma membranes or accumulation near them may play a role in the pH reversal commonly seen in malignancy. Deregulated, passive inward channel flux (without requiring ATP) or loss of normal diffusely distributed outward K^+^ flow (necessitating focally concentrated outward flow as needed) leading to accumulations of K^+^ ions at inner cell membranes with H^+^ bound to fixed anions potentially leads to K^+^/H^+^ exchange to release H^+^ as an initial step leading to pH reversal.

### 1.2. Cytosolic Proton Transfer Related to K^+^ and Hydration

The proposed K^+^ replacement of bound H^+^ would augment adjacent hydronium ion (H_3_O^+^) accumulation which can be seen at water-lipid interfaces [23], such as along inner cell membranes. This putatively sets the stage for pH reversal via subsequent steps. The mobility of unbound H^+^ (H_3_O^+^ at 36.2 x 10^-4^ cm^2^ V^-1^s^-1^) is very high compared to other univalent cations such as K^+^ and Na^+^ [24]. In immortalized human embryonic kidney cells, H^+^ diffusion through the cytosol occurred 60-fold faster than when limited to the inner plasma membrane surface, i.e. 0.6 μm*s*^-1^ versus 0.01 μm*s*^-1^ [25]. In cardiac myocytes a comparable high speed of cytoplasmic H^+^ distribution, 0.7 μm*s*^-1^, has been detected. This was based on using HCl in KCl (140 mM) solutions (pH 3.0) for acid diffusion studies with membrane acid transport simultaneously inhibited [26]. Another study has shown dissociation of HCl in a microaqueous environment so that the acid’s protons were each transferred to a neighboring water molecule and then another in clusters of HCl(H_2_O)_n_, where n=1 to 5 and 7. With the addition of a fifth and sequential seventh water molecule, the covalent H-Cl bond transitioned to the contact ion pair, H_3_O^+^Cl^-^, plus free hydrogens. The HCl(H_2_O)_5_ and HCl(H_2_O)_7_ clusters were spontaneously dissociated structures with contact ionic bonds [27]. Thus, H_3_O^+^ may be distributed to the cytosol after putative K^+^/H^+^ exchange on inner membranes and after spontaneous dissociation of acids (multiple types in cancer cells) within cytosols, possibly microaqueous in nature.

Cytosolic proton transfer has been previously attributed to small molecules, such as taurine, carnitine, carnosine, or homocarnosine, behaving as mobile buffers that shuttle H^+^ to enable cytosolic proton transfers [26]. However, proton transfers from H_3_O^+^ may also be enabled by a rapid mechanism consistent with behavior described originally by C.J.T. Grotthuss in 1806 [28] that fits with this scenario. Proton transfers may occur within the cytoplasm via H-bonded complexes, possibly involving Eigen and Zundel cations, H_9_O_4_^+^ and H_5_O_2_^+^, respectively, or by wires of protonated water molecules linked by interconversions in a manner that is consistent with the behavior of salts and water observed between electrodes in the early 1800s. According to the Grotthuss mechanism, due to interconversions, a proton that enters a chain or matrix of water molecules is not the same proton that exits elsewhere [24,28–34]. The conformational changes for proton transfer in water can occur as a rapid falling domino effect. Thus, a wide-reaching, flexible delivery of cytoplasmic protons to membrane extruders may result. Putative K^+^/H^+^ exchange that frees bound H^+^ from cell membranes would generate water cations because free H^+^ becomes H_3_O^+^ almost immediately which then potentially interacts with cytosolic chains of H_9_O_4_^+^ and H_5_O_2_^+^ or protonated water wires for distribution. Extended hydrogen bonded chains that undergo rapid conformational changes potentially mediate proton release distal from the point of proton entry much faster than by simple diffusion. H^+^ can be transferred in this manner to cell exit sites, such as exchangers swapping out H^+^, monocarboxylate transporters venting lactate that requires accompanying protons, one for one, exiting through gap junctions, etc. This type of rapid transfer of H^+^ in water has also been referred to as a “proton jump” [35]. Although in a previous study, the Grotthuss mechanism was not thought to mediate extracellular tumor acidity [36], it was not ruled out as occurring intracellularly. The Grotthuss mechanism describes a potential almost “everywhere, all at once” strategy for delivery of free protons to cell exit sites as needed. Therefore, putative inner membrane proton replacement by K^+^ to initiate formation of H_3_O^+^ is probably a highly regulated process in normal cells, consistent with the large, complex KCN family controlling K^+^ flux. Loss of regulation with differential expressions of KCN genes in malignant cells offers one approach to begin investigating the potential ionic dynamics of tumor cells related to pH reversal. Genetic damage of the KCN family members or altered regulation by activated oncogenes and/or modified histones/DNA putatively plays a role in pH reversal for tumor cell survival in glycolytic cells producing lactate if it cannot be removed in another way (such as by forming intracellular glycogen).

Also, potassium has a relationship with water that is important. A lack of water due to poor circulation, cell crowding, pressure from fibrotic, stiff surroundings, etc., especially during invasion, produces generalized microaqueous conditions within cancer cells. Regional hydration may be possible by redistribution of water molecules loosely associated with solvated ions, such as potassium, K^+^-7H_2_O. Redistribution of water via its association with potassium may enhance Grotthuss type protonated water structures close to the inner surface of cell membranes but this is unknown.

Volume changes related to ions are known to occur in malignant cells during division and migration. Protrusion of pseudopodia or invadopodia (increased volume in cell projections) and shrinkage of their retracted rear cell bodies that are accomplished with oscillations of cell membrane potential and osmotic changes via KCl uptake and release per potassium and chloride channels, along with aquaporin participation, and possibly with K^+^ refilling/recycling [37–42]. The spontaneous dissociation of acids generated in a microaqueous environment undergoing changes in the number of water molecules available for solvation of acids into ion pairs [27], along with rapid proton transfers via the Grotthuss mechanism, putatively enhance proton delivery to the leading edge of migrating cells for efflux to preserve glycolysis from lactate inhibition. Pseudopodia are glycolytic as indicated by proteomic analysis [43]. Also, enhanced proton release from cancer cells leads to extracellular acidosis that aids tumor invasion via activation and induced expression of metalloproteases for matrix degradation [44–50]. Thus, tumor cells can escape poorly perfused or crowded, microaqueous tissue regions via pH reversal relying on proton release putatively following K^+^/H^+^ exchange as the initiating step.

### 1.3. Need for Proton Extrusion in Cancer

The metabolism of cancer cells produces excess protons [51]. In addition to acids produced by metabolic pathways, free radicals generated are also acidic [52–54]. Extrusion of protons avoids cytoplasmic acidosis within tumor cells, so this not only protects glycolysis via lactate release but also prevents cytochrome c-mediated activation of caspase-3 to stop apoptosis [55], protects gap junctions to maintain cell to cell exchange of ions and metabolites [56–59], and increases the activity of ATP citrate lyase (ACLY) that connects lipid and carbohydrate metabolism. The optimal pH of ACLY is in the 8.5 - 8.7 range [60,61]. ACLY’s removal of citrate (inhibitor of glycolysis) is protective in glioblastomas [62]. ACLY also regulates histone acetylation in response to growth factor stimulation [63,64]. Thus, proton removal from the cytoplasm favors cell fitness in tumor cells [65] as well as enabling migration for their escape from adverse conditions.

### 1.4. Complexity of K^+^ Flux

As mentioned earlier, direction (inward or outward) and strength of K^+^ channel flow is complex. The KCN family has more than 90 members [66,67]. Complexity is also due to numerous regulators (in addition to those encoded by KCN genes) present in biological milieus. Regulators, gating agents, and cofactors, include polyamines, voltage, H^+^, Ca^2+^, Mg^2+^, Zn^2+^, Na^+^, ATP, miR-133b, reactive oxygen species (ROS), reactive nitrogen species, reactive sulfur species, nitric oxide, hydrogen sulfide, hypoxia, hyperglycemic levels of glucose, nucleotides, protein kinase A, protein kinase C, protein kinase G, guanosine-triphosphate-binding proteins (G-proteins), regulator of G protein signaling (RGS) proteins, phosphatidyl inositol 4,5 bisphosphate (PIP2), steroids (estrogen, testosterone, etc.), cholesterol, sulfonylurea, β-hydroxybutyrate, hypoxia inducible factor 1α, the 14-3-3 protein, β-coat protein, dipeptidyl aminopeptidase-like proteins, the neurotransmitter GABA, NAD(+), NADP(+), NADH, NADPH, glutathione, glutathione disulfide, mechanical stimulation, cell cycle progression, products of *LINGO1, 2*, and BKγ1-4 encoded by *LRRC26*, *52*, *55*, and *38* [68–90]. Also, 19% (or more) of proteins targeted by small-molecule drugs used to treat numerous diseases include ion channels [91–93]. Thus, the complexity of K^+^ channel flux hinders planning studies to identify specific regulatory mechanisms lost in cancer.

Complexity is further enhanced because the ROS affecting potassium channels are generated via multiple sources such as (*i*) mitochondrial respiratory chain electron transport redox reactions [84,94–97], as well as (*ii*) the NOX family (6 members) of NADPH oxidases [71,94,95,98]. ROS from these sources can reach the cytoplasm [95]. Colon carcinoma cell lines with acid-resistant growth showed significantly upregulated mRNA levels of *NOX1* encoding NADPH oxidase 1 [99] that produces ROS. Interestingly, isolated intact respiring rat brain mitochondria increased ROS production when their medium was alkalinized which raises the possibility that this could also happen in alkaline cytosol of cancer cells as pH reversal develops [100].

The widespread chromosomal distribution of the 90 (or more) KCN genes plus those of the numerous non-family regulators suggests that failsafe redundancies for genomic damage may exist that increase the chances of selecting cells that can maintain levels of K^+^ for survival but not in a manner that is optimal for differentiation and normal activity. Numerous genomic/expression scenarios selected in rapidly dividing tumor cells may work around mutational losses or gains with adaptations so that cells maintain their K^+^ levels for supporting membrane integrity and are putatively also able to exchange with fixed H^+^ on inner membranes in invasive cells to incite pH reversal. Not only do the monocarboxylate transporters, MCT1 and MCT4, transport negatively charged lactate molecules coupled one-for-one with protons across the cell membrane [101], protons can also be extruded from cancer cells via Na^+^/H^+^ exchanger family members, ATP driven pumps, etc. as detected in numerous studies [102] and in earlier cited studies. Therefore, pertinent genes among the non-KCN DEG are also of stochastic interest in this study.

Mediators of potassium flux, despite their complexity, and the relationship of K^+^ and H^+^ at cell membranes should be clarified. K^+^ accumulation due to a lack of normal diffuse K^+^ efflux or buildup of K^+^ around large outwardly conducting potassium channels, could putatively exchange with bound protons on inner membranes or it could happen with deregulated inwardly channeled K^+^ flux (possibly with K^+^ being recycled for refilling as needed if efflux becomes excessive). These events with K^+^ could occur without affecting the functions of key structural proteins that would be seen if Na^+^ was used [103]. Although both potassium and sodium are the available monovalent cytoplasmic cations to potentially replace membrane bound H^+^, there should be a preference for potassium due to its weaker binding of water compared to sodium ions. Less energy is required to displace water from potassium than sodium [104,105]. K^+^ ions permeate their membrane channels in a desolvated (‘naked’) state rapidly so at this point they release their associated water molecules. Also, there is exclusion of smaller Na^+^ ions that have stronger water binding from the channels selective for potassium [106–108]. Also, compared to Na^+^, K^+^ is better at maintaining solubility of cell components containing carboxylates and phosphates to help explain its higher concentration in the cytoplasm [105,109]. Numerous KCN family scenarios may be involved with anomalous potassium flux in cancer. Deregulated diffuse, passive K^+^ entry, such as the *KCNJ*/Kir (inwardly rectifying, also bidirectional, flux), or disrupted K^+^ outflow via KCN family members, such as the numerous members of the Kv (voltage dependent) KCN subfamily and the *KCNK*/K2P (tandem two pore “leak”) subfamily members, may be important. Also, Ca^2+^ activated potassium channel families, *KCNM*/BK for *MA1* plus other members that regulate it, and the *KCNN*/SKs and *KCNN4*/IK family members that mediate outward flow, with BK and IK providing large and medium flux, respectively, likely play important roles due to their relatively high flow rates. High flow rates may indicate localized versus diffuse K^+^ release and in addition these channels can be affected by the release of calcium from cancer cells. However, all types of K^+^ transits may be involved and a role for Na^+^ in releasing H^+^ cannot be ruled out given the abnormally high levels of Na^+^ found in tumor cells [110–112]. Additionally, some K^+^ channels are influenced by Na^+^.

Loss of normal cellular responses to pH via loss of KCN channels that function with H^+^-sensitivity may alter cells in cancer. Malignant cell behavior suggests that tumor cells respond opportunistically to sources of K^+^, thus achieving levels of K^+^ ions at inner membrane surfaces or flow rates that force K^+^/H^+^ exchanges that initiate pH reversal to improve tumor cell survival and invasion. Deregulated potassium uptake by glioblastoma cells may occur from the available sources *in vivo* as briefly mentioned earlier. These (with additional information) include (*i*) neurons and axons firing action potentials where K^+^ is released to maintain electroneutrality, (*ii*) dying cells, including red cells (K^+^, 80 - 120 mmol/L) and necrotic tissue releasing cytoplasmic K^+^ [113–116], and (*iii*) damaged blood vessels leaking plasma (potassium, 3.8 - 5.0 mmol/L [117]). Importantly, deregulation via loss of a K^+^ channel is illustrated by showing in an experimental mode that the loss of H^+^-sensitive *KCNK1* (among a small group of genes identified) was advantageous for breast cancer cell survival of induced necrosis l [118].

### 1.5. Biological Milieu Constituents to Consider in Studying K^+^ Flux

Biological milieus contain many factors regulating potassium channels as described earlier. More details to consider in studying these channels in cancer are listed here. Serum includes calcium (2.30 - 2.74 mmol/L), magnesium (0.74 - 1.23 mmol/L), phospholipids (1.50 - 3.80 g/L), sodium (136 - 142 mmol/L, adult plasma value) [117], and polyamines (0.1 - 1.0 μM, fetal bovine serum [3]). Magnesium and polyamines (positively charged) are important gating molecules of inwardly-rectifying *KCNJ*/Kir family channel members [119]. Removing fetal bovine serum from the medium of cultured fibroblasts resulted in decreased potassium intracellularly [120]. Also, fetal bovine serum lost the ability to depolarize the resting membrane potential of airway smooth muscle cells after dialysis against distilled water but its freeze-dried dialysate retained this ability after reconstitution [121], consistent with potassium as the responsible constituent. Elevations of polyamines have been reported in cerebrospinal fluid in instances of brain tumors [122] and are in the tumor-associated myeloid cells that infiltrate glioblastomas [123]. Also, polyamines are formed when L-arginine, released from apoptotic bodies, is metabolized by Arg1 [124]. Other agonists and modifiers in biological milieus within regions of necrosis include heme and other hemoglobin break-down products [125,126]. Tumor associated macrophages are influenced to promote malignancy (increased expression of *ARG1, VEGF*, *TGFB*, etc.) by lactate, presumably shuttled from cancer cells, such as seen in transitional cell carcinoma of the bladder, gastric cancer, and others [127–129]. As mentioned earlier, many types of tumors outside of the CNS exhibit perineural invasion [130–133].

There may be reciprocal gain of potassium by tumor cells from nerves and axons as the tumor cells provide lactic acid back to neurons and axons as energy substrate for a mutually beneficial metabolic interchange. If constituents of biological milieus modulate K^+^ transits indirectly via their effects on downstream potassium channel activity, then sources of extracellular potassium may attract tumor cells to fulfill their needs. Although specific sources of K^+^ *in vivo* have not been generally appreciated for influencing the behavior of cancer cells, they may be important as loss of regulation required for complex normal cell functions occurs in dedifferentiation.

### 1.6. Highlighted Findings in This Study

Fetal bovine serum (with and without hepatocyte growth factor (HGF)) had been seen previously to promote glioblastoma cell migratory behavior, including protrusion of pseudopodia through 3 μm filter pores, migration of whole cells through 8 μm filter pores, and invasion through a Matrigel matrix and rat brain tissue slices, often under glycolytic conditions [43,62,134–136]. Glioblastoma cell clustering around dying cells and their migration to serum, with compelling background information on astrocytes, suggested that anomalous potassium flux could play a role in tumor cell behavior. Cell migration of multiple glioblastoma cell lines to serum among eight tested in this study and the response in a separate assay to a positive gradient of potassium supported abnormal membrane flux or lack of it via some of the potassium channels as a putative aid for tumor cells.

The large scope of K^+^ channel candidates for loss or gains in expression and the potential influence of numerous known modulators were daunting. However, stochastic analysis of numerous published studies to gather unbiased data was reasonable to survey as many contributing factors as possible at this point. Incidentally detected KCN DEG and related non-KCN DEG in genomic landscapes of experimental cancer scenarios and databases permitted a search for pertinent functional similarities among the DEG of the large KCN family. Meta-analysis of 70 malignant (or models related to cancer) expression studies found that the studies fell into 8 categories of 7 oncogenes and histone/DNA modifiers. These categories were determined by the authors of the expression (E), E1-E70, studies if related DEGs were also detected in any of the 70 studies. The studies included 29 types of malignancy (breast, lung, brain, etc.) that incidentally detected one or more KCN DEG. *KCNMA1* and *KCNN4* were the most frequently detected KCN DEG. The encoded products of the 53 detected KCN DEG are predominantly H^+^-sensitive (62.26%) and 52 (74.3%) of the 70 studies identified at least one H^+^-sensitive KCN DEG. There was a significant propensity for KCN DEG to encode H^+^-sensitive family members in the studies among the 8 categories of 7 oncogene and histone/DNA modifier related scenarios compared to the 20 studies in “no category”. A concurrent search of the National Institutes of Health REpository of Molecular BRAin Neoplasia Data (REMBRANDT) database for expression levels of KCN genes in invasive glial brain tumors (glioblastomas versus oligodendrogliomas that are less invasive) of patients on anti-seizure medications showed a significant decrease in *KCNJ16* and trends in three other KCNs also encoding H^+^-sensitive proteins in glioblastomas. Correlation with a genomic study of 6 glioblastoma cell lines [22] confirmed relatively decreased RNA expression of *KCNJ16* among KCN genes and provided additional relevant genomic data. Altered expression of KCN family members, possibly with compensatory activities of other KCNs and non-KCN factors, may provide quantities of K^+^ sufficient for preservation of cell membrane integrity and putative K^+^/H^+^ exchange on inner membranes to release bound H^+^. As an initial step K^+^/H^+^ exchange may incite pH reversal and invasive behavior.

## 2. Materials and Methods

### 2.1. Materials and Cell Culture

All chemicals and reagents were from Sigma-Aldrich (RRID:SCR_008988), St. Louis, MO, USA, unless otherwise stated. Anti-Met was from Cell Signaling Technology, Beverly, MA, USA, Cat# 4560, RRID:AB_2143887), 1:1000 dilution, for the beta Met subunit (145 kDa). Diff Quik (Allegiance, McGaw Park, IL, originally developed by Harleco, now a product line of Sigma-Aldrich (RRID:see above) is sold as Hemacolor Solutions I, II, and III. It is also sold by many medical suppliers with reference to Harleco and/or the cytology literature [137]. A172 (RRID:CVCL_0131), C6 (TKG Cat# TKG 0242, RRID:CVCL_0194), F98 (ATCC Cat# CRL-2397, RRID:CVCL_3510), LN18 (RRID:CVCL_0392), LN229 (RRID:CVCL_0393), T98G (RRID:CVCL_0556), U87 (RRID:CVCL_0022), and U373 (RRID:CVCL_2219) glioblastoma/astrocytoma cell lines (human (A172, LN18, LN229, U87, U373) and rat (F98 and C6)) were obtained from the American Type Culture Collection, Manassas, VA, USA RRID:SCR_021346. No cell lines were generated from live animals for this study by the author. The cells were maintained in Minimal Essential Media (MEM) Eagle(138) or alpha-MEM (Cellgro, Mediatech, Herndon, VA, USA) with 10% fetal bovine serum, FBS, (Invitrogen, Carlsbad, CA, USA).

### 2.2. Statistics Section

Pearson product-moment correlation, R, also termed cor (with *p* values for a true correlation being greater than 0), the Student’s t-Test, paired and unpaired, and Fisher’s Exact Test for Count Data, in either Excel Statistics (RRID:SCR_017294) or the R Project for Statistical Computing (RRID:SCR_001905) were used.

### 2.3. Databases

For general gene information, the National Institutes of Health database, https://www.ncbi.nlm.nih.gov, the Online Mendelian Inheritance in Man (OMIM) database, OMIM (RRID:SCR_006437), https://omim.gov, and GeneCards (RRID:SCR_002773) https://www.genecards.org/ were used. GeneCards was also used to obtain protein-protein interaction data. The Ensembl https://www.ensembl.org/ database (RRID:SCR_002344) was used to obtain information on the structure of genes, including features of exons and introns. The National Library of Medicine’s PubMed database, https://pubmed.ncbi.nlm.nih.gov/, was searched for the numbers of publications available using 2 and 3 word searches for subjects of interest. The Online Mendelian Inheritance in Man (OMIM) database was searched for KCN family member genes expressed in the nervous system with exclusion of genes uniquely expressed in heart, kidney, and other non-neural sites, and to identify their chromosomal loci (March 2019). The REpository for Molecular BRain Neoplasia DaTa (REMBRANDT) (RRID:SCR_004704), originally at the NIH, became part of the Georgetown Database of Cancer (G-DOC), https://sites.google.com/georgetown.edu/g-doc/home or via the G-DOC Hub platform at https://gdochub.georgetown.edu/ as a project of the Georgetown Lombardi Comprehensive Cancer Center, Washington, D.C., designed to provide translational research tools to the scientific community [20,21]. It was queried for gene expression on microarrays of KCN family members, selected genes important in cancer and brain tumors, and housekeeping genes. Medians of reporter probes for each gene were available in Adobe Flash (RRID:SCR_017258) readouts through 12/31/2020 (date of last service). Genomic sequence and expression results of 6 glioblastoma cell lines were obtained from the Supplemental Materials of Patil et al. [22].

### 2.4. Ethics Statement

No human subjects or animal experiments were used by the author in this study. Only commercially available human and rat cell lines were used i*n vitro* for laboratory studies. Published medical science manuscript information, mostly open access, with the authors of the studies following the guidelines of each their institutions, was also used for data.

### 2.5. Cell Settling with Fluid Removal Assays

Human glioblastoma cells (U87, LN229, A172, T98G) were maintained in routine culture conditions. On the day of each assay, cells were detached with trypsin and allowed to recover for 4 hours in MEM with 10% FBS at 37° C in a CO_2_ tissue culture incubator. For each assay, a Cell Settling Chamber Kit (Neuro Probe, formerly Gaithersburg, MD, USA, Ph: (301) 417-0014), with 8 wells, each 7 mm in diameter, was assembled. A provided glass slide, 1 mm thick, 25 x 75 mm, was placed in the recessed bottom plate. A provided pre-punched piece of blotting paper, pre-moistened with PBS, a gasket, and the upper plate were stacked on top of it. Thumbnuts fitted on the top plate were securely hand tightened. Cells were centrifuged (500 rpm, 5 min) and resuspended with gentle pipetting, 500,000 cells per ml in culture media with 10% FBS. Cell suspensions were added, 50 microliters per well. After 30 minutes with media visibly absorbed out of the wells by the blotting paper, the chamber was disassembled, the slide was removed and stained with Diff Quik solutions, I, II, and III followed by 3 quick rinses in tap water in beakers. At least two replicate wells per cell line were prepared. Stained cells were examined microscopically. Cell vacuoles in U87 cells (single and clustered) were counted in the settled cellular monolayers. When vacuoles collectively had a diameter of at least one third of an average nuclear diameter in the same 100X microscopic field, the cell was counted as positive for vacuolization (surrogate indicator of degeneration). Cells were analyzed in a midline sweep of the 100X microscopic fields across the 7 mm width of each well. Pearson product moment correlation coefficient, R, determinations, were performed for the percentage of cells vacuolated versus the number of cells in each cluster. Morphological features, such as cell clustering, were noted. An Olympus BH2 microscope (Olympus Corp., Lake Success, NY, USA) with an Olympus FV II digital camera using Olympus Microsuite Five Software for Imaging Applications (Soft Imaging System, Lakewood, CO, USA) was used to capture the black and white images. An American Optical Spencer Microscope (Buffalo, NY, USA, sold to Reichert Technology, then Leica Microsystems, Inc. (Deerfield, IL (RRID:SCR_008960)) with an OptikamB9 Digital Camera 10 MegaPixels (Optika, www.optikamicroscopes.com, Via Rigla 32, 24010 Ponteranica (BG), Italy) with Adobe Photoshop v5.1 and Adobe Photoshop Elements 2024 for Windows (Adobe Systems Incorporated, San Jose, CA, USA (RRID:SCR_014199) was used to generate the color microscopic images.

### 2.6. Cell Migration Assays

Cells of A172, C6, F98, LN18, LN229, T98G, U87, and U373 glioblastoma cell lines were maintained in normal tissue culture conditions with half volumes of culture media changed one day prior to each assay. Confluent cells were detached with trypsin on the day of assay and allowed to recover in media with 10% FBS for 2 hours at 37°C in a CO_2_ tissue culture incubator. Cells were centrifuged (<500 rpm, 5 min) and resuspended at 2 million/ml with gentle pipetting in media for migration, MEM (includes 5.4 mM KCl [138] but it is 5.3 mM according to Thermo Fisher among its inorganic salts), with 0.1% bovine serum albumin (BSA). Chemoattractant/chemokinetic stimulants, recombinant hepatocyte growth factor (HGF) and/or FBS were added to the bottom wells (29-30 μl total volume) as indicated in Figure 2. The 48-well modified Boyden Chambers (Neuro Probe) were assembled with porous (8 μm diameter pores, 1,000 pores/mm^2^, laser generated), air-dried polycarbonate filters, 8 μm thick (Neuro Probe), coated with 0.01% porcine gelatin (solution of 1 ml 10% acetic acid, 1 ml 1% gelatin (fully dissolved, no floating globules) from refrigerated stock (10%, gently warmed to liquify), and 98 ml H_2_O). One corner of each filter was diagonally cut off sufficient for orientation before each assay. Cell suspensions were added to the upper wells, 60 μl each well. The chambers were incubated at 37°C for 5 hours in a CO_2_ (5%) tissue culture incubator. Upon completion of the assay, each filter (easily disturbed by air flow) was removed by lifting one end with a pair of fine, curved tip tweezers, and then clamped into a large square plastic clip at the other end of the filter and then releasing the tweezers without disturbing any cell pellets. The filter attached to the large clip was stained with Diff Quik solutions (I-III) by hanging the filter by its attached clip on a metal weighing spatula placed over tops of beakers with Diff Quik solutions, followed by rinses in 3 different beakers of water using quick dips, and then carefully placed on a large glass slide (2” x 3”) using tweezers and the clip with migrated cells on the downside of filters. Unmigrated cells were gently removed from the upper side of a filter with cotton swabs or Kimwipes so that only the stained migrated cells remained, located between the filter and glass slide. There were 4 replicates per data point unless pre-assay cell harvests only allowed fewer replicates. Slides with migrated cells were digitized using a regular scanner (HP OfficeJet Plus 8715 printer/scanner (Hewlett Packard (formerly RRID:SCR_011873), Spring, TX, USA). Densitometry (UN-SCAN-IT Gel Analysis Software, 7.1 (RRID:SCR_017291) Silk Scientific, Inc., Provo, UT, USA) with correction for background filter staining density to quantify cell migration results. An Epson Perfection 2450 PHOTO (Epson America, Long Beach, CA, USA) was also used for obtaining some of the images of migration assays. The first two rows of migrated cell pellets, digitized for densitometry, were the results used to calculate cell migration specifically to HGF, i.e. HGF cell migration, defined as the 2^nd^ rows of cells which had migrated to HGF divided by the 1^st^ rows of cells representing background (unstimulated) migration because they migrated to media without the added chemoattractants, HGF and/or FBS. HGF cell migration results are stated as times(X) the corresponding unstimulated, background cell migration for each cell line. Additional migration assays of U87 cells evaluated the motogenic/chemokinetic properties of FBS and HGF by placing them in either the bottom or both top and bottom wells and also separately, MEM versus Dulbecco’s PBS was evaluated in the bottom wells in a small dose curve of FBS with Migration media (Dulbecco’s PBS (contains 2.7 mM KCl, 1.8 mM KH_2_PO_4_, 137.9 mM NaCl, 10 mM Na_2_HPO_4_) with added 0.210 mM MgCl_2_, 0.243 mM CaCl_2_, 5 mM pyruvate, 5.551 M glucose, and 0.1% bovine serum albumin) in all of the top wells.

### 2.7. Immunoblot of Met, Receptor for HGF

One dimensional gel electrophoresis was performed for immunoblotting. Lysates of unmigrated cells from 8 cell lines, equalized for total protein content (10 μg per lane), were electrophoresed in separate lanes of 10% polyacrylamide gels under reducing conditions. Protein standards (MagicMark, MultiMark, Invitrogen (now merged with Thermo Fisher Scientific (RRID:SCR_008452)) were loaded 8 μg per lane. The gel was stained with Coomassie blue (Novex Colloidal Blue Stain Kit, Invitrogen, now merged with Thermo Fisher Scientific (RRID:SCR_008452)), as shown, and destained with tap water. Gel contents were transferred onto polyvinylidene difluoride membranes (Invitrogen, now merged with Thermo Fisher Scientific (RRID:SCR_008452)), blocked (Detector Block, Protein Detector Western Blot Kit Lumi-GLO System, Kirkegaard & Perry Laboratories, Gaithersburg, MD, USA, now merged with SeraCare Lifesciences:(RRID:SCR_004535)) and reacted with anti-Met (1:1000), reactive with a 145 kDa band. The secondary antibody 1:1000 (Kirkegaard & Perry Laboratories, now merged with SeraCare Lifesciences:(RRID:SCR_004535)) was horseradish peroxidase-labeled anti-mouse. Immunoreactive bands were visualized via horseradish peroxidase’s conversion of a luminol-based solution to produce chemiluminescence, and the reactive blot was scanned to generate a digital image. An Epson Perfection 2450 PHOTO was used for obtaining digital images with densitometry obtained using UN-SCAN-*It* gel, early version 5.1.

### 2.8. Differentially Expressed Genes of Potassium Channels

Potassium channel (KCN) family members were found as differentially expressed genes (DEG) in cancer genomic landscapes described in 70 studies (1301 authors) recently published (2007-24) in 15 journals through subscriptions and open access. The KCN DEG were detected with non-KCN DEG in at least one published figure (heatmap, Venn diagram, volcano plot, etc.) or table (none from Supplemental Materials of the manuscripts) in each publication. Detection of one or more KCN genes in a group of genes designated as DEG by the authors (with no initial focus on KCN genes) was used for selecting these studies. Types of studies included mostly common malignancies, such as breast, lung, gastrointestinal, brain, bone marrow, liver, soft tissue sarcomas, etc. Categories (determined after all E1-E70 studies were completed) were according to descriptions by the authors of each paper with the caveat that there had to also be a relevant DEG reported in at least one of the E1-E70 studies.

### 2.9. Expression of Potassium Related Genes in Glial Tumors

REMBRANDT (G-DOC) contained tumor expression data on 67 oligodendrogliomas (Oligo) and 220 glioblastoma (GBM) patients, including subgroups treated with anti-seizure (AS) medications, i.e. 24 Oligo-AS and 75 GBM-AS. Gene expressions were determined by reporter probes on microarrays with their median values provided. Differences in median values for multiple probes (when available) were found with t-tests, paired via reporter probes for the two comparison groups when possible, and otherwise, unpaired t-tests were performed. Housekeeping genes (HKG) that lacked relevance for glial tumors via literature searches were also analyzed for comparisons. Additionally, Oligo-minus-GBM (OMG) differences were calculated for each probe. OMG differences for each gene of interest were compared to the OMG differences for HKG (*B2M*, *PPIA*, *RRLP0*, and *YWHAZ*). The *t*-tests were performed for unpaired samples with unequal variance in OMG comparisons. Published studies [22,139,140] were used for confirmation of REMBRANDT results.

## 3. Results

### 3.1. Settled Glioblastoma Cells with Fluid Removed

At the end of each cell settling assay, the extracellular fluid had been wicked away. Cell groups could be seen macroscopically in some as viewed in Appendix A (Figure. A1). For each cell line (A172, LN229, T98G, and U87) a striking feature was the clustering of robustly viable tumor cells around single degenerating cells of the same lineage on slide surfaces in low numbers. Cell surfaces were exposed to ambient air, glass, or surfaces of other cells by the end of the assays. The tight clustering of viable cells suggested an opportunistic attraction to the dying single cells by possibly taking up released contents from moribund cells to enhance fitness. Although the tint of staining varied, some viable cells clustered around single dying cells displayed basophilic cytoplasmic staining (Figure 1A-B). Although some faint, thin connecting strands between cells may have been present, mostly there was a narrow linear open space between the apposed cells, best seen in black and white photographs. Cells, single and in groups without visible degenerating central cells, and scattered debris were also seen. Cytoplasmic vacuolization was most prominent in the U87 cells (Figure 1A-C). The number of vacuoles was highest in single cells and decreased as the size of cell clusters increased. Representative U87 cells, single and in groups, were layered together into one image with a broad white stroke added around each layer for delineation and contrast with the added pink background (Figure 1C). In counts of vacuolated settled cells performed on the cell monolayers from two separate wells, 15% and 72% of single cells were vacuolated whereas vacuolated cells were only 11% and 18% in cell groups comprised of greater than twenty cells in the 1^st^ and 2^nd^ wells, respectively. Pearson regressions of the cells in clusters with vacuoles for the two wells had significant R^2^ values of 0.266 (*p* = 0.0410) and 0.544 (*p* = 0.0062), 1^st^ and 2^nd^ wells, respectively, as seen in scatterplots (Figure 1D). The cells in clusters exhibited less vacuolization as a sign of less degeneration possibly due to sharing their cytoplasmic contents as needed and also possibly due to the contributions of contents from dying cells of the same lineage.

**Figure 1.**
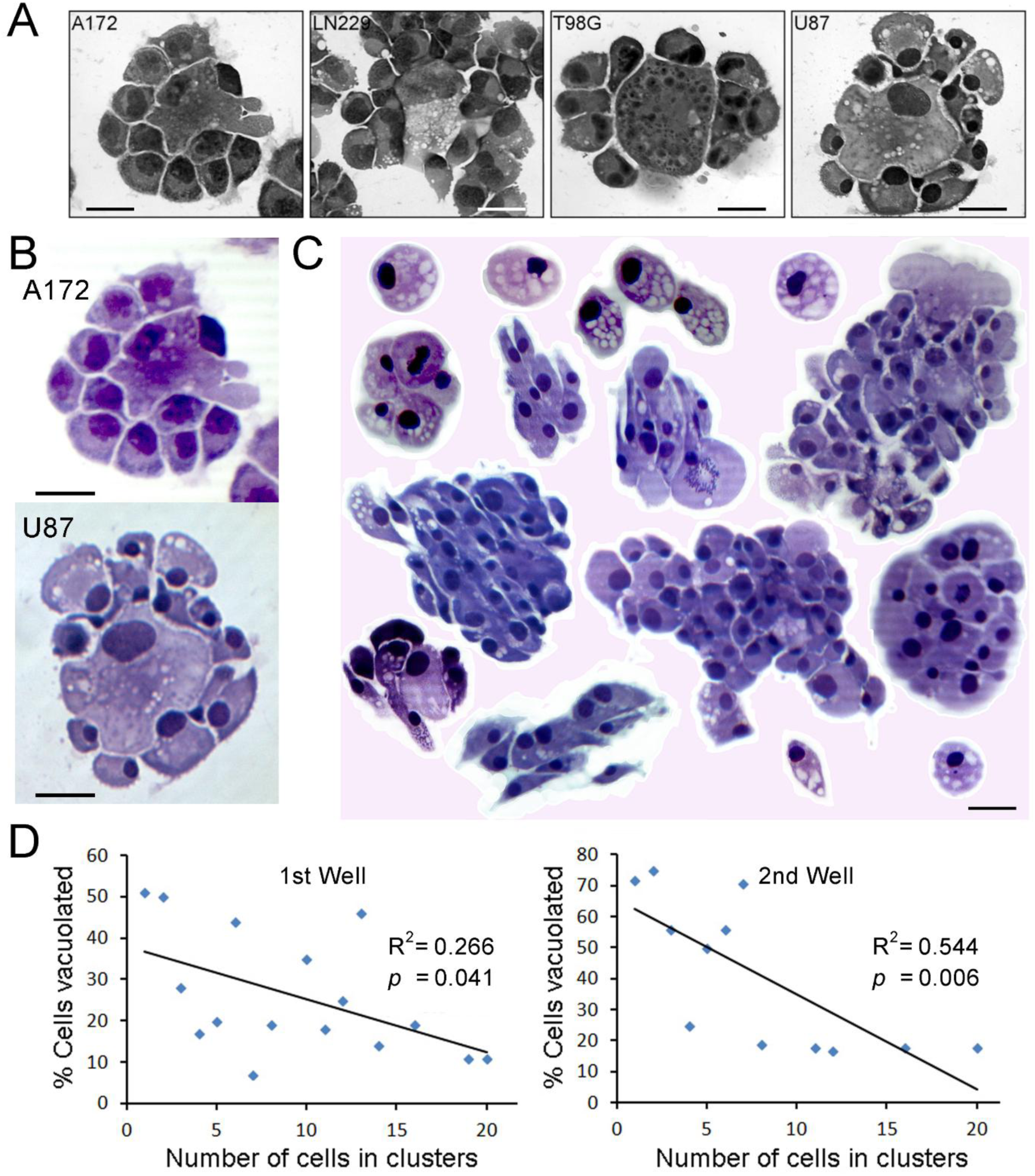
Settled glioblastoma cells deprived of extracellular fluid. (A and B) The morphology of each cell line (A172, LN229, T98G, and U87) after cell settling with fluid wicked away, included a few clumps of robust appearing, viable tumor cells clustered around single degenerating cells of the same lineage. Cell surfaces, after settling and withdrawal of suspension media, were exposed to ambient air, glass, or surfaces of other cells. Although the tint of Diff Quik staining varied, some viable clustered cells’ cytoplasm stained relatively basophilic. The black and white photographs suggest some thin connections between cells crossing the narrow linear open spaces separating them. Single and grouped cells without a visible degenerating central cells and scattered cell debris were also seen in wider views. (C) Cytoplasmic vacuolization, most prominently seen in U87 cells, is shown in the representative single and grouped cells that were outlined with digital white strokes around each merged layer for delineation and contrast with the added pink background. More basophilic cytoplasmic staining with Diff Quik seen in some cell groups would be consistent with an alkaline pH but no pH measurements were made. The numbers of vacuoles were highest in single cells and decreased as the numbers of cells in the cellular groups increased. (D) In counts performed on the cell monolayers from two separate wells, 51% of 75 and 72% of 25 single cells were vacuolated, 1^st^ and 2^nd^ wells, respectively. However, vacuolated cells in groups were only 11% of 122 and 18% of 39 cell groups comprised of greater than twenty cells, 1^st^ and 2^nd^ wells, respectively. As shown in the 2 scatterplots, Pearson regressions of the numbers of cells contiguous in groups or clusters that had vacuoles had R^2^ values of 0.266 (*p* = 0.0410) and 0.544 (*p* = 0.0062), 1^st^ and 2^nd^ wells, respectively. Scale bars, 25 μm.

### 3.2. Cell Migration with HGF and Serum Chemoattraction

Earlier studies of U87 and LN229 glioblastoma cell lines and fresh glioblastoma cells from a clinical specimen reported chemoattraction to HGF and serum [43,62,134–136]. Testing additional glioblastoma cell lines for migration to HGF and FBS, their chemokinetic effects (more extensively than previously), comparing cellular levels of HGF’s receptor, Met, and including a comparison of MEM versus Dulbecco’s PBS as the media in bottom wells for chemoattraction in a FBS dose curve with U87 cells are reported here. Boyden chamber assays (48 well chambers), 3 per cell line, for 2 rat astrocytoma cell lines (C6 and F98) and 5 human glioblastoma cell lines (T98G, LN18, U87, LN229, and U373) showed cell migration to HGF alone as a chemoattractant (row 2), HGF and FBS together as chemoattractants (rows 3-9), and to FBS alone as a chemoattractant in a dose curve (rows 10-12) compared to background (unstimulated) cell migration in the top rows. Concentrations of chemoattractants are indicated in the left columns. Notably, with the same amount of HGF present, adding increased amounts of FBS increased cell migration in 7 of the 8 cell lines (Figure 2A).

**Figure 2.**
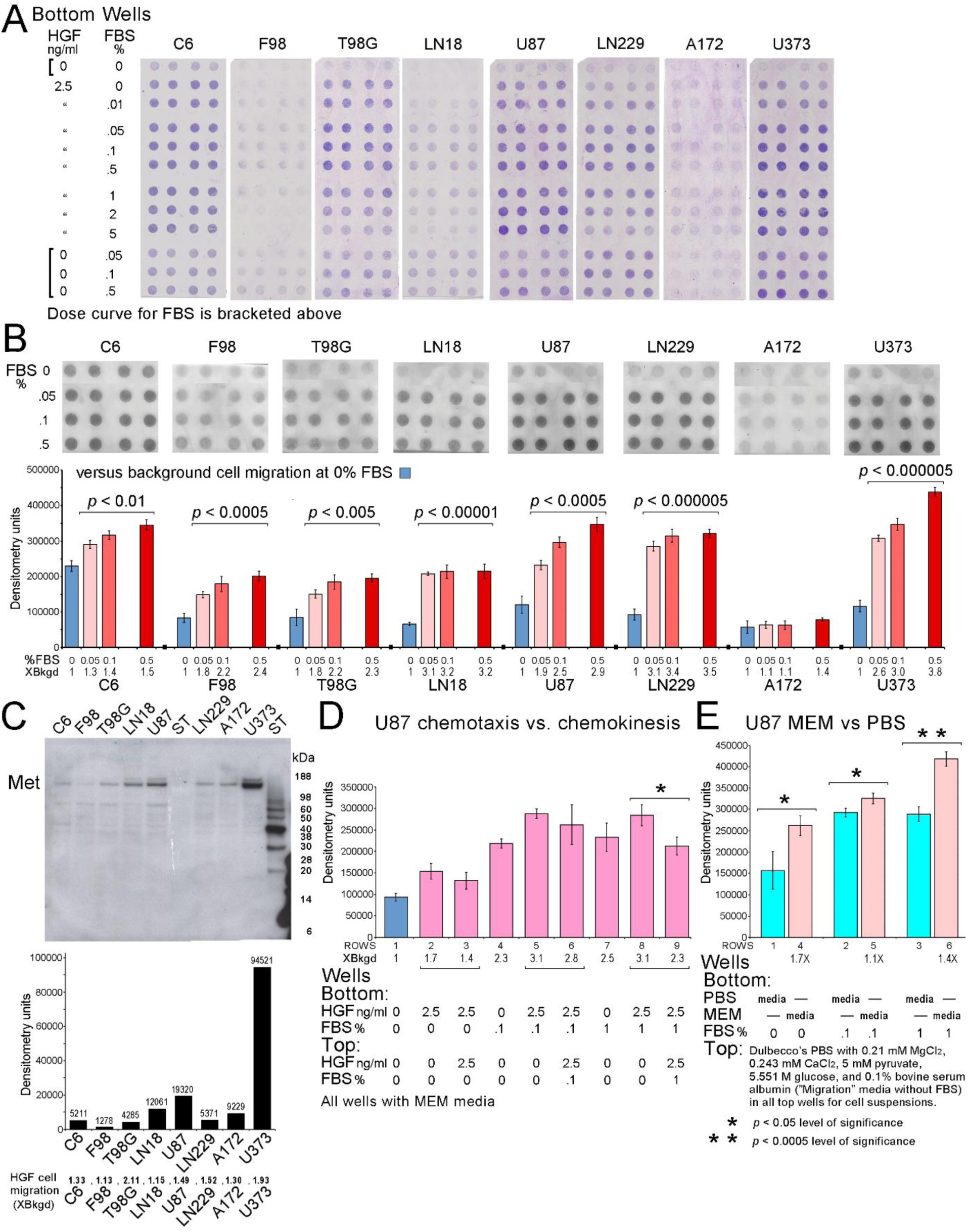
Migration of glioblastoma cell lines to hepatocyte growth factor (HGF) and fetal bovine serum (FBS). (A) Representative Boyden chamber assays of 2 rat astrocytoma (C6 and F90) and 6 glioblastoma cell lines (T98G, LN18, U87, LN229, A172, and U373) showed unstimulated (background) cell migration (row 1, at the top), cell migration to HGF (row 2), HGF and FBS (rows 3-9), and to FBS alone (rows 10-12) using concentrations in the bottom wells as indicated in the left columns. All cell lines, except for A172, migrated to FBS alone. With the same amount of HGF present, adding increasing amounts of FBS increased chemotactic cell migration in the other 7 cell lines. These images were obtained by using a transparency scanner. (B) Portions of scanned (regular scanner) assay images using FBS alone as the chemoattractant were digitized for densitometry (total pixels for units) with results shown as bar charts. All cell lines showed significant migration to FBS except for A172 which only had a trend, *p* = 0.0638, for 0.5% FBS. The entire collection (3 assays per cell line) is available in Appendix A for C6, F98, T98G, and LN18 (Figure A2) and U87, LN229, A172, and U373 (Figure A3). The first assays for C6, F98, LN18, and LN229, the second assay for T98G, and the third assays for U87, A172, and U373, were digitized for densitometry. (C) An immunoblot with a bar graph of densitometry results underneath shows the relative amounts of HGF’s receptor, Met (β subunit, 145 kDa) in each cell line. Also, HGF-stimulated cell migration is listed below the bar graph. HGF-stimulated cell migration (digitized densitometry of row 2 of the migration assays divided by background (no stimulation) migration for row 1 cell pellets in B) is stated in units of times(X) background cell migration. The HGF-stimulated cell migration compared with levels of Met lacks notable correlation. The gel for the immunoblot showing equalized protein loading is available in Appendix A (Figure A4A). (D) Unstimulated cell migration (blue bar representing the average of the assay replicates) in U87 cells is shown for contrast with the chemotactic and chemokinetic strength of HGF and FBS (pink bars representing averages of replicates in each row of the assays using HGF alone or HGF and FBS) that were both greater than unstimulated cell migration. The chemoattractant strength of HGF alone (bar for row 2) and with 0.1% and 1% FBS (bars for rows 5 and 8, respectively) shows impressive enhancement by FBS. Chemokinetic strength of HGF alone (bar for row 3), with 0.1% and 1% FBS (bars for rows 6 and 9, respectively) also showed impressive enhancement by FBS. The combination of HGF, 2.5 ng/ml, and 1% FBS for chemoattraction (bar for row 8) significantly, *p* = 0.012, outperformed their chemokinetic effects (bar for row 9) on cell migration. The scanned image of the assay is also available in Appendix A (Figure A4B). (E) An assay that evaluated MEM for U87 migration studies included gradients for K^+^ and Cl^-^. In the absence of FBS, Dulbecco’s PBS in the bottom wells provided K^+^ as 2.7 mM KCl + 1.8 mM KH_2_PO_4_ and Cl^-^ as 2.7 KCl + 137.9 mM NaCl. MEM in the lower wells provided K^+^ as 5.3 mM KCl, and Cl^-^ as 5.3 mM KCl + 1.8 mM CaCl_2_-2H_2_O. For all upper wells without FBS, the Migration media provided K^+^ as 2.7 mM KCl + 1.8 mM KH_2_PO_4_ and Cl^-^ as 2.7 KCl + 137.9 mM NaCl + 0.21 mM MgCl_2_ + 0.243 mM CaCl_2_. See text for total mM for each ion. When the bottom wells contained Dulbecco’s PBS there was no significant gradient created for K^+^ or Cl^-^. However, when MEM was in the bottom wells, positive K^+^ and negative Cl^-^ gradients were generated. Both K^+^ and Cl^-^ ions are used for volume changes in cell migration. The positive gradient for K^+^ is supported as being a chemoattractant in view of the significantly increased migration, *p* = 0.0144, in MEM versus Dulbecco’s PBS in the lower wells. None of the wells contained FBS for the above comparison. However, MEM in the lower wells was associated with significantly increased cell migration with 0.1 and 1% FBS also present, *p* = 0.0136 and *p* = 0.000442, respectively, compared to Dulbecco’s PBS in the lower wells having the same FBS concentrations. Averages of rows for the Densitometry units indicated by bars are shown with 95% confidence limits (B, D, and E). Variability in staining of the background migration, as shown in the top rows of representative assays (A) and assay-to-assay variation for each cell line in Appendix A (Figures A2 and A3), are routine findings. Each circle of stained migrated cells shown is 3 mm in diameter.

Portions of scanned assay images of FBS alone as the chemoattractant in a dose curve are shown as migrated cell pellets with densitometry results in bar charts below their images. All cell lines, except A172, showed significant migration to FBS alone. U87 and U373 results showed the most impressive stepwise increase in chemotactic migration at all 3 concentrations of FBS versus background migration with all *p* values < 0.0005 and 0.000005, respectively. U373 chemotaxis to 0.5% FBS, 3.8 times the background (unstimulated) migration, was the highest level for all cell lines. A172 cells showed a trend for increased migration to 0.5% FBS, *p* = 0.0638. Some of the cell lines showed flattening of their responses as levels of FBS increased (Figure 2B). All assays (3 per cell line) are shown in Appendix A for C6, F98, T98G, and LN18 (Figure A2) and U87, LN229, A172, and U373 (Figure A3). Cell migration specifically for HGF in each cell line is stated as a multiple (X) of the background cell migration, obtained by dividing the 2^nd^ row of assays’ cell pellets densitometry results by the corresponding 1^st^ row of background cell migration, bottom row (Figure 2C). The immunoblot shows reactivity for HGF’s receptor, Met (β subunit, 145 kDa). The Met reactivity with U87 and LN229 has been published previously [134] but here the relative amounts of Met in 8 cell lines are reported, including rat (C6 and F98) and human glioblastomas (T98G, LN18, U87, LN229, A172, and U373). As shown in the bar chart there was a lack of correlation with the cell motility results shown below it (Figure 2C). In Appendix A, the gel image showing equalized protein loading for all 8 cell lines in the immunoblot is available (Figure A4A). The order for HGF cell migration from highest to lowest results for all 3 assays of each cell line was T98G (2.11X), U373 (1.93X), LN229 (1.52X), U87 (1.49X), C6 (1.33X), A172 (1.30X), LN18 (1.15X), and F98 (1.13X). Twelve of the 92 data points of visibly migrated HGF cell pellets had densitometry results showing slightly less than 1X background cell migration and were rounded up to 1X for these results. A few replicates did not show visible unstimulated or stimulated cell pellets.

Earlier studies had noted chemokinesis with HGF and 0.1% FBS in U87 cell migration assays [136] and also chemokinesis with 5% FBS alone in LN229 cell invasion assays [135]. The dose curve for FBS includes additional data points as shown in Figure 2D. A dose curve was also published earlier with fewer data points for chemoattraction using HGF, FBS, and both for U87 and LN229 cells previously (134) but the assays here include additional data points and 6 more cell lines. The chemoattractant strength of HGF alone (row 2), and with 0.1% and 1% FBS (rows 5 and 8, respectively) was enhanced by both FBS concentrations in the U87 cells. Chemokinetic strength of HGF alone (row 3), with 0.1% and 1% FBS (rows 6 and 9, respectively) was also enhanced by FBS. Chemotaxis and chemokinesis at 0, 0.1, and 1% FBS demonstrated strong enhancement of FBS versus HGF alone for either type of U87 cell migration. The combination of HGF, 2.5 ng/ml, and 1% FBS for chemoattraction significantly, *p*= 0.012, outperformed their chemokinetic effects on cell migration (Figure 2D). A scanned image of the assay is also available in Appendix A (Figure A4B). A portion of the immunoblot for Met in U87 and LN229 cells with the gel was published previously (134) but without the entire immunoblot for U87 and LN229 cells plus 6 additional cell lines and all standards as included in this study

Another assay to evaluate MEM for migration studies included gradients of K^+^ and Cl^-^ in migration assays of U87 cells. Both ions are redistributed with volume changes during cell migration. In the absence of FBS, Dulbecco’s PBS in the bottom wells provided K^+^ as 2.7 mM KCl + 1.8 mM KH_2_PO_4_ (**4.5 mM K^+^**) and Cl^-^ as 2.7 KCl + 137.9 mM NaCl (**140.6 mM Cl^-^**). MEM in the lower wells provided K^+^ as 5.3 mM KCl (**5.3 mM K^+^**) and Cl^-^ as 5.3 mM KCl + 1.8 mM CaCl_2_-2H_2_O (**8.9 mM Cl**^-^). See Methods for a minor issue with KCl molarity in MEM. For all upper wells without FBS, the Migration media provided K^+^ as 2.7 mM KCl + 1.8 mM KH_2_PO_4_ (**4.5 mM K^+^**) and Cl^-^ as 2.7 KCl + 137.9 mM NaCl + 0.21 mM MgCl_2_ + 0.243 mM CaCl_2_ (**141.5 mM Cl^-^**). Potassium did not generate any gradient for cells migrating from upper to lower wells when Dulbecco’s PBS was used in the bottom wells and there was almost no gradient for chloride either. However, when MEM was used in the lower wells a reverse gradient for Cl^-^ was generated with more Cl^-^ in the upper versus the lower wells. With MEM in the lower wells there was a positive gradient to potassium for migrating cells (5.3 mM in the lower wells versus 4.5 mM in the upper wells). Chloride ions present in a negative gradient are probably not responsible for the 1.7X increase in cell migration when MEM was used in the bottom wells. However, the positive gradient for K^+^ may have contributed to the significantly increased migration, *p* = 0.0144, without FBS present. MEM as the media in the lower wells was also associated with significantly increased cell migration when 0.1 and 1% FBS were present, *p* = 0.0136 and *p* = 0.000442, respectively, compared to using Dulbecco’s PBS for the media in the lower wells having the same FBS concentrations (Figure 2E). MEM contains 5.6 mM dextrose, 1 mM pyruvate, and 2 mM l-glutamine, whereas Migration media contained 5.6 M glucose and 5 mM pyruvate. Therefore, MEM as a source of fuel in the bottom wells was probably not a significant differentiating factor in the results. MEM also contains 17.856 mM sodium bicarbonate, 0.8 mM magnesium sulfate (more Mg^2+^ than Migration media (0.21 mM MgCl_2_), less sodium than Dulbecco’s PBS and Migration media, amino acids, vitamins, etc. A scanned image of the assay is available in Appendix A (Figure A4C). These results support the role of K^+^ in enhancing cancer cell migration. Variable background migration, as shown in the top rows of representative assays (Figures 2A-B) and assay-to-assay variation for each cell line as shown in Appendix A files (Figures A2 and A3) are routine for cell migration assays.

### 3.3. KCN Genes Differentially Expressed in 70 Studies

#### 3.3.1. General Features of Detected Differentially Expressed KCN Genes

The availability of K^+^ for cancer cell survival and migration suggested by experimental studies just described led to consideration of the large KCN family of genes being altered in cancer cells. Published genomic landscapes, i.e. heat maps (46 studies), Venn diagrams, volcano plots, tables, etc. were found in 70 cancer studies including KCN (potassium channel) family members as one or more of 53 KCN genes that were differentially expressed genes (DEG) along with numerous co-expressed non-KCN DEG. Selection criteria for DEG (mostly *p* < 0.05 differences, chance of false discovery, etc.) were set by the authors who incidentally detected and reported the KCN DEG among groups of DEG Appendix B (Table B1, sample view, and the full list, AllE1E70DEG3239Total, is available after the References and Disclaimer in a table format). Parameters were applied in each study for a particular type of malignancy, without or with perturbations, such as tumor treatment, transcription factor, over-expression of an oncogene(s), or according to survival, disease progression, primary versus metastasis, recurrence, etc. The 53 KCN DEG are distributed across 17 chromosomes (1-3, 5-12, 17-21, and X). Although many of these KCN genes encode membrane pore subunits with K^+^ selectivity, some provide modulatory/regulatory effects on pore function when present with other subunits that may also function independently as pores. Heterodimers among KCN family members are common. Members of 4 KCN subfamilies, Kv (voltage dependent, related, *KCNA*, *KCNQ*, *KCNH*, etc.), Kir (inwardly rectifying, *KCNJ*), K2P (tandem two pore domain, *KCNK)*, and KCa (calcium dependent, related, *KCNM*, *KCNN*, etc.) were found in 32(46%), 10(14%), 14(20%), and 29(41%), respectively, of the 70 studies.

#### 3.3.2. Proton(H^+^) Sensitivity of Detected KCN DEG

In keeping with the connection proposed for K^+^ and H^+^, the H^+^-sensitivity of KCN DEG encoded proteins was relevant. At least 33 (62.3%) of the 53 KCN DEG encode gene products that are H^+^-sensitive according to 135 publications listed in Appendix B (Table B2). Fifty-two (74.79%) of the 70 studies contained at least one KCN DEG identified as being H^+^- sensitive. One, 2, 3, and 6 H^+^-sensitive KCN DEG genes were found in 36, 14, 1, and 1 of the studies, respectively. H^+^- sensitive genes in each of the 4 subfamilies, Kv, Kir, K2P, and KCa were found as DEG in 17 (24%), 9 (13%), 13 (19%), and 21(30%) of 70 studies, respectively. The most frequently detected DEG, *KCNN4* and *KCNMA1*, present in 9 and 6 studies, respectively, encode H^+^-sensitive proteins (red H^+^ above columns) and are also Ca^2+^-sensitive (Figure 3A).

**Figure 3.**
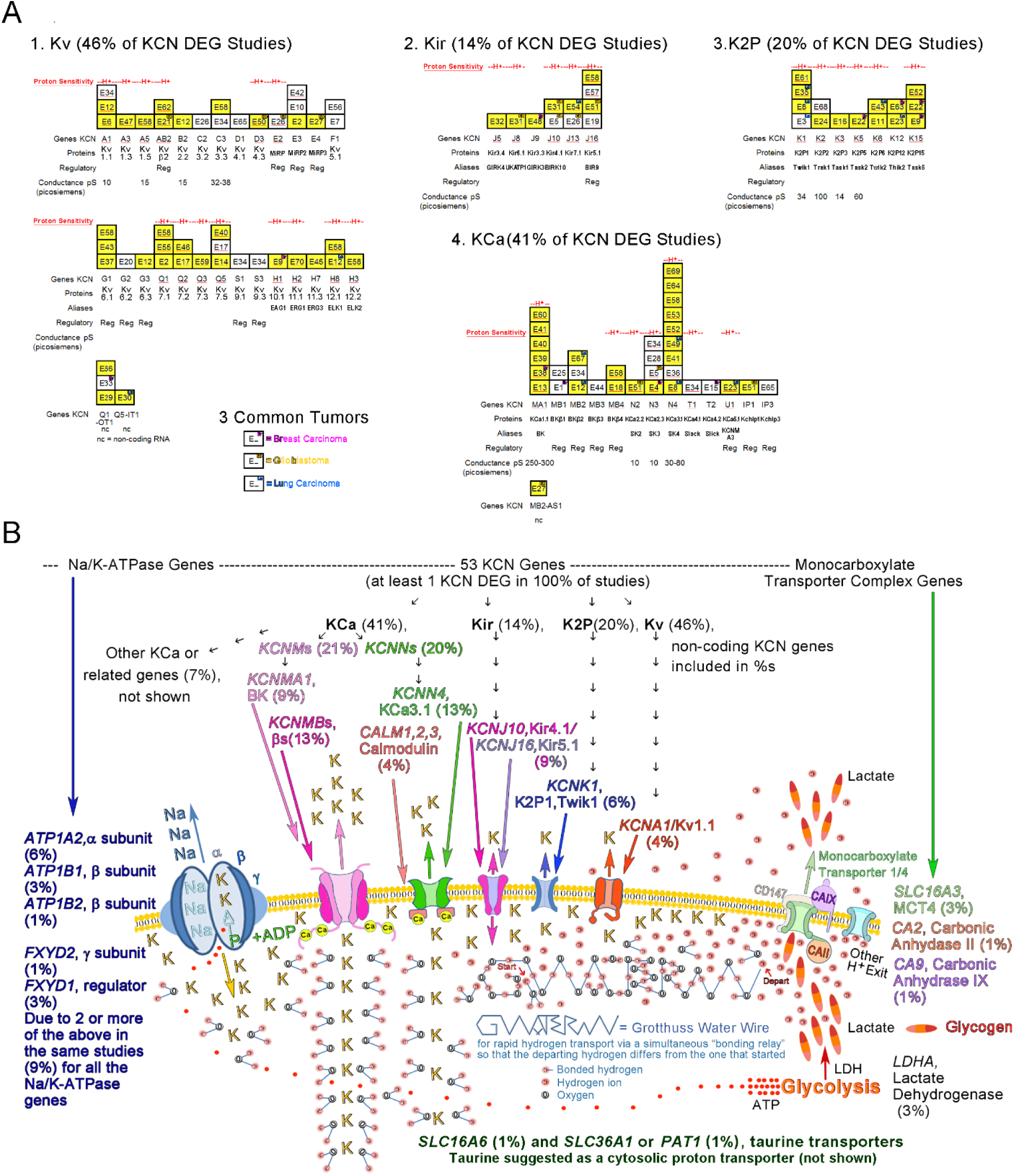
The KCN DEG and related non-KCN DEG detected in E1-E70 studies with relevant information are shown. (**A**) 53 KCN DEG of 4 KCN subfamilies detected are listed along the x-axes in the specific studies that are listed vertically in boxes above the specific KCN gene names, reflecting their frequency in 70 studies. The 3 most frequent types of tumors (breast, glioblastoma, and lung) are highlighted with color-coded superscripts, as listed under the first group of histograms. The 33 (62.26%) of KCN DEG that encode H^+^-sensitive proteins are labeled with red H^+^ above each column. Studies with KCNs detected that encode H^+^-sensitive proteins also have their names underlined in red within the boxes of the histograms. 52(74.29%) of the 70 studies contain at least one KCN encoding a H^+^-sensitive protein. Yellow shading of the boxes labeled for specific studies indicates that these studies were included in 1 or more of 8 categories shown later (Figure 4A). The non-coding KCNs are in separate small histograms included with their appropriate KCN subfamily group. If only studies that included at least one protein-coding KCN DEG are considered, then 52(78.79%) of those 66 studies contained at least one KCN encoding a H^+^-sensitive protein. Multiple KCN DEG were detected in 18 (27.27%) of 66 studies. (**B**) A stochastic view of K^+^-related dynamics in cancer cells is presented based on collective incidences of both KCN and related non-KCN DEG identified in E1-E70. Images of the most frequent members of each of the 4 KCN subfamilies along with pertinent non-KCN membrane proteins and their complexes are consistent with published graphical representations [141–148]. There were relatively few studies in E1-E70 with differential expressions detected for genes encoding Na/K-ATPases (9% for 5 genes), the monocarboxylase transporter complex (5% for 3 genes), lactate dehydrogenases (3% for 1 gene), taurine transporters (2% for 2 genes), and 0% for related functional analogues not listed. Therefore, abnormal K^+^ flux allowing K^+^/H^+^ exchange, if present in these same cancer studies with KCN DEG present, suggests KCN DEG for playing a role in the pH changes following rapid cytosolic H^+^ transport (putatively Grotthuss type) to exit sites that include release of protons with lactate. Also, the lack of DEG for glycogen synthases assumes only normal (if any) intracellular storage of lactate as glycogen. Abnormal accumulations of K^+^ at inner membrane surfaces may have occurred due to KCN DEG (in assorted combinations) to exchange with bound H^+^, either due to genetic loss or down regulation of any of the numerous KCN membrane proteins mediating outward flux or a gain via KCNs mediating increased inward flux. The movement of K^+^-7H_2_O (represented by 2 H_2_O per K^+^ shown in a “cross sectional” view) within the cells may bring water to outer cytoplasmic regions, possibly enhancing Grotthuss conditions near inner membranes in cancer cells. The putative K^+^-related changes in cancer cells that at the same time maintain relatively normal expression levels of pertinent non-KCN genes, as seen in the majority of the 70 KCN DEG studies, would be consistent with the complex scenario shown (**B**). E1-E70 citations are listed in *Section 3.3.6. Categories Among Studies with E1-E70 Citations Per Category*.

#### 3.3.3. Coding and Multiplicity of KCN DEG and Non-KCN DEG

Four of the 70 studies identified only non-coding KCN genes, *KCNQ1-OT1* (E29, E33, and E66) and *KCNQ5-IT1* (E30), as DEG. Study E27 identified *KCNMB2-AS1* as well as *KCNE4* as DEG. Disregarding the 4 studies with only non-coding KCNs detected as DEG, multiple (2 - 9) KCN DEG were found in 18 (27.3%) of 66 studies with protein-coding KCN DEG. Thirty-eight (71.70%) of the 53 KCN DEG were detected with other coding KCN DEG in at least one of the 66 studies. There were 102 instances of 53 KCN DEG in a total of 3,239 DEG instances (2,490 non-KCN DEG without repeats). The average numbers of KCN DEG and all DEG per study were 1.53 +/- 0.33 (95% CI) and 45.91+/- 7.92 (95% CI), respectively. The correlation coefficient for number of KCN DEG and size of all DEG (KCN and non-KCN) groups was 0.379. If only the studies that included at least one protein coding KCN DEG are considered, then 52 (78.79%) of the 66 studies detected at least one H^+^-sensitive KCN DEG (Figure 3A).

#### 3.3.4. Malignancies in E1-E70

The 3 most frequently found types of malignancies in the 70 studies were lung cancer, breast cancer, and glioblastoma with detected genes encoding H^+^- sensitive products **bolded** in the following list. The <u>lung cance</u>r studies with their KCNs include: E3(***KCNK1***), E8(***KCNK1***, ***KCNN4***), E12(***KCNH8***, *KCNMB2*), E23(***KCNU1***), E30(*KCNQ5-IT1*), E35(***KCNK1***), E43(***KCNK6***), E49(***KCNN4***), 2), E54(***KCNJ13***), E63(*KCNK12*), and E67(*KCNMB2*). The <u>breast cance</u>r studies include: E1(*KCNMB1*), E4(***KCNN3***), E9(***KCNH1***, ***KCNK15***), E15(*KCNT2*), E22(***KCNK5***, ***KCNK15***), E33(*KCNQ1-OT1*), E38(***KCNMA1***), E48(*KCNJ9*), and E63(*KCNK12*). The <u>glioblastoma</u> studies include: E5(***KCNJ10***, ***KCNN3***), E21(***KCNAB2***), E26(***KCNE2***, ***KCNJ13***), E27(KCNE4, KCNMB2-AS1), E31(***KCNJ10***), E50(***KCND3***), E51(KCNIP1, ***KCNJ16***, ***KCNN2***). H^+^-sensitive KCNs were found in 86%, 73%, and 44% of the 7 glioblastoma, 11 lung cancer, and 9 breast cancer studies, respectively. All three of the melanoma studies detected H^+^- sensitive KCN DEG. Numerous types of malignancy listed in Appendix B, including NCI-60 cell lines (E16), detected H^+^-sensitive KCN DEG, see Appendix B (Figure B1A). Common informative protein abbreviations in the KCN family are also listed (Figure B1B). E1-E70 citations for E1 – E70 are listed in *Section 3.3.6. Categories Among Studies with E1-E70 Citations Per Category*

#### 3.3.5. Stochastic View of K^+^-Related Dynamics in E1-E70

A percentage based stochastic view of K^+^-related gene expression dynamics in cancer cells is presented based on collective incidences of both KCN and related non-KCN DEG identified by the authors of each paper. Images of the most frequent members of each of the 4 KCN subfamilies along with pertinent non-KCN membrane proteins and their complexes are consistent with published graphical representations [141–148]. In view of the relatively small number of 70 studies with differential expressions detected for genes shown in Figure 3B that encode Na/K-ATPase (9% for 5 genes), the monocarboxylase transporter complex (5% for 3 genes), lactate dehydrogenases (3% for 1 gene), taurine transporters (2% for 2 genes), and 0% for genes encoding functional analogue proteins not listed, the abnormal K^+^ flux in these same cancers with KCN DEG can be considered for supplying K^+^ for a K^+^/H^+^ exchange that initiates pH changes. Rapid cytosolic H^+^ transport (putatively involving Grotthuss water wires) to exit sites for proton release with lactate is included to complete the pH reversal scenario. The pertinent lack of DEG in the E1-E70 studies for glycogen synthases, to mediate anomalous storage of lactate without extrusion, is included because glycogen storage of lactate occurs in humans, mice, bacteria, etc. [149–152]. Abnormal accumulations of K^+^ at inner membrane surfaces are illustrated since they are proposed to promote K^+^/ H^+^ exchange that releases bound H^+^. Accumulations of K^+^ occur either due to genetic loss or down regulation of KCN membrane proteins mediating outward flux or increased inward flux due to a gain or loss in copy number and/or expression of the KCN genes found to be differentially expressed in E1-E70. The movement of K^+^-7H_2_O (indicated with 2 H_2_O per K^+^, as a “cross-sectional” view in Figure 3B within cells may bring water to outer cytoplasmic regions, possibly promoting Grotthuss conditions in cancer cells near inner membranes. These putative K^+^-related changes in cells that maintain relatively normal expression levels of pertinent non-KCN genes as seen in the majority of the KCN DEG studies suggest the scenario shown (Figure 3B).

#### 3.3.6. Categories Among Studies with E1-E70 Citations Per Category

Eight categories describe 50 of the 70 studies (Figure 4A). These were author-designated which also had relevant DEG in at least one of the E1-E70 studies. Some studies belonged in more than one category and 20 studies were in no category as designated last item in the list below.

**Figure 4.**
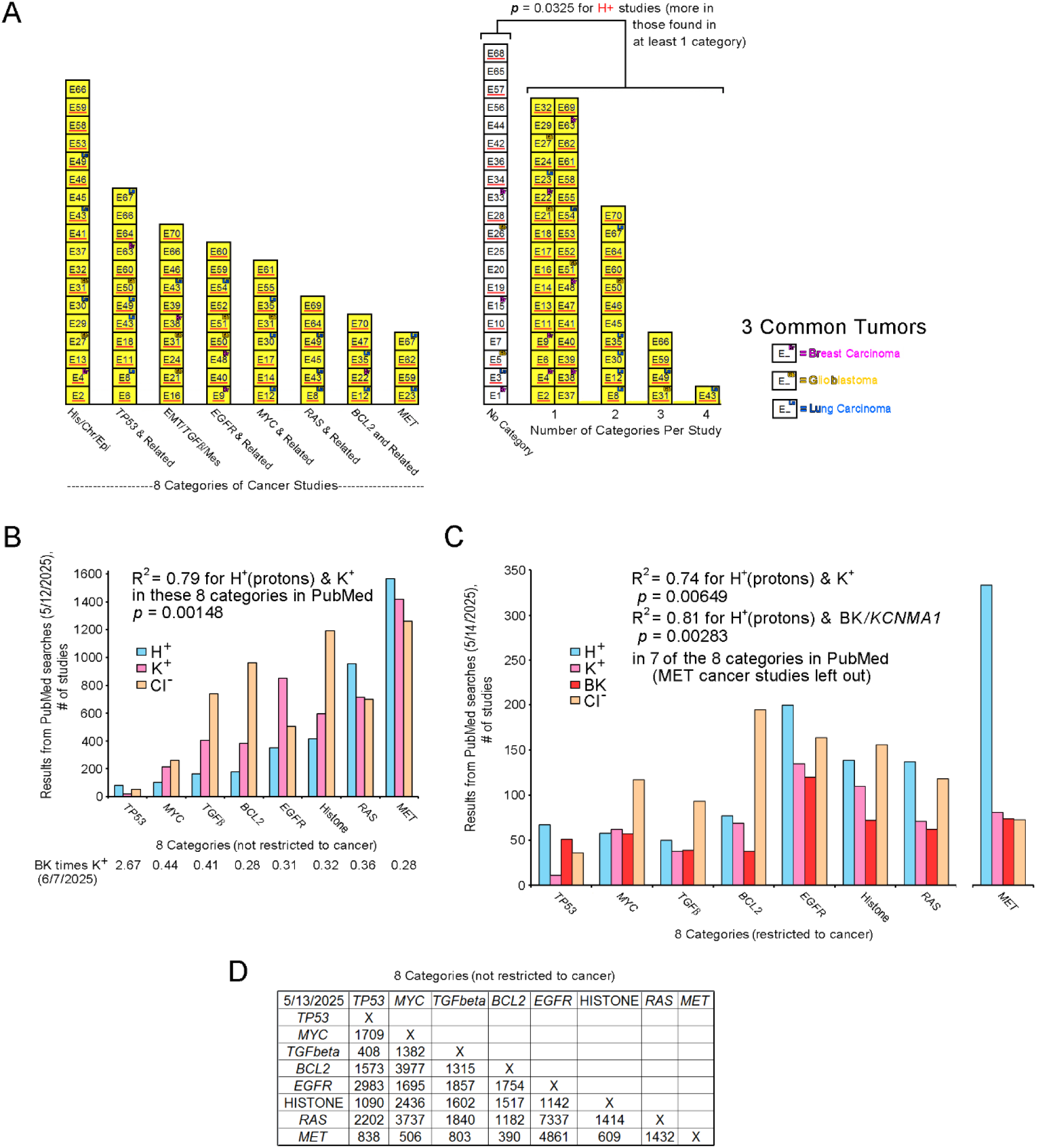
Oncogenes and DNA modifiers may play a role in cancer studies detecting KCN DEG encoding H^+^-sensitive proteins. (**A**) The KCN DEG that encode H^+^-sensitive proteins are highlighted with red underlining of the study names. Histograms of 8 categories (x-axis) of cancer studies among E1-E70 that contain KCN DEG are shown in the left histogram. The right histogram also includes 20 studies in “no category” (see text) in the first column. Comparison of the studies in “no category” versus those included in at least one of eight categories shown in the left histogram indicated a significant difference for the number of studies detecting KCN DEG encoding H^+^-sensitive proteins. With a Fisher’s Exact test studies that detected one or more KCN DEG that encoded a H^+^-sensitive protein showed a significant difference in the numbers of studies if the studies were in at least one of the eight categories versus no category, *p* = 0.0325. (**B – D)** The PubMed database (more than 38 million citations) was checked with word searches for the incidence of oncogene and histone publications related to H^+^, K^+^, BK (encoded by *KCNMA1*), and Cl^-^ (contrasting ion) to detect correlations related to findings in the 70 studies selected by detection of KCN DEG. (**B**) On May 12, 2025, searches for each category of “TP53”, “MYC”, “TGFbeta”, “BCL2”, “EGFR”, “histone”, “RAS”, and “MET” were performed separately for “proton”, “potassium”, and “chloride”. For all 8 categories, the total numbers of citations for “proton”, “potassium”, and “chloride” were 1565, 1418, and 1261, respectively, and the Pearson’s product-moment correlations for “proton” with “potassium” and “proton” with “chloride” were R = 0.891 (*p* = 0.00148) and 0.599 (*p* = 0.0584), respectively. The Pearson’s product moment correlation for “potassium” and “chloride” was 0.689 (*p* = 0.0295). R^2^ for “proton” with “potassium” was 0.79. “BK” and “potassium” searches on 6/7/2025 revealed “BK” as a potential large component of the “potassium” citations, approximately one third for 7 of the categories. (**C**) When the same searches also included “cancer” as a third word, the numbers of citations for “proton”, “potassium”, “BK”, and “chloride” on May 14, 2025, were 1061, 577, 513 (88.9% of “potassium” search results), and 952, respectively. However, when MET was left out for better “proton” correlations, the numbers of citations for “proton”, “potassium”, “BK”, and “chloride” on May 14, 2025, were 728, 496, 439 (88.5% of “potassium” search results), and 879, respectively, and the Pearson’s product-moment correlations for “proton” with “potassium” and “chloride” were R = 0.860 (*p* = 0.00649) and 0.462 (*p* = 0.148), respectively, across the 7 categories. Pearson’s product-moment correlation for “proton” with “BK” was R = 0.901 (*p* = 0.00283) across the 7 categories. R^2^ for “proton” with “BK” was 0.81. R^2^ for “proton” with “potassium” was 0.74. The results for correlation between numbers of “Met” citations and those for the ions differed in results from the “cancer” searches in (C) compared to search results not restricted to “cancer” shown in (B). (**D**) The numbers of citations shared among the 8 categories of 7 oncogenes and histone are shown. E1-E70 citations are listed in *Section 3.3.6. Categories Among Studies with E1-E70 Citations Per Category*

1. Of the 18 studies in the <u>first category</u> of histone/chromatin/epigenetic (His/Chr/Epi or histone/DNA), 10 were in this category due to specific genes described by the authors of the studies, including: E2(*N6AMT1*, previously *KMT9*), E13(*MSH2*, *BET*), E32(*H3K27me3*, *EZH2*), E41(*HDAC8*), E46(*KDM2B*), E49(*KMT2D*), and E53(*H3K27*). Also, E27’s and E29’s authors focused on chromatin access and imprinting, respectively. Two imprinted genes, *KCNQOT1* and *PAX8*, were DEG in E29. Twenty-one imprinted DEG (*AXL*, *DLK1*, *GDAP1L1*, *GNG7*, *GRB10*, *IGF2*, *KCNQ1*, *KCNQOT1*, *LRRTM1*, *MEG3*, *NDN*, *NNAT*, *NTM*, *PAX8*, *PEG3*, *PEG10*, *PHLDA2*, *PLAGL1*, *RHOBTB3*, *RTL1*, and *TFPI2*) were detected among a total of 2560 DEG for a rate of 0.82% in E1 - E70. When repeats are included, there were 32 instances of imprinted genes among 3,242 instances of DEG in E1-E70 for a rate of 0.99%. Detection of 0.82% or 0.99% imprinted genes is expected from the genome (approximately 200, i.e. less than1% of all genes) according to Tucci and colleagues who also provided a list of human imprinted genes [153]. Therefore, the presence of imprinted genes was not used as an inclusion criterion for the His/Chr/Epi category. In addition to the 10 studies per author designations, 8 additional studies detected His/Chr/Epi related DEG as follows: E4(*DNMT3B*), E30(*KDMA-AS1*), E31(*H2A.X*, previously *H2AFX*), E37(*H2AC18*), E43(*H2AC18*, *H2BC5*, *H2BC12*), E45(*NUPR1*), E58(*DNMT3B*), and E66(*BDR4*, *H1-4*(previously *HIST1HIE*)). The entire His/Chr/Epi category includes the following studies with their citations: E2 [154], E4 [155], E13 [156], E27 [157], E29 [158], E30 [159], E31[160], E32 [161], E37 [162], E41 [163], E43 [164], E45 [165], E46 [166], E49 [167], E53 [168], E58 [169], E59 [170], and E66 [171] (Figure 4A, left side).

2. In the <u>second category</u> there were 9 studies on *TP53* according to the authors’ focus in E6, E8, E11, E49, E49, E60, E63, E64, and E67. Additionally, 3 studies found *TP53* and related genes as DEG as follows: E18(*MDM2*), E43(*TP53INP1A*), and E50(*TP53*). The 12 studies in this category are as follows: E6 [172], E8 [173], E11 [174], E18 [175], E43 [164], E49 [167], E50 [176], E60 [177], E63 [178], E64 [179], E66 [171], and E67 [180] (Figure 4A, left side).

3. In the <u>third category</u> there were 7 studies on Epithelial Mesenchymal Transition (EMT)/ *TGFb*/ Mesenchymal Tumor (EMT/*TGFb*/Mes) according to the authors’ focus with specific genes, *CDH1*, *2*, *VIM*, *TGFb1*, *2*, *3*, in some of these studies as follows: E16(*CDH1*,*2*, *TGFb1*, *VIM*), E24(*TGFb1*), E31, E38, E39, E43(*CDH2*), and E46. Three additional studies also found some of these genes as follows: E21(*TGFb1*), E66(*VIM*), and E70(*CDH2*, *TGFb2*, *TGFb3*). The 10 studies this category are as follows: E16 [181], E21 [182], E24 [183], E31 [160], E38 [184], E39 [185], E43 [164], E46 [166], E66 [171], and E70 [186] (Figure 4A, left side).

4. In the <u>fourth</u> <u>category</u> there were 5 studies on *EGFR* and related genes according to the authors with corresponding DEG in two of their studies as listed, E9, E48, E50(*EGFR*, *ERBB2*), E51(*EGFR*), and E52. Additionally, 4 studies found related DEG as follows: E40(*NRG1*), E54(*NRG2*), E59(*NRG1*), and E60(*ERBB4*). The 9 studies in this category are as follows: E9 [187], E40 [188], E48 [189], E50 [176], E51 [190], E52 [191], E54 [192], E59 [170], and E60 [177] (Figure 4A, left side).

5. In the <u>fifth category</u> there were 5 studies on *MYC* according to the authors with *MYC* as a DEG in 2 of the studies as seen in the following list: E14(*MYC*), E30, E31, E55, and E61(*MYC*). Additionally, 3 studies had *MYC* or one of its related genes as seen here: E12(*MYCL*), E17(*MYCN*), and E35(*MYC*). The 8 studies this category are as follows: E12 [193], E14 [194], E17 [195], E30 [159], E31 [160], E35 [196], E55 [197], and E61 [198] (Figure 4A, left side).

6. The <u>sixth category</u> is comprised of 6 studies on *RAS* per the authors’ interest, with *KRAS* found as a DEG in 2 studies as follows: E8, E43, E45, E49, E64(*KRAS*), and E69(*KRAS*). The 6 studies this category are as follows: E8 [173], E43 [164], E45 [165], E49 [167], E64 [179], E69 [199] (Figure 4A, left side).

7. In the <u>seventh category</u> the authors focused on *BCL2* in E47 but did not list it as a DEG. However, 4 other studies identified *BCL2* as a DEG and 1 also identified *BCL2L1* as well, as seen in the following list: E12(*BCL2*), E22(*BCL2*), E35(*BCL2*), and E70(*BCL2*, *BCL2L1*). The 5 studies in this category are as follows: E12 [193], E22 [200], E35 [196], E47 [201], E70 [186] (Figure 4A, left side).

8. In the <u>eighth category</u> the authors focused on *MET* in 1 study and found it as a DEG, E23(*MET*). Additionally, 3 studies also identified *MET* as a DEG, E59(*MET*), E62(*MET*), and E67(*MET*). The 4 studies in this category are as follows: E23 [202], E59 [170], E62 [203], E67 [180] (Figure 4A, left side). All studies in the 8 categories are shaded yellow in Figure 4.

9. No category. Twenty studies were not in any of the above 8 categories and are listed as follows: E1 [204], E3 [205], E5 [206], E7 [207], E10 [208], E15 [209], E19 [210], E20 [211], E25 [212], E26 [213], E28 [214], E33 [215], E34 [216], E36 [217], E42 [218], E44 [219], E56 [220], E57 [221], E65 [222], and E68 [223] (Figure 4A, right histogram, no yellow shading).

#### 3.3.7. H^+^-Sensitivity Significant in E1-E70 Studies Within 8 Categories

A histogram (Figure 4, right side) shows the studies collectively in the 8 categories versus those not in any category, with the studies that detected H^+^-sensitive KCN DEG underlined with red. The difference for detection of H^+^-sensitive KCN DEG was significant between the two groups, *p* = 0.0325, Fisher’s Exact Test for count data. Eleven studies were in 2 categories (E8, E12, E30, E35, E45, E46, E50, E60, E64, E67, and E70). Four studies were in 3 categories (E31, E49, E59, and E66), and E43 was included in 4 categories. All 8 categories were among those that were shared in studies (Figure 4A, right histogram, yellow shading). E1-E70 citations are listed in *Section 3.3.6. Categories Among Studies with E1-E70 Citations Per Category*.

#### 3.3.8. PubMed Citations for “Proton”, “Potassium”, etc. in Categories

The PubMed database was checked with word searches for the incidence of 7 oncogenes and histone publications being related to those for H^+^, K^+^, BK (encoded by *KCNMA1*), and Cl^-^(a potential contrasting ion), without and with “cancer”, to compare with the 70 studies selected based on detection of KCN DEG. On May 12, 2025, searches for each category of TP53, MYC, TGFbeta, BCL2, EGFR, histone, RAS, and MET were performed separately for “proton”, “potassium”, and “chloride” and “cancer” was not included. For all 8 categories, the total numbers of citations for “proton”, “potassium” and “chloride” were 1565, 1418, and 1261, respectively, and Pearson’s product moment correlations for “proton” with “potassium” and “chloride” were R = 0.891 (*p* = 0.00148) and 0.599 (*p* = 0.0584), respectively. Pearson’s product-moment correlation for “potassium” with “chloride” was 0.689 (*p* = 0.0295). R^2^ for proton with potassium was 0.79. Later searches on 6/7/2025 for “BK” and “potassium” revealed “BK” citations were probably a large component of those for “potassium”, possibly about one third in 7 categories (TP53 was much greater than one third) and are listed also below the x axis (Figure 4B). When the searches also included “cancer” as a third word, the numbers of citations for “proton”, “potassium”, “BK”, and “chloride” on May 14, 2025, were 1061, 577, 513, and 952, respectively. However, when MET was left out, the numbers of citations for “proton”, “potassium”, “BK”, and “chloride” on May 14, 2025, were 728, 496, 439, and 879, respectively, and Pearson’s product-moment correlations for “proton” with “potassium” and “chloride” were improved, R = 0.860 (*p* = 0.00649) and 0.462 (*p* = 0.4623), respectively, using only 7 categories. Analysis prior to MET removal is not shown. R^2^ for “proton” with “potassium” for 7 categories after MET removal was 0.74. Pearson’s product-moment correlation across the 7 categories for “proton” with “BK” was R = 0.901 (*p* = 0.00283). R^2^ for correlation of “proton” with “BK” across the 7 categories was 0.81. It is unknown why citations in the MET category no longer positively participated in the correlation seen in previous searches without “cancer” for “proton” and “potassium” but may have been due to the tendency for negative studies to remain unpublished. The answer could be important but is beyond the scope of this study. It also was noted that compared to searches without “cancer” shown in Figure 4B, the “BK” citations appeared to constitute an even larger proportion of “potassium” searches when “cancer” was included, while assuming that “potassium” in the searches included all or most of “BK” citations (Figure 4C). Numerous citations were shared pairwise in 2-word searches (not restricted to cancer) among the 8 categories of 7 oncogenes and histone with the numbers shown (Figure 4D). Thus, many studies involving the 8 categories describe them collectively. The E1-E70 studies provide a broad view of KCN genes among potassium studies. Citation searches of several KCN proteins other than BK did not produce sufficient numbers for comparisons.

#### 3.3.9. Non-KCN DEG Repeats in E1-E70

The non-KCN DEG found repeatedly in up to 5-7 studies were often detected with *KCNN4* and some with *KCNMA1*. Among the non-KCN repeat DEG, *ITGA2/*Integrin Alpha-2 was detected most frequently (7 studies). Six of these also included H^+^-sensitive KCN DEGs which are bolded as follows: **E8** (***KCNK1*, *-N4***), E15, **E39** (***KCNMA1***), **E41**(***KCNMA1***, ***-N4***), **E49**(***KCNN4***), **E64**(***KCNN4***), and **E69**(***KCNN4***). *KCNN4* was the most frequently found DEG, 9 times total and in 5 of 7 studies with *ITGA2* as a DEG. The four studies of co-expressed *KCNN4*, *ITGA2*, and *PLAUR*, E8 [173], E41 [163], E49 [167], and E69 [199], included 2 lung cancers, 1 melanoma, and 1 pancreatic cancer, in the categories of *His/Chr/Epi*, *TP53*, and *RAS* as listed (Figure 5A, left Venn diagram). *HMGA2/*High Mobility Group AT-hook 2, *LIF/*Leukemia Inhibitory Factor, and *PLAUR/*Plasminogen Activator, Urokinase Receptor (UPAR) were identified as non-KCN DEG in 6 studies each that also often detected *KCNN4* as a DEG.

**Figure 5.**
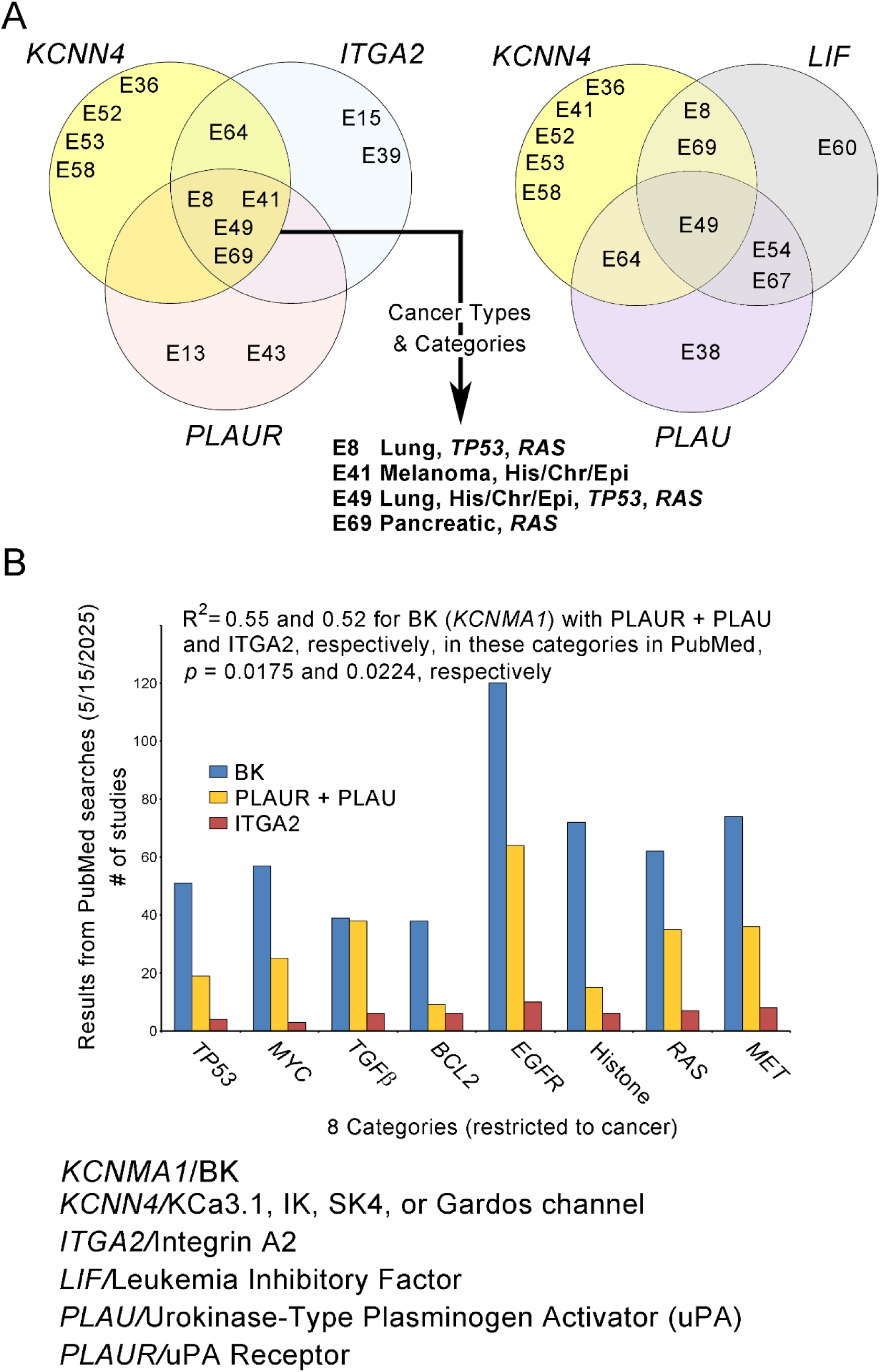
Repeat non-KCN DEG were found, often with *KCNN4* and some with *KCNMA1*. (A) Among the non-KCN repeat DEG, *ITGA2/*Integrin Alpha-2 was detected most frequently (7 studies). Six of these studies also included H^+^-sensitive KCN DEGs which are bolded as follows: **E8** (***KCNK1*, *-N4***), E15, **E39** (***KCNMA1***), **E41**(***KCNMA1***, ***-N4***), **E49**(***KCNN4***), **E64**(***KCNN4***), and **E69**(***KCNN4***). *KCNN4* was the most frequently found DEG, 9 times total and in 5 of 7 studies for *ITGA2*. The four studies of co-expressed *KCNN4*, *ITGA2*, and *PLAUR*, E8 [173], E41[163], E49 [167], and E69 [199], included 2 lung cancers, 1 melanoma, and 1 pancreatic cancer, in the categories of *His/Chr/Epi*, *TP53*, and *RAS* as listed (top left Venn diagram). One study detected *KCNN4*, *LIF*, and *PLAU* together as DEG (top right Venn diagram). One study, E41 [163], detected *KCNMA1*, *KCNN4, ITGA2*, and *PLAUR* together as DEG (not shown). (**B**) PubMed word searches for shared citations among “BK” (encoded by *KCNMA1)* and “PLAUR” + “PLAU” and “ITGA2” found correlations of R^2^ = 0.55, *p =* 0.0175, and R^2^ = 0.52, *p* = 9.0224, respectively, across the 8 categories of 7 oncogenes and histone.

Two studies detected *KCNN4*, *ITGA2*, and *PLAU* (found in 5 studies, ligand for PLAUR) together as DEG (not shown). *PLAUR* and *PLAU* were DEG together in only one study. Ten studies had either *PLAUR* and/or *PLAU* as DEG. Three studies detected *KCNN4*, *ITGA2*, and *LIF* together (not shown) and one study detected *KCNN4*, *LIF*, and *PLAU* together as DEG (Figure 5A, right Venn diagram). One study, E41 [163], detected *KCNMA1*, *KCNN4, ITGA2*, and *PLAUR* together as DEG (not shown).

Searches of PubMed for shared citations in cancer searches with “BK” with “PLAUR + PLAU” and then “ITGA2” in 3-word searches including “cancer” found significant Pearson’s product-moment correlations of R = 0.742 (*p* = 0.0175) and 0.718 (*p* = 0.0224), respectively, and R^2^ = 0.55 and 0.52, respectively, across the 8 categories of 7 oncogenes and “histone” when each was used as a third word (Figure 4B). Although none of the other potassium channels could be checked in this way, repeats in E1-E70 may have value for eventually identifying functionally relevant co-expressed non-KCN DEG’s products to study experimentally with KCN gene products, especially *KCNN4* and *KCNMA1*.

Also, non-KCN DEG found in E1-E70 overlap with those in recent studies of genes encoding circulating proteins associated with risk for numerous types of cancer. The following genes found as DEG in 4 - 6 of the E1-E70 studies, that are shared with the two studies, are as follows with their frequency (X = times): *FGFR3* (5X), *GDF15* (4X), *IGFBP3* (5X), *IGFBP7* (4X), *KRT19* (4X), and *PLAUR* (6X) [224,225].

#### 3.3.10. Non-KCN DEG Superfamilies in E1-E70

Nine superfamilies of non-KCN DEG were detected in E1-E70 with at least 15 occurrences. The instances of superfamily occurrences included as follows: 74X for SLC (Solute Carrier), 32X for COL (Collagen), 27X for ITG (Integrin), 22X for TMEM (Transmembrane), 21X for CD (Cluster of Differentiation), 19X for GPR (G-Protein coupled Receptor), 18X for SOX (Sry-related HMG bOX), 15X for SERPIN (SERine Proteinase INhibitors), and 15X for TNFRSF (Tumor Necrosis Factor Receptor Super Family) superfamily members. DEG repeated in at least 2 studies were present in all of the superfamilies listed.

The largest numbers of repeat non-KCN DEG occurrences were the 7X, 5X, and 4X DEG occurrences of *ITGA2* (previously mentioned), *SOX2*, and *SLC1A2/*Excitatory Amino Acid Transporter 2 (EAAT2), respectively, among the superfamilies. EAAT2 clears glutamate from the extracellular space at synapses [226–228]. *SLC16A3*, monocarboxylate transporter 4, was a DEG in 2 studies as shown (Figure 3B). Six Regulators of G protein Signaling (RGS) family members were also found as DEG in 7 studies. Eighteen (25.7%) of the 70 studies had either GPR or RGS family member DEG. Among the GPR DEG found, none are known as being H^+^-sensitive as described in Appendix B (Text B1). Four carbonic anhydrase genes, *CA2*, *CA9*, *CA12*, and *CA14* were found among 5 studies. *CA2* and *CA9* are shown (Figure 3B). Genes encoding subunits of Na/K-ATPases, *ATP1A2*, *ATP1B1*, and *ATP1B2*, were DEG in 4, 2, and 1 of the E1-E70 studies, respectively (See Fig. 3B). Aquaporin family genes, *AQP1*, *AQP3* and *AQP4*, were collectively DEG in ten of the E1-E70 studies.

#### 3.3.11. KCNs Encoding H^+^-Sensitive Proteins in E1- E70 in Ten Cancer Reviews

Ten reviews describing KCN family members in malignancy were compared with detection of KCN members in E1-E70. The numbers of KCN genes described in reviews are listed as follows: 7 in a review of glioblastomas [229], 3 in a meeting report for a colloquium on cancer [230], 23 in a review listing 18 cancer types [231], 13 in a review that listed 10 cancer types [232], 9 in a review mentioning 17 cancer types [233], 21 in a review that listed 14 cancer types [234], 12 in a review mentioning at least 13 cancer types [235], 15 in a review that listed 7 cancer types [236], 13 in a review that listed 13 cancer types [237], and 13 in a review that mentioned 11 cancer types [238]. Twenty KCNs found in 35 (one half) of E1 - E70 studies overlap with those in the reviews and are listed as follows: *KCNA1*, *A3*, *A5*, *D1*, *H1*, *H2*, *H7*, *J5*, *J8*, *J10*, *K2*, *K3*, *K5*, *K15*, *MA1*, *N2*, *N3*, *N4*, *Q1*, and*T2*. The ten reviews combined with E1-E70 highlight KCN genes that encode H^+^-sensitive proteins. At least one KCN gene in each of 29 studies (82.85% of 35 overlapping studies from E1 – E70) encodes a H^+^-sensitive protein and these studies are listed as follows: E4, E5, E6, E8, E9, E12, E13, E16, E22, E24, E28, E31, E32, E34, E36, E38, E39, E40, E41, E47, E49, E52, E53, E58, E60, E64, E68, E69, and E70. The remaining overlapping studies included E15, E29, E33, E45, E65, and E66. Citations for E1-E70 are in *Section 3.3.6. Categories Among Studies with E1-E70 Citations Per Category*.

#### 3.3.12. Long Introns (20,000 bp or longer) in E1-E70’s KCN DEG

Many genes in the KCN family have long introns (at least 20,000 bp in length). Long introns can affect gene expression and may play a role in gene regulation. Also, they may be affected by genetic damage as cancer develops and evolves. Thirty-one (58.5%) of the 53 KCN DEG detected in E1-E70 contain long introns, some much greater than 20,000 bp. Seven KCN DEG (*KCNC2*, *D3*, *H1*, *H8*, *MA1*, *Q1*, and *T2*) each contain at least one intron that is 100,000 – 199,999 bp in length. Five KCN DEG (*KCNH*7, *IP1*, *Q3*, *MA1*, and *MB2*) each contain at least one 200,000 - 299,999 bp intron and *KCNB2* and *KCNQ5* each contain at least one intron that is 300,000 – 399,999 bp long. Fourteen of the KCN DEG in E1-E70 contain multiple long introns. *KCNMA1* and *KCNT2* each contain seven long introns. *KCNN4* is among the KCN DEG that do not contain any long introns. Forty-one (58.6%) of the E1-E70 studies contained at least one KCN DEG that had at least one long intron. Nine studies had multiple (2 – 4) KCN DEG with long introns. However, *KCNA3*, *A5,* and *F1* in E1-E70 have only one exon and therefore do not contain any introns. Intron data from the Ensembl database, was obtained in 2023 and 2025, last accessed 6/22/2025, Ensembl browser 114 (May 2025 release). The contrast between the introns of *KCNMA1* and *KCNN4* is striking and raises the possibility of increased preservation of *KCNN4* as a smaller target when genetic damage occurs but this is unknown.

### 3.4. Study of Selected KCN DEG in REMBRANDT

#### 3.4.1. Rationale and Preliminary REMBRANDT Survey

Based on glioblastoma cell adherence to dying cells, “palisading” around necrotic tumor regions, chemotaxis to serum and possibly a potassium gradient, and the tendency of glioblastoma cells to invade white matter tracts, the possibility of potassium acting as a chemoattractant in the CNS biological milieu led to a search for KCN DEG in the brain tumor database, REMBRANDT, concurrently while searching for KCN DEG in studies of various cancer types already described. REMBRANDT (as a part of the online Georgetown Database of Cancer (G-DOC) was queried. Among potassium channel related genes, KCN family members were selected because of their H^+^- sensitive and modulatory/regulatory members, independence from energy requirements for many members, susceptibility to genomic damage due to the wide genomic distribution of the large KCN family (at least 90 members), potential to be downstream effectors of oncogenes, possible interactions with G-proteins, complex relationships with biological milieus, and additionally their major roles in glial and neuronal functions. Initial queries in the Online Mendelian Inheritance in Man (OMIM) database (https://omim.gov) in 2018-2019 and literature descriptions of KCN genes [239] yielded 41 KCN genes with neuronal as well as astrocytic expressions in the human brain to consider. All 41 genes were found in loci that corresponded to known regions of chromosomal losses, gains, or both in glioblastomas, published as Figure 3-76 in a book authored by Burger PC and Scheithauer BW [1]. These genes are distributed as follows: Chromosome (Ch) 1 (*KCNA2*, *A3*, *C4*, *N3*, *J9*, *J10*, *K2*, *K1*); Ch2 (*KCNJ3*, *J13*); Ch3(*KCNMB2*, *MB3*); Ch5 (*KCNN2*, *MB1*); Ch6 (*KCNK17*), Ch8 (*KCNQ3*); Ch9 (*KCNT1*); Ch10 (*KCNMA1, K18*), Ch11 (*KCNJ11*, *C1*, *A4*, *K4*, *J5*); Ch12 (*KCNA1*, *A5*, *A6*, *J8*, *MB4*); Ch14 (*KCNK10*), Ch17 (*KCNJ12*, *J16*, *J2*); Ch19 (*KCNN1*, *N4*, *C3*); Ch20 (*KCNB1*, *Q2*); Ch21 (*KCNJ15*, *J6*); and Ch22 (*KCNJ4*). The most numerous were in the KCNJ subfamily genes (*KCNJ2*,*3*,*4*,*5,6*,*8*,*9*,*10*,*11*,*12*,*13*,*15*, and*16*) encoding inwardly rectifying potassium channel forming proteins(Kir) and their modulators/regulators, Kir2.1, 3.1, 2.3, 3.4, 3.2, 6.1, 3.3, 4.1, 6.2, 2.2, 7.1, 4.2, and 5.1. Kir channels can be relatively occluded by Mg^2+^ or polyamines when cell membrane potentials are positive relative to the equilibrium potential of K^+^ that allows only a small outward current. At membrane potentials negative to the equilibrium potential of K^+^, Mg^2+^ and polyamines leave the channels by flowing into the cell so a large inward K^+^ current can occur [240,241]. Kirs, including Kir4.1, may also extrude K^+^. Kir 4.1 and 5.1 are present in both astrocytes and oligodendrocytes [39,242].

A preliminary survey of REMBRANDT for expression of 21 KCN family members,12 other potassium flow related genes, and 4 housekeeping genes (HKG) checked for differences in glioblastomas from patients, with and without a history of seizure medications. Seizures could be an indicator for abnormal levels of potassium [243–245]. With “unknowns for gene expression” included, there were 84 and 18 glioblastomas, with and without a history of anti-seizure medication, respectively. Compared to the absolute difference in the expression of HKG (*PPIA*, *RPLP0*, *YWHAZ*, and *B2M*) between the two groups, the single or average absolute median values of reporter probes for *KCNJ10*, *KCNK3*, *ATP1A1*, *KCNN3*, *KCNJ13*, *KCNMB2*, *SLC12A7*/KCC4, *KCNN2*, *KCNMB1*, *GJA1*, *KCND2*, *KCNK9*, *SLC1A2*/EAAT2, *KCNJ2*, and *KCNJ16* exceeded the average HKG difference (0.1131) of median values. Others, including *ATP1A2*, *KCNA2*, *KCNA3*, *KCNK1*, *KCNIP3*, *KCNMB3*, *SLC1A3*/EAAT1, *SLC12A2*/NKCC1, *KCNT1*, *KCNMA1*, *LRRC55*, *KCNA1*, *KCNMB4*, *GJB2*, *GJB6*, *SLC12A6*/KCC3, and *FXYD7*, did not exceed the average HKG difference in expression. The preliminary data (not further analyzed) suggested that the subset of glioblastomas in patients with seizures (according to medication history) had expressions of KCN family member genes that could be compared to the same genes in oligodendrogliomas (less invasive glial tumor) from patients who also had seizures. Abnormalities in potassium-related gene expressions, such as among the KCN family members, resulting in potassium flux related to worse malignant behavior could potentially underly differences in behavior between the related gliomas, oligodendrogliomas and glioblastomas.

#### 3.4.2. Specificity in Gene Expressions Reflecting Cell of Origin

Checks were made for expected findings. Expression of *ATP1A2* (alpha subunit of Na/K-ATPase specific for glial cells) was higher in both types of glial tumors versus its neuronal specific counterpart (*ATP1A1*) in the patients on anti-seizure medications. The medians of two reporter probes available for each gene yielded significant differences, *p* = 0.0345 and 0.0278, respectively, for oligodendrogliomas and glioblastomas using 2- tailed, unpaired t-tests (Figure 6A). The specificity of expression for connexin genes in astrocytic cells, *GJA1*/Cx43, *GJB6*/Cx30, and *GJB2*/Cx26, was used to verify the expected cells of origin for the two types of glial tumors, oligodendrogliomas and glioblastomas [242,246,247], in REMBRANDT patients on anti-seizure drugs. The medians of single reporter probes for expression of astrocytic type connexins, collectively as a group, were significantly greater in the glioblastomas, *p* = 0.0477 (1-tailed, paired t test), (Figure 6B). Results using multiple reporters for oligodendrocyte type connexins were not available. More prominent vascularization of glioblastomas versus oligodendrogliomas was consistent with a trend for greater expression of *VEGFA*, *p* = 0.166, in the glioblastomas among the patients with known anti-seizure treatment (not shown).

**Figure 6.**
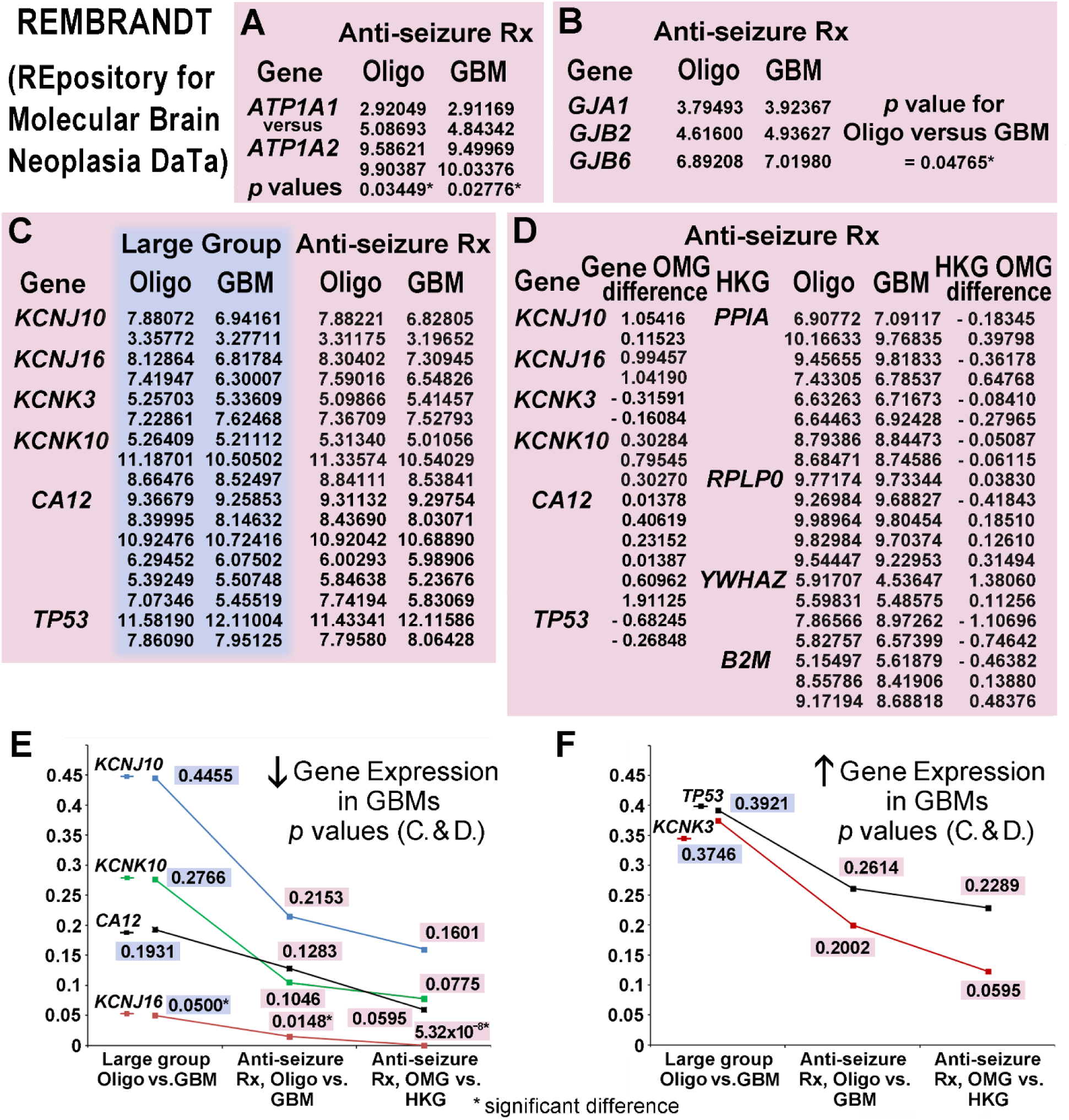
Results for oligodendrogliomas (Oligo) and glioblastomas (GBM) from REMBRANDT. (**A – D**) Median reporter probe values of microarray gene expression results from Oligo and GBM tumor tissue are shown in columns as indicated. Pink backgrounds (A – F) indicate results are from patients on anti-seizure medications (Anti-seizure Rx). Results with a blue background (C, E, F). are from Large Group patients (medication history not used for criteria). (**A**) The *p* values of t-tests are at the bottoms of columns for comparisons of *ATP1A1*/Na/K-ATPase subunit alpha 1, neural specific, versus *ATP1A2*/Na/K-ATPase subunit alpha 2, glial specific in Oligo and GBM with significant differences confirming glial origins. (**B**) Single median reporter probe values for astrocytic-specific (versus oligodendroglial) values for gap junction genes with higher levels of expression in GBM versus Oligo as expected. (**C**) Values listed for median reporter probes for 4 KCN genes (See text) and 2 other genes, *CA12*/carbonic anhydrase 12 and *TP53*/p53. (**D**) An additional method to evaluate expression of genes shown (C), calculated Oligo Minus GBM (OMG) differences of the median probe results for each gene of interest and the housekeeping genes (HKG). These differences were then compared to correct for variation in tissue handling, assay-to-assay variation within and between institutions, etc. (**E – F**) The *p* values are shown graphically for decreased and increased gene expressions in GBM, respectively. Significance improved dramatically for *KCNJ16* with this approach and trends became stronger for other genes. Oligo = oligodendroglioma, GBM = glioblastoma, Rx = medication, HKG = housekeeping genes. REMBRANDT (part of the Georgetown Database of Cancer (G-DOC).

#### 3.4.3. KCN Gene Expressions in Patients on Seizure Medications

Limiting tumor comparisons to patients on anti-seizure medications improved detection between the two types of invasive glial tumors. Without using the history of seizure medications, a difference in expression of *KCNJ16*/Kir5.1 in glioblastomas (less expression), *p* = 0.05, of borderline significance, became significant in glioblastomas using the smaller group of patients on anti-seizure drugs, *p* = 0.0148. Trends also improved for differences in expressions of *KCNJ10*Kir4.1, *KCNK3*/TASK1, and *KCNK10*/TREK2, (all H^+^-sensitive, Appendix B (Table B2) and [248,249]) with reporter probe data (Figure 6C) and *p* values (Figures 6E - F) shown along with other genes, *CA12*/carbonic anhydrase 12 and *TP53*/p53. *SLC16A1*/MCT1, *PTEN*, and other non-KCN genes (not shown), also showed some improvement in the difference between oligodendrogliomas and glioblastomas when comparisons were limited to patients on anti-seizure drugs. Median results of five *SLC16A1*/MCT1 probes indicating increased expression in glioblastomas, but without significance, had *p* = 0.269 and *p* = 0.255 in the smaller group with a history of anti-seizure medication.

#### 3.4.4. <u>O</u>ligodendroglioma <u>M</u>inus <u>G</u>lioblastoma (OMG) Normalization

Housekeeping gene (HKG) normalization was used to correct for loss of genomic integrity, differences in tissue handling, assay to assay variation within and between institutions, etc. in the comparisons using the microarrays of two different type of tumors in REMBRANDT. It was used for comparisons in patients on anti-seizure medications. <u>O</u>ligodendroglioma <u>M</u>inus <u>G</u>lioblastoma (OMG) differences (increased or decreased) were calculated for each reporter probe’s median expression. These OMG differences for each DEG were compared to the OMG differences for all 4 HKGs. Enhancement of the previously detected differences resulting in better *p* values as shown (Figures 6D-F). The OMG difference for *KCNJ16*, decreased in glioblastomas, became highly significant, *p* = 5.532 x 10^-8^. The trends for several other genes improved. The *t*-tests were calculated for unpaired samples with unequal variance in the OMG comparisons. The median probe data (Adobe Flash format) in Figures 6A-D is no longer available online in REMBRANDT (Georgetown website, see Materials and Methods, Subsection *2.3. Databases*).

#### 3.5.1. KCN DEG Shared in E1-E70, REMBRANDT, and Glioblastoma Cell Lines [22]

Whole exome and RNA sequencing data of LN18, LN229, T98G, U87, U343, and U373 human glioblastoma cell lines, available as Open Access in Patil et al.’s Supplementary Materials [22], revealed data for many of the KCN DEG and genes of interest found in the E1-E70 and REMBRANDT studies. Also, 3 and 5 of these cell lines were also used for the cell settling and cell migration studies, respectively, described earlier. Their data availability is described in the **Data Availability Statement** that follows this article’s text. They described their methodology in detail [22], with some information also included here. For each of the GBM cell lines, U87, T98G, LN229, LN18, U343, and U373, a 62 Mb region of the genome was targeted. Alignment was to human reference genome hg19. Their single nucleotide variant (SNV) was considered to be a potential Cancer Specific Mutation (CSM) if present in the COSMIC database (Reference 18 in [22]) and/or was present in GBM TCGA exome sequencing data (their references 8,9). Data for 4 cell lines, U87, T98G, LN229, and LN18, had an average concordance of 65.3% with the COSMIC database (their reference 18) and 74.74% with the Broad Institute’s CCLE database (their reference 19). In addition to CSMs, Patil et al. reported small insertions and deletions (indels) in a size range of −49 to +29 bases. Differential gene expression data per Patil and colleagues was obtained via comparisons of their RNA-seq data from six GBM cell lines, U87, T98G, LN18, LN229, U343, and U373, with expression data of five normal brain tissue samples from TCGA, with a cut off value of a 2-fold change in absolute expressions. Their results for 2,214 genes compared with those that were also present in TCGA GBM tumor samples had a concordance of approximately 73%. Of the 2,214 genes in the Patil study,1,831 (82.7%) were down regulated. For the 1,621 genes similarly up or down regulated in both the TCGA and Patil et al. studies, 79.64% were down regulated.

Among the KCN genes identified by Patil and colleagues, 42 KCN genes had CSM and indels. The DNA changes identified in 18 of the 42 KCN genes occurred in at least 3 of their glioblastoma cell lines. The KCN family group, with the highest number of members (ten) with reported DNA changes, included *KCNJ1*, *2*, *3*, *5*, *6*, *10*, *12*, *13*, *14*, and*16*. Patil and colleagues identified 69 CSM among KCN family member genes, representing 4.33% of 1,594 CSM found in all the genes they sequenced. Also, the 68 indels (25 insertions and 43 deletions) that they found in KCN family members represented 1.75% of all 3,892 indels found in the 6 glioblastoma cell lines. For comparison with this study of KCN DEG in E1-E70 and genes of interest in REMBRANDT, shared KCN DEG with their sequencing results revealed DNA alterations including CSM, INS, and DEL. Twenty-three of the E1-E70 KCN DEG and 4 of the REMBRANDT study’s KCN genes of interest based on expression had changes described in the genomes of 6 glioblastoma cell lines by Patil et al. Those detected in E1-E70 (glioblastomas (7) with 28 other cancer types) are marked with light green squares on the right side of lists, and genes of interest (significant difference or trends) found in REMBRANDT are bolded (*KCNJ10*, *KCNJ16*, *KCNK3*, and *KCNK10*), (Figure 7A). Patil et al. found that all 51 KCN family members with gene expressions detected in the six glioblastoma cell lines were down regulated versus normal with the 51 expressed KCN family member genes representing 1.48% of all 3428 genes found to be differentially expressed in the cell lines. Average gene expression levels of RNA determined by Patil et al. are listed in descending order of abundance, including 27 and 4 of the E1-E70 and REMBRANDT KCN genes, respectively, along with names of encoded proteins (Figure 7B). *KCNJ16*, which was significantly down regulated in REMBRANDT’s glioblastomas, when compared to oligodendrogliomas, was ranked among the lowest detectable expressing genes in the 6 glioblastoma cell lines studied by Patil et al. Only four of the 51 KCN genes with expression values ranging from <0.001 to 803.719 (modeled counts) had lower expression than detected for *KCNJ16* (Figure 7B). The log2 fold RNA changes from normal calculated using their Supplementary Table S6 [22], are shown for the 51 KCNs with detectable RNA in the glioblastoma cell lines. The 27 genes (52.94%) with encoded H^+^- sensitive proteins and the 24 KCN genes with DNA changes are highlighted on the graph (Figure 7C). Of the 53 KCNs in E1-E70 there are 38 (71.70%) also described by Patil et al. in their glioblastoma cell lines, either with the DNA changes shown and/or with RNA expressions detected. Of the seven GBM studies in E1-E70, six (85.71%) each shared 1 - 3 KCN DEG with those described by Patil et al. The specific glioblastoma studies in E1-E70 with these KCN DEG are E5, E21, E26, E27, E31, and E51. Only one, E50, of the E1-E70 glioblastoma studies did not detect a KCN DEG corresponding to those with RNA expression detected by Patil et al. [22]. There is a large overlap of KCNs encoding proteins with H^+^-sensitivity in E1-E70, REMBRANDT, and those described by Patil et al. (Figure 7C). References for H^+^-sensitivities are cited in Appendix B, Table B2, and earlier in Section *3.4.3. KCN Gene Expressions in Patients on Seizure Medications,* for the encoded proteins of *KCNA1*, *KCNA3*, *KCNA5*, *KCNAB2*, *KCNH1*, *KCNH3*, *KCNJ10*, *KCNJ16*, *KCNK1*, *KCNK3*, *KCNMB4*, *KCNN3*, *KCNQ1*, *KCNQ2*, *KCNQ3*, *KCNQ5*, and *KCNT1*. References for H^+^-sensitivity of the proteins encoded by additional KCN genes shown (Figure 7C) are as follows: *KCNA2*/Kv1.2 [250], *KCNA4/*Kv1.4 [251,252], *KCNC4*/Kv3.4 [253], *KCNH4*/ELK1 [254], *KCNJ3*/GIRK1 [255,256], *KCNJ11*/Kir6.2,IKATP [257,258], *KCNJ12*/Kir2.2,HIRK1 [259] (variant), *KCNK9*/K2P9,TASK3 [260,261], and *KCNK17*/K2P17.1,TALK2,TASK4 [262].

**Figure 7.**
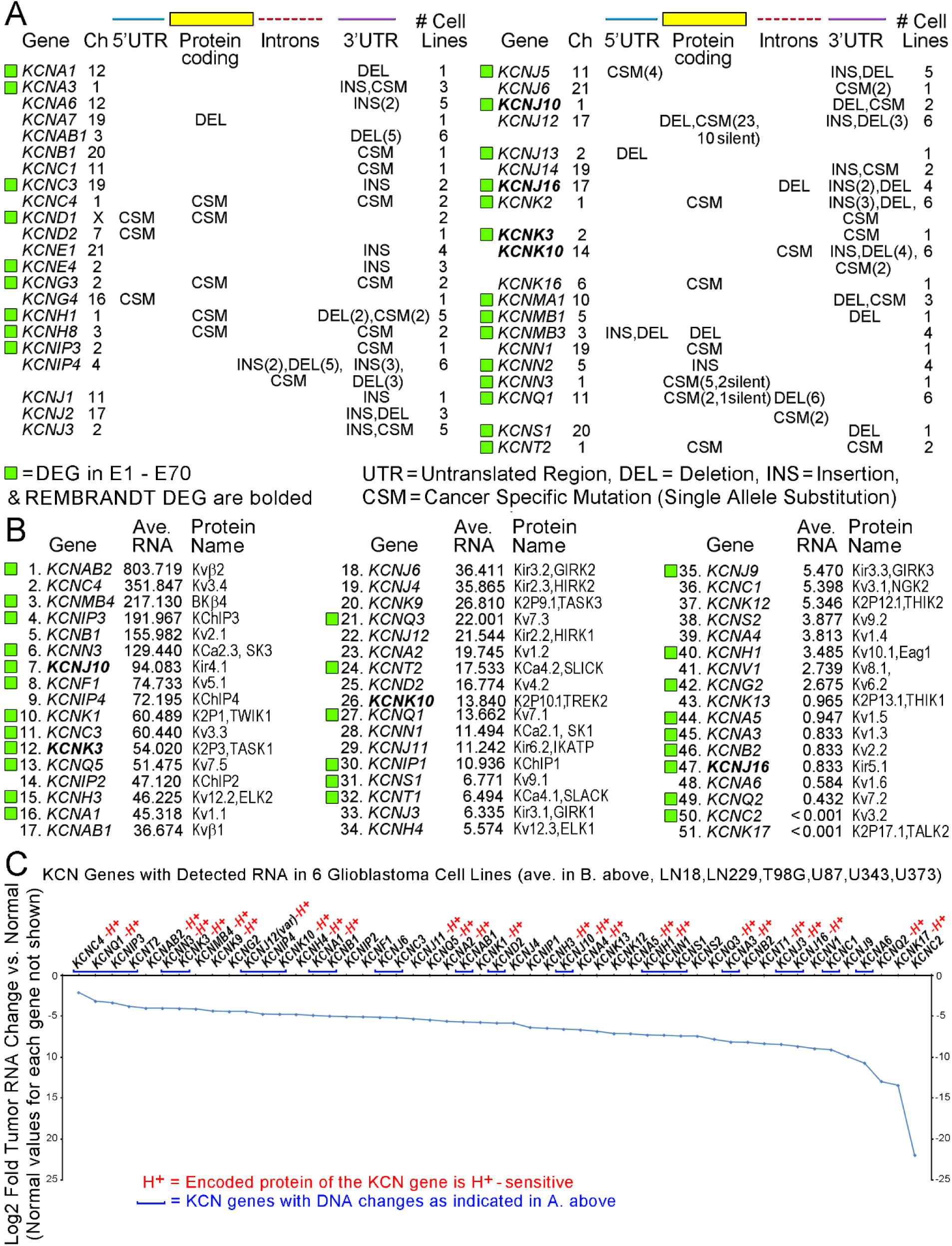
Many KCN DEG detected in E1-E70 studies (Figures 3 - 5). and in REMBRANDT (Figure 6). were included in whole genome and expression studies of 6 glioblastoma cells lines by Patil et al. [22]. They reported on LN18, LN229, T98G, U87, U343, and U373 cell lines. For comparisons, the shared KCN DEG detected in E1-E70 (29 cancers including 7 glioblastomas) are marked with light green squares and the KCN DEG found in REMBRANDT are bolded (**A** and **B)**. (**A**) Sequencing results revealed DNA alterations including cancer specific mutations (CSM), insertions (INS), and deletions (DEL). Twenty-three KCN DEG (of 53 in E1-E70, See Figures 3 and 4) and those with potential significance in REMBRANDT (3 with trends and 1 significant KCN DEG, See Figure 6C - F), were detected with changes in the genomes of 6 cell lines (usually > 1) by Patil et al. Several shared KCN DEG were genetically altered in all 6 cell lines. (**B**) Average gene expression levels of RNA determined by Patil et al. are listed in descending order of abundance, including 27 and 4 of the E1-E70 KCN DEG and REMBRANDT’s KCN DEG and genes of interest, respectively, along with names of their encoded proteins. (**C**) Log2 fold RNA changes in the combined tumor cell lines versus normal (see text) are shown graphically. KCNs encoding H^+^-sensitive proteins are highlighted as red (H^+^) and the KCNs found to have DNA changes in (A) are highlighted with underlining blue brackets (˽). Of the 51 KCNs with detectable RNA in the glioblastoma cell lines, 27 (52.94%) encoded proteins that are H^+^- sensitive. Of the 53 KCNs in E1-E70 there are 38 (71.70%) also described by Patil et al. in glioblastoma cell lines, either with altered DNA and/or RNA expression values. Of the seven GBM studies in E1-E70, six (85.71%) shared 1 - 3 KCN DEG with those described by Patil et al. [22]. See the text and Appendix B (Figure B1A) for specific glioblastoma studies in E1-E70 and the references for each study are given in Section *3.3.6. Categories Among Studies with E1-E70 Citations Per Category.*

## 4. Discussion

### 4.1. Input of Functional U87 Studies

The decision to stochastically investigate potassium in cancer was based on earlier FBS cell migration assays and then observing cytology of glioblastoma cell in settling assays. After fluid was wicked away as cells settled on glass slides, not only was expected cell death and cytoplasmic vacuolization seen, but also small groups of cancer cells clustered around single dying cells. Although malignant cells from body fluid specimens sometimes ingest whole cells, the tight cell adherence of robustly viable cells to dying settled cells that were formerly in cell suspension was a surprise. Uncentrifuged cancer cells from clinical body fluid specimens viewed microscopically, after gentle filtration and dissolution of filters [263,264], do not have this appearance, at least routinely. However, the morphology of the settled, viable cells clustered around single necrotic cells did resemble characteristic rimming of deeply staining glioblastoma cells (“pseudopalisades”) around necrotic foci in glioblastoma tumor tissue as described in the Introduction, Section *1.1. General*. Also, basophilic staining in tumor cells of tumors with central necrosis with more alkaline cytoplasmic pH detected in necrotic areas has been established experimentally in cervical cancer cells [265]. Clues, as mentioned earlier, to form a hypothesis for the glioblastoma cells in settling assays and their rimming around necrosis in tissue included (*i*) the viability of cancer cells rimming single dying cells seemed protected from the negative effect of fluid removal, (*ii*) relatively darker staining of some viable clustered tumor cells possibly reflected a pH change in them, and (*iii*) when ATP is depleted, cells open their cation channels to release K^+^ (major cation) before rupturing [266–268]. Potassium efflux has been seen in cancer cells [269].

A chemotactic role for potassium was supported in view of cell migration assays, previously performed for planning the use of FBS to optimize glioblastoma cell lines migrating to HGF. Results could not rule out an unidentified serum constituent acting as a chemoattractant. Multiple cell lines migrated to FBS alone with dose responses. Also, U87 cells migrated significantly (*p* = 0.0144) with a positive gradient present of K^+^ generated by MEM (5.3 mM K^+^ in a milieu of amino acids, vitamins, etc.) versus 4.5 mM K^+^ in “Migration” media, in the absence of FBS and HGF, and was improved in the presence of FBS (Figure 2E). Support for potassium as the chemoattractant possibly inciting their reversal of pH is also considered.

### 4.2. Mechanistic Proposal for a New Route to pH Reversal

The KCN family members, many pH sensitive, cooperate to control potassium levels at cell membranes in a putatively H^+^-sensitive manner. A new hypothesis is that K^+^, as a monovalent exchange cation for H^+^, frees bound protons from inner cell membranes for rapid cytoplasmic transfers to proton extruders, and instigates pH reversal in cancer if the control of potassium flux is deregulated. Whether rapid delivery of H^+^ to exit sites occurs via the Grotthuss mechanism during pH reversal is also a new consideration that could explain how freed protons become available for rapid cytoplasmic proton transfers to cell exit sites during pH reversal in at least some malignancies. Although the Grotthuss mechanism of rapid proton transfer via “water wires” has been described for proteins and the tight junction of the blood brain barrier [270–272], cytoplasmic Grotthuss type proton transfer in cancer cells has not been recognized. As migrating tumor cells flatten and squeeze into small spaces such as between a nerve fiber and its surrounding nerve sheath or along and then through dense basement membranes, bone, cartilage, etc. there should be microaqueous intracellular regions of tumor cells generated temporarily. The cell settling assays removed fluid to create a microaqueous condition for all cells with simultaneous exposure to K^+^ from single dying cells within some clusters of cells. These centralized dying cells could also have become “sinks” for protons released from the surrounding viable tumor cells.

Putatively, entry of K^+^ across membranes into the viable cells in settling assays provided replacement ions for H^+^ bound to fixed inner membrane anions (and possibly to H^+^ bound to cytosolic proteins) that permitted newly freed H^+^ (rapidly formed H_3_O^+^) to diffuse through the cytoplasm, possibly via cytosolic Grotthuss water wires formed under microaqueous conditions to serve as a “root system” for supplying protons to extruders. In tissue, viable microaqueous malignant cells may use a potassium flux, K^+^/H^+^ exchange, proton transfer in water, and then cell exit of protons as a response to stress or oncogenes to acidify the microenvironment with activation of proteases, etc. for cell migration. Simultaneously a higher cytosolic pH is provided to activate alkaline-sensitive intracellular proteins promoting cell survival. An alkaline pHi leads to maximal tumor cell proliferation coupled with increased glycolysis and adaptation to hypoxia *in silico* [273] and is associated with increased flux of glycolytic enzymes, migration, and invasion experimentally [274]. Also, when pHi increases, H^+^ diffusion increases [275], possibly enhancing pH reversal for survival of single, migrating cancer cells.

### 4.3. Further Consideration of KCN Genes in Cancer, Including Key Negative Findings

Simultaneous studies of multiple KCN genes of interest in REMBRANDT and those in KCN cancer reviews, including those with H^+^-sensitivity, supported the ongoing stochastic investigation of KCN family members as DEG in unbiased cancer studies. The relatively large number of studies allowed investigation of H^+^-related associations with the expression of the KCN gene family’s numerous members differentially expressed in malignancy. KCNs in E1-E70 studies overlapped with KCN genes of interest in REMBRANDT (*KCNJ16*, *KCNJ10*, *KCNK3*, and *KCNK10*, all 4 encoding H^+^-sensitive proteins), and with 20 KCN genes covered in ten reviews. At least one KCN DEG in each of 29 studies (82.85% of the 35 studies from E1-E70 overlapping with the KCNs in ten reviews) encodes a H^+^-sensitive protein. Potassium ions may redistribute in cancer cells to exchange with H^+^ in pH reversal via differential expression in the KCN family members that encode H^+^-sensitive proteins, possibly related to oncogenes, genetic damage, etc.

The KCN family offers the possibility of a large variety and range of dynamic distributions of K^+^ via its numerous members, networking, biological milieus, modulation by oncogenes, histone/DNA modifiers, etc. Also, potassium is supplied by blood pulsing through the vascular system, although locally haphazard in cancer, and by the firing of action potentials along axons in nerve fibers and white matter tracts. The formation of synapses between tumor cells and neurons provide K^+^ as well as other synaptic substances [276]. These sources may be increased or decreased in cancer. Activation of proteases by extracellular acidosis that degrade connective tissues provide increased access to neural and vascular K^+^ during migration and metastasis. Multiple proton extruders have been proposed for glycolytic/plurimetabolic type glioblastoma cells [277]. Also, the microaqueous surface conditions in cell settling assays and during tissue invasion *in vivo* suggest a lower probability of CO_2_ dissolving in surrounding extracellular fluid to form carbonic acid that dissociates to hydrogen and bicarbonate ions for cell uptake. The possibility of having taurine function significantly as a carrier of cytoplasmic H^+^ seems less likely since the only taurine transporters found as DEG in E1 – E70, *SLC16A6* and *SLC36A1* (or *PAT1*, former name), were present in low numbers (Figure 3B). No non-KCN DEG were found for carnitine transporters. Adaptations took place in the cell settling assays and in some invasive scenarios without extracellular fluid being available. Also, the aquaporin genes, *AQP1* and *AQP4* genes, whose products mediate water flux, were only found in 3 of the E1-E70 studies. *AQP3*, found in 4 studies, encodes aquaglyceroporin that mediates glycerol, uncharged solutes, and water flux [278].

In the E1 - E70 studies important negatives include non-KCN DEG representing proton extruders, carbonic anhydrases, etc. with some mentioned in the Results and earlier in this section of the Discussion. Importantly, only a few studies found *ATP1A2* encoding Na/K-ATPase and related genes (as a group < 10%) to be differentially expressed in E1- E70. Its role as an important negative is supported by a report that digoxin, an inhibitor of Na^+^/K^+^ ATPase, an active inward K^+^ transporter, has been found to reduce the size of circulating tumor cell clusters in women with metastatic breast cancer, −2.2 cells per cluster [279]. Others are also shown (Figure 3B). Some negatives (lacking significant differential expression) were described in the REMBRANDT analysis.

In support of finding *KCNK* genes as DEG in 20% of E1-E70, with *KCNK1*/K2P1 being most frequent, this is consistent with the loss of *KCNK1* being associated with breast cancer cell survival when necrosis was induced [118]. Also, the ten reviews of KCN genes found in cancer as described in the RESULTS, Section 3.*3.3.11. KCNs Encoding H^+^- Sensitive Proteins in E1-E70 in Ten Cancer Reviews*, included overlap with 4 members of the K2P (*KCNK* genes) and 4 members of the KCa (*KCNM* and *KCNN* gene groups) subfamilies and others. In E1-E70, *KCNN4* and *KCNMA1*, were the most frequently detected KCN DEG, 9 and 6 studies, respectively.

### 4.4. E1-E70 Studies Related to 7 Oncogenes and Histone/DNA Modulation

Among the E1-E70 studies, 50 were in 8 categories of histone/DNA modulator involvement and/or having relationships to one or more of 7 oncogenes (*TP53*, *TGFβ*, *EGFR*, *MYC*, *RAS*, *BCL2*, and *MET*). Sixteen studies were in more than one category. Migration of 7 of the 8 glioblastoma cell lines responded to HGF, the ligand of *MET*’s encoded Met [280,281] as shown (Figure 2). Others have proposed a role of multiple oncogenes, including *EGFR*, *RAS* (*KRAS* and H*RAS*), *MYC*, and *TGFβ*, that overlap with the oncogenes in E1-E70 studies, and histone/DNA modifiers to interact on paths to malignancy in a context driven manner [282]. Lactate can activate *TGFβ* [283–286]. Also, “upregulation of oncogenic pathways” has been described in clinically evident precancerous lesions of 12 different types of malignancies in studies reviewed recently [287].

The collective influence of multiple oncogenes on downstream potassium channels may promote K^+^ flux at inner membranes of tumor cells, possibly with refilling needed if outflow via large flow K^+^ channels is anomalously high. Oncogenes could be involved upstream in promoting K^+^/H^+^ exchange leading to pH reversal. The 50 KCN studies related to 7 oncogenes and/or histone/DNA modulation were significantly (*p* = 0.0325) more likely to have at least one or more KCN DEG encoding a H^+^-sensitive protein than the other 20 KCN studies in “no category”.

Retrieval of citation data from PubMed revealed a significant correlation in the numbers of studies of H^+^ with K^+^ across all 8 categories of 7 oncogenes and histone in 2- word searches without restriction to cancer. However, the category for MET had to be removed to achieve a comparable correlation of H^+^ with K^+^, when searches were restricted to “cancer” using 3-word searches (Figure 4C). Although explanations require definitive studies, this may reflect under-reporting of negative studies. The PubMed citations reveal that citations for BK (*KCNMA1* encoded) probably constitute a large proportion (approximately one third) of studies involving K^+^ and H^+^ across the categories of 7 oncogenes and “histone” not restricted to cancer and the BK citations may represent most of the studies when restricted to “cancer” (Figure 4B-C). Findings in this study are consistent with at least some H^+^-sensitive KCN DEG in addition to *KCNMA1*/BK being influenced by six oncogenes (those listed previously but not MET) and histone modulation related to K^+^ flux (possibly via activation of ACLY, see *Section 1.3. Metabolic Need for Proton Extrusion in Cancer*). Oncogenes and histone modulation through KCN genes/proteins may play a role in H^+^ dynamics.

### 4.5. Non-KCN DEG in Multiple E1-E70 Studies

In E1-E70, the most frequent non-KCN DEG found was *ITGA2* in 7 studies, and *HMGA2*, *LIF*, and *PLAUR* were each found in 6 studies with *ITGA2*, *LIF*, and *PLAUR* shown in Venn diagrams (Figure 5A). Also, to support the relevance of cancer gene expression studies with unbiased detection of KCN DEG, six non-KCN DEG, i.e. *FGFR3*, *GDF15*, *IGFBP3*, *IGFBP7*, *KRT19*, and *PLAUR*, each included in at least 4 of the E1-E70 studies, were also identified by others as encoding circulating proteins associated with a risk of cancer [224,225]. *PLAUR*, found in 6 of E1-E70 studies, was also co-expressed with the most frequent KCN genes that are also H^+^-sensitive, *KCNN4* and *KCNMA1* in 4 and 2 studies, respectively. *KCNN4*, *PLAUR*, and *PLAU* were detected together as DEG (Figure 5A) in E49 [167], a study associated with 3 categories, RAS, TP53, and His/Chr/Epi. BK (encoded by *KCNMA1*) citations in PubMed correlated with those for PLAUR + PLAU and ITGA2 (Figure 5B).

### 4.6. Specific Roles for Redistribution of K^+^ in Cancer

Studies on intracellular pH-regulating systems, potassium channel inhibitors have been used with interesting results, including “slower recovery from CO_2_ acidification” in early frog skeletal muscle experiments [288]. Potassium channels have been recognized as playing important roles in cancer, including volume changes [289,290] but without also considering K^+^ for exchange with H^+^ on fixed anions to then incite cytoplasmic proton transfers to cell exit sites. The intermediate step of rapid cytoplasmic proton transfer is plausible. Volume changes in cancer cells, problems in their blood supply, stiff tissues constraining migrating cells, etc. that lead to microaqueous conditions in the cytoplasm may help generate Grotthuss conditions that build a putative water-wire type root system to supply proton extruders.

Removal of K^+^ from cell interiors may occur via flux of unsolvated K^+^ through KCN channels after it has lost its aggregated water. Large outward (*KCNMA1*/BK), intermediate outward (*KCNN4*/IK), bidirectional (*KCNJ10*-*KCNJ16*/Kir4.1-Kir5.1 dimers or homomeric versions), etc. may alter cytosolic flows to bring centrally located intracellular water via K^+^-7H_2_O (loosely aggregated) to the cell periphery where water wires form to provide rapid proton transfers. Alternatively, simultaneous changes in Cl^-^ channels (CLCN detected only once as a DEG in E1-E70 was *CLCN7*) and aquaporins (only a few DEG in E1-E70, see earlier Discussion) could help redistribute intracellular water but stochastically their participation as major players seemed unlikely in E1-E70 based on DEG detected. Na^+^ binds water tightly so it is probably not responsible for water redistribution (See Introduction *1.4. Complexity of K^+^ Flux,* 4^th^ paragraph for related information on Na^+^). Redistributed water regionally within cells may promote appropriate Grotthuss conditions, especially during cell migration but is not established as doing so.

K^+^ released outside the cells through high volume calcium gated K^+^ channels, etc. could be recycled via refilling through KCN channels (if available) or by Na/K-ATPase that brings K^+^ in across cell membranes that could also displace fixed H^+^ ions. Accumulated K^+^ near the high flow outward channels prior to its release may displace fixed H^+^ ions. The intermediate and large conductance K^+^ channels encoded by, *KCNN4* and *KCNMA1*, respectively, found 9 and 6 times, respectively, and other KCNNs (*KCNN2*, *3*) and the KCNMBs (*KCNMB1*, *2*, *3*, *4* whose products regulate *KCNMA1*/BK) were found as DEG in 5 and 8 studies, respectively, (only a few in the same studies) in E1-E70. Kir4.1 and Kir5.1 encoded by *KCNJ10* and *KCNJ16*, respectively, were found in 2 and 4 studies, respectively, in E1-E70 studies. Also, *KCNJ16* was detected as a DEG (less expression in glioblastomas versus oligodendrogliomas) in REMBRANDT, and with less expression in the 6 glioblastoma cell lines versus normal when studied by Patil, et al. [22]. Also, *KCNMB2* and *KCNJ16* are genes predicted to be negatively affected on chromosomes 3 and 17, respectively, in U87 glioblastoma cells, Supplemental Table 3 in Clark et al. [139]. Additionally, redistribution of K^+^ away from mitochondria within the cells’ interior may affect mitochondrial K^+^-dependent enzymes to influence malignant behavior. The alkaline pH of cancer cell cytoplasm in pH reversal activates ACLY which can lead to histone changes to possibly help explain the correlation of 50 studies in E1-E70 with KCN DEG that encode H^+^-sensitive proteins with 8 categories of histone/DNA modulators and 7 oncogenes (Figure 4A).

The E1-E70 study results only revealed gene expressions that crossed thresholds set by the authors of each study for significance (mostly *p* < 0.05) independently of any KCN consideration. Lower gene expression levels may be collectively biologically relevant in a network of KCN encoded proteins mediating potassium flux. Co-expression of non-KCN DEG (Figure 5) may become relevant as a signature of K^+^ dynamics if they occur in additional studies.

## SUMMARY

Migration and settling assays provided data to support and focus further experimental investigation of potassium flux in cancer cells. Although the reversed pH of cancer cells, generated via acid export by multiple extruders, is well established and of potential therapeutic value, the initiating events potentially involving potassium in providing intracellular protons to proton extruders could yield additional therapeutic targets. The multiple tumor models and range of study approaches used by many authors produced stochastic evidence for the large KCN potassium channel family playing a role in at least a subset of malignancies, with H^+^-sensitive members as potential downstream effectors of oncogenes. Although any future therapeutic suggestions regarding alterations of potassium channels carries concern regarding effects on the heart [291], the role for potassium channels as mediators of K^+^ flux that promotes K^+^/H^+^ exchange for proton transfer and subsequent efflux needs further exploration in cancer. The stochastic view of cancers based on unbiased studies detecting KCN genes incidentally in genomic landscapes suggests that encoded H^+^-sensitive KCN proteins may be downstream effectors of one or more drivers in cancer. Oncogenes may promote adaptation to adverse events and genetic damage to cells, including KCN genes, to optimize cell fitness for survival, possibly as a spectrum of ongoing changes that include the extreme outcome of becoming dedifferentiated, single migrating cells, via pH reversal. Also, cancer specific structural alterations of surface proteins encoded by genetically flawed KCN genes may add to the repertoire of targets for immune attack in personalized medical treatments. Author curated, published gene landscapes in complex cancer studies are useful for stochastically identifying candidates in the large KCN gene family that encode downstream effectors of oncogenes that may initiate pH reversal. They also offer cancer antigens to consider. Also, tumor models are identified in this manner where significant changes in KCN gene expression have been seen in malignancy.

## Data Availability Statement

Data generated in this study will be available upon request from the corresponding author. Please note that data from REMBRANDT per Adobe Flash (not available after 12/31/2020) can be seen in Figure 6 except for a few instances described which can also be made available on request. KCN gene data obtained from the published manuscript by Patil, V. et al. (22) follows the terms of their statement “an open access article distributed under the terms of the Creative Commons Attribution License, which permits unrestricted use, distribution, and reproduction in any medium, provided the original author and source are credited”. 34%, 31%, 13%, and 6% of the papers used in the E1-E70 study were found in Cancer Research, Cancer Cell, Nature Communications, and Journal of Molecular Diagnostics, respectively. Others are from 11 journals, 1 paper each. Papers for E1 – E70 are cited in Section *3.3.6. Categories Among Studies with E1-E70 Citations Per Category*.

## Acknowledgements

Ian F. Pollack, MD, University of Pittsburgh, Children’s Hospital of Pittsburgh, provided laboratory space, supplies, technicians, and glioblastoma cell lines, George K. Michalopoulos, MD, PhD, Department of Pathology, University of Pittsburgh, provided information on hepatocyte growth factor and MET. The Departments of Pathology and Neurological Surgery, University of Pittsburgh, provided institutional support for cell settling and migration experimental work. The 1301 authors of 70 papers (E1-E70) are thanked for outstanding contributions to the medical science literature used for gathering stochastic information on the KCN gene family while at Kent State, Kent, OH. I thank the Brain Health Research Institute of Kent State University, Kent, Ohio, as a current member, for technical advice and ongoing academic research activities.

## Author Contributions

The author performed the cell settling and migration assays, and the immunoblot at a previous institution, The University of Pittsburgh, Pittsburgh, PA. The author performed analysis of REMBRANDT, collection and analysis of 70 studies detecting KCN DEG, etc. and all other aspects of the study (text, figures, etc.) while associated with Kent State University, Kent, OH. Consultation for statistical advice and the use of EndNote was obtained and appreciated but the offered acknowledgement for the advice was not desired.

## Abbreviations

**The following abbreviations are used in this manuscript.**

ACLY: ATP Citrate Lyase
Arg1: Arginase 1
AS: Anti-seizure
BK: Big K (encoded by *KCNMA1)*
BSA: Bovine serum albumin
CD: Cluster Differentiation
*CLCN*: Chloride Channels gene family, *CLCN1,2,3*, etc.
COL: Collagen
CSM: Cancer Specific Mutation
Cx: Connexin
DEG: Differentially expressed gene(s)
E: Expression
EAAT: Excitatory Amino Acid Transporter
EAG1: Ether-A-Go-Go-K^+^ Channel Protein 1
ELK1,2: EAG Like K^+^ Channel Proteins 1,2
EMT: Epithelial Mesenchymal Transition
ERG1,3: EAG Related Gene K^+^ Channel Proteins 1,3
FBS: Fetal bovine serum
GABA: Gamma-aminobutyric acid
GBM: Glioblastoma
G-DOC: Georgetown Database of Cancer
Girk3,4: G protein-activated Inward Rectifier K^+^Channel Proteins, 3,4
GPR: G-protein Coupled Receptor
G-proteins: Guanosine-triphosphate-binding proteins
GWATERW: Grotthuss water wire
HGF: Hepatocyte growth factor
His/Chr/Epi: Histone/Chromatin/Epigenetic
HKG: House-keeping gene
IK: Intermediate K (encoded by *KCNN4*)
Indels: Insertions and Deletions
ITAG2: Integrin 2
ITG: Integrin
K2P: Tandem 2 pore domain KCN subfamily proteins
Kca: Calcium stimulated KCN subfamily proteins
Kchip1,3: Kv channel interacting proteins 1,3
KCNA: Subgroup of KCN genes, *KCNA1,2,3,etc.*
KCNAB: Subgroup of KCN genes, *KCNAB1,2,3*
KCNB: Subgroup of KCN genes, *KCNB1,2*
KCNC: Subgroup of KCN genes, *KCNC1,2,3,etc.*
KCND: Subgroup of KCN genes, *KCND1,2,3*
KCNE: Subgroup of KCN genes, *KCNE1,2,3, etc.*
KCNG: Subgroup of KCN genes, *KCNG1,2,3,etc.*
KCNH: Subgroup of KCN genes, *KCNH1,2,3, etc.*
KCNIP: Subgroup of KCN genes, *KCNIP1,2,3, etc.*
KCNJ: Subgroup of KCN genes, *KCNJ1,2,3, etc.*
KCNK: Subgroup of KCN genes, *KCNK1,2,3, etc.*
KCNM: Subgroup of KCN genes, *KCNM,A1,B1,B2, etc.*
KCNN: Subgroup of KCN genes, *KCNN1,2,3, etc.*
KCNQ: Subgroup of KCN genes, *KCNQ1,2,3, etc.*
KCNS: Subgroup of KCN genes, *KCNS1,2,3, etc.*
KCNT: Subgroup of KCN genes, *KCNT1,2*
Kir: Inwardly directing KCN subfamily proteins
Kv: Voltage dependent KCN subfamily proteins
LDH: Lactate dehydrogenase
LIF: Leukemia Inhibitory Factor
MCT: Monocarboxylate transporter
MEM: Minimal Essential Media
Mes: Mesenchymal
MiRP: MinK (minimal K^+^channel protein related peptide)
Na/K ATPase: Sodium/potassium ATPase
NOX: NADPH oxidases
Oligo: Oligodendroglioma
OMG: Oligodendroglioma Minus Glioblastoma
OMIM: Online Mendelian Inheritance in Man
PBS: Phosphate buffered saline
pHe: Extracellular pH
pHi: Intracellular pH
PIP2: Phosphatidyl inositol 4,5 bisphosphate
R: Pearson product-moment correlation
REMBRANDT: Repository of Molecular Brain Neoplasia Data
RGS: Regulator of G protein signaling
ROS: Reactive oxygen species
RRID: Research Resource Identifier
SERPIN: Serine proteinase inhibitors
SNV: Single Nucleotide Variant
SK: Small K (encoded by some *KCNN* genes)
Slack: Sequence like a calcium -activated K^+^ channel protein
SLC: Solute Carrier
Slick: Sequence like intermediate Conductance K^+^ channel protein
SOX: Sry-related HMG BOX
TasK1,2,5: Twik-related Acid Sensitive K^+^ channel proteins 1,2,5
Thik2: Tandem pore domain Halothane Inhibited K^+^ channel protein 2
TMEM: Transmembrane
TNFRSF: Tumor Necrosis Factor Receptor Super Family
Trek1: Twik-Related K^+^ channel protein 1
TwiK1,2: Tandem pore domain in Weak Inward Rectifying K^+^ Channel proteins 1,2
uPA: Urokinase-Type Plasminogen Activator

## Appendix A

**Figure A1.**
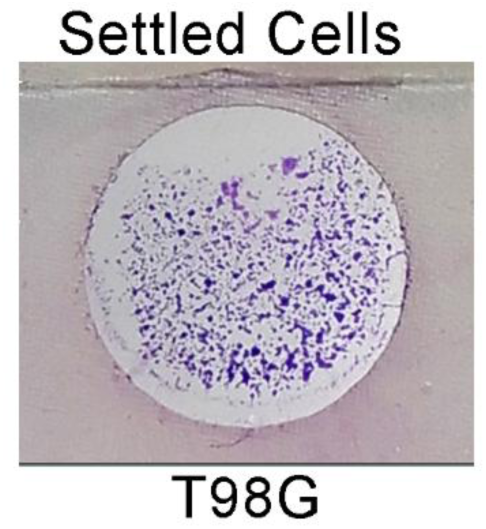
Macroscopic view of settled T98G glioblastoma cells on a Neuro Probe microscopic slide. One of 8 round 7 mm diameter areas that had been the bottom of a settling chamber well. Some cell clumping is noted. Diff Quik stain.

**Figure A2.**
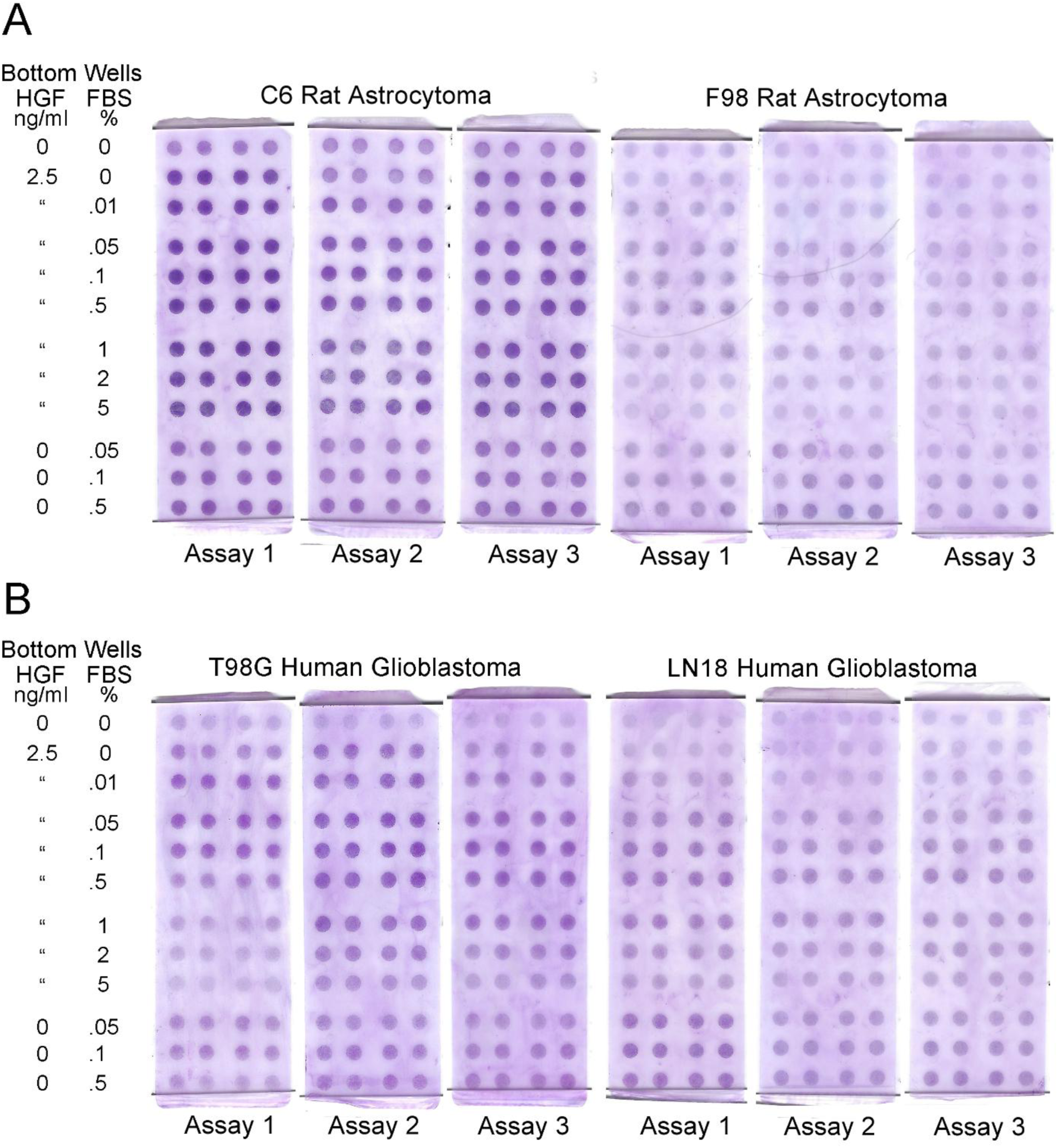
All three migration assays for each cell line (C6, F98, T98G, and LN18) in studies of HGF and FBS chemoattraction. One assay per cell line is shown in Figure 2A. Portions of assays that were digitized and analyzed are shown in Figure 2B. Text in Section 3.2. *Cell Migration with HGF and Serum Chemoattraction* and the Figure 2 legend provide additional information. Each circle of stained migrated cells is 3 mm in diameter.

**Figure A3.**
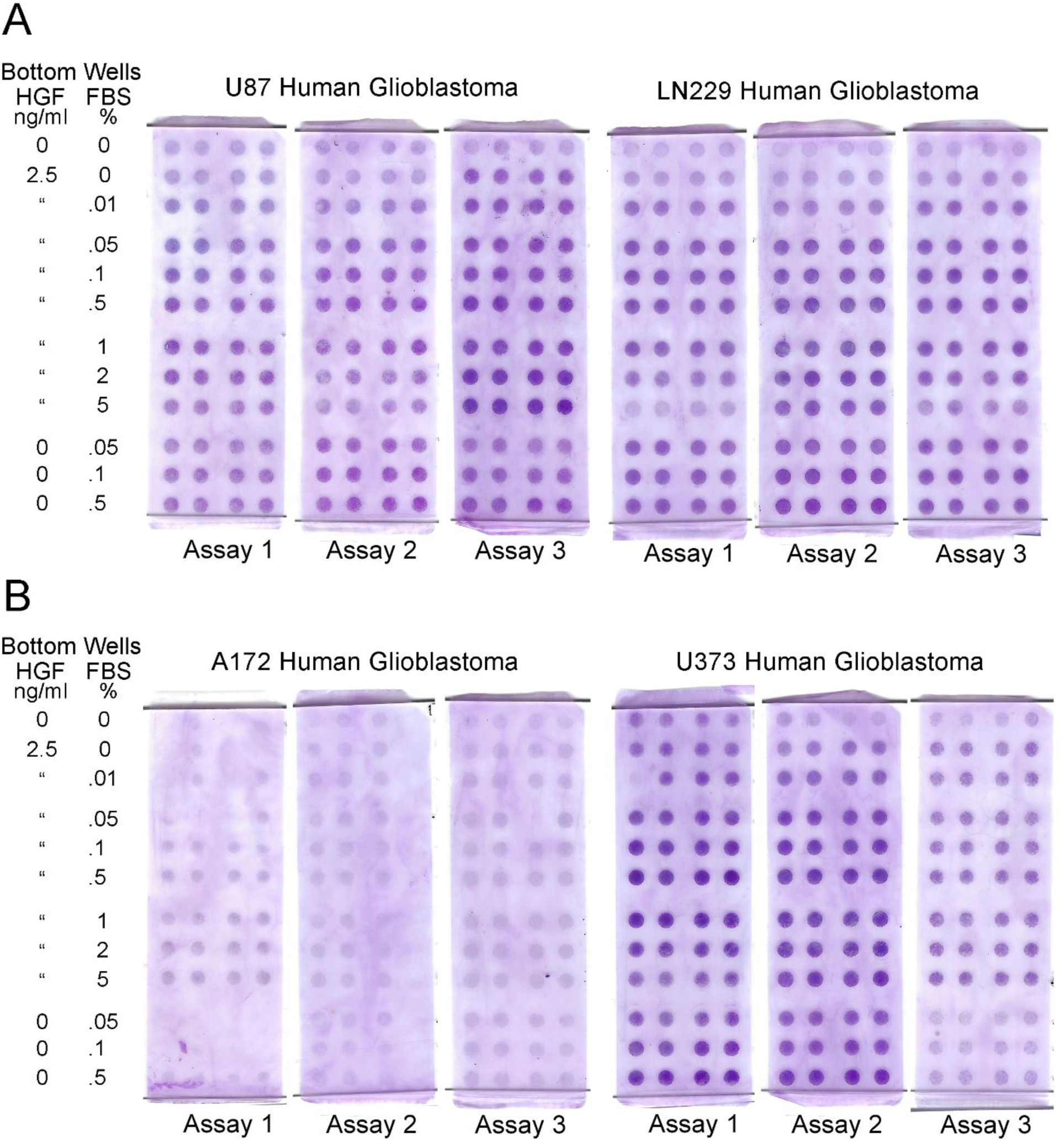
All three migration assays for each cell line (U87, LN229, F98, A172, and U373) in studies of HGF and FBS chemoattraction. One assay per cell line is shown in Figure 2A. Portions of assays that were digitized and analyzed are shown in Figure 2B. Text in Section 3.2. *Cell Migration with HGF and Serum Chemoattraction* and the Figure 2 legend provide additional information. Each circle of stained migrated cells is 3 mm in diameter.

**Figure A4.**
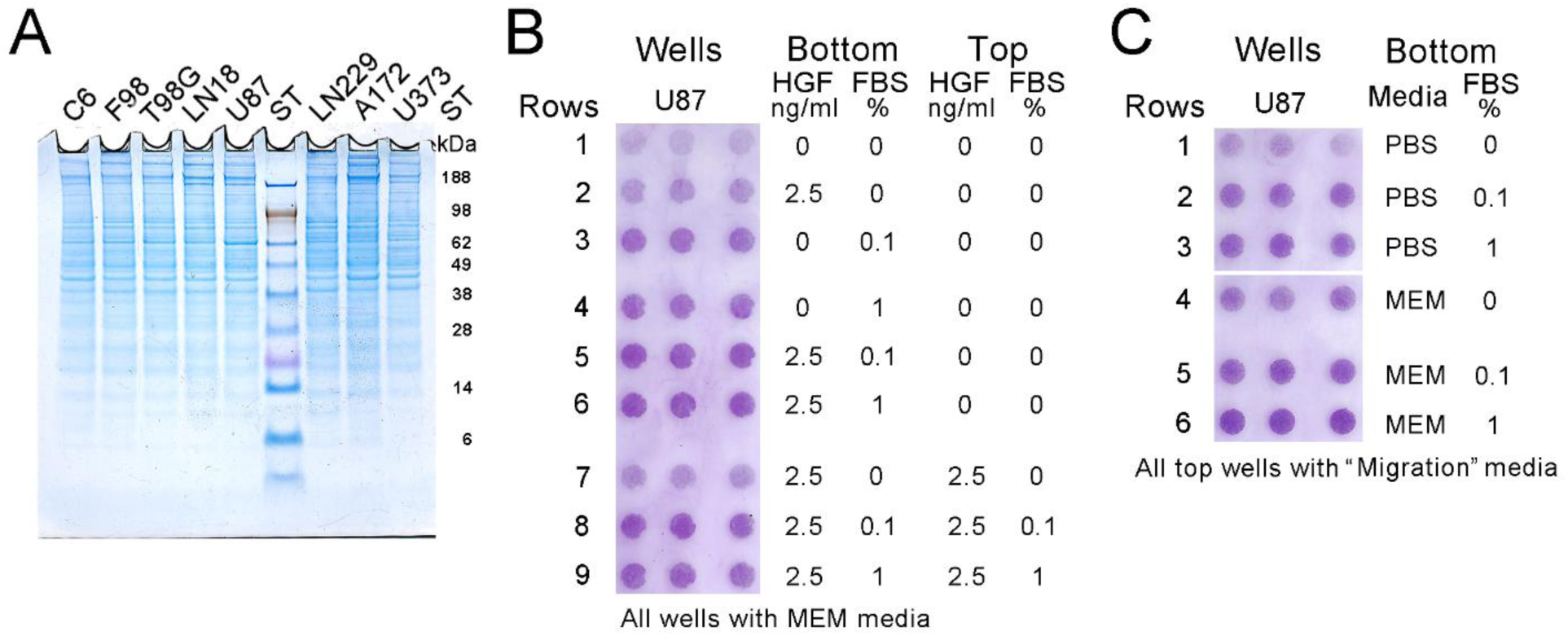
Additional images for Figure 2. (A) Gel image for the immunoblot in Figure 2C showing equal loading of protein in each lane for all 8 cell lines. (B) Scanned image of the U87 migration assay shown in Figure 2D. Chemoattraction of HGF and 2 concentrations of FBS and chemokinesis of HGF, with and without 2 concentrations of FBS, were analyzed in U87 cells. (C) Scanned image of the U87 migration assay shown in Figure 2E. Dulbecco’s PBS and MEM were tested as media in the lower wells for migration without and with 2 concentrations of FBS present. The image of the filter removed (white horizontal line) an unrelated portion of the assay that tested a different batch of FBS for suitability in future migration assays. Text in Section 3.2. *Cell Migration with HGF and Serum Chemoattraction* and the Figure 2 legend provide additional information. Each circle of stained, migrated cells is 3 mm in diameter.

## Appendix B

**Table B1.**
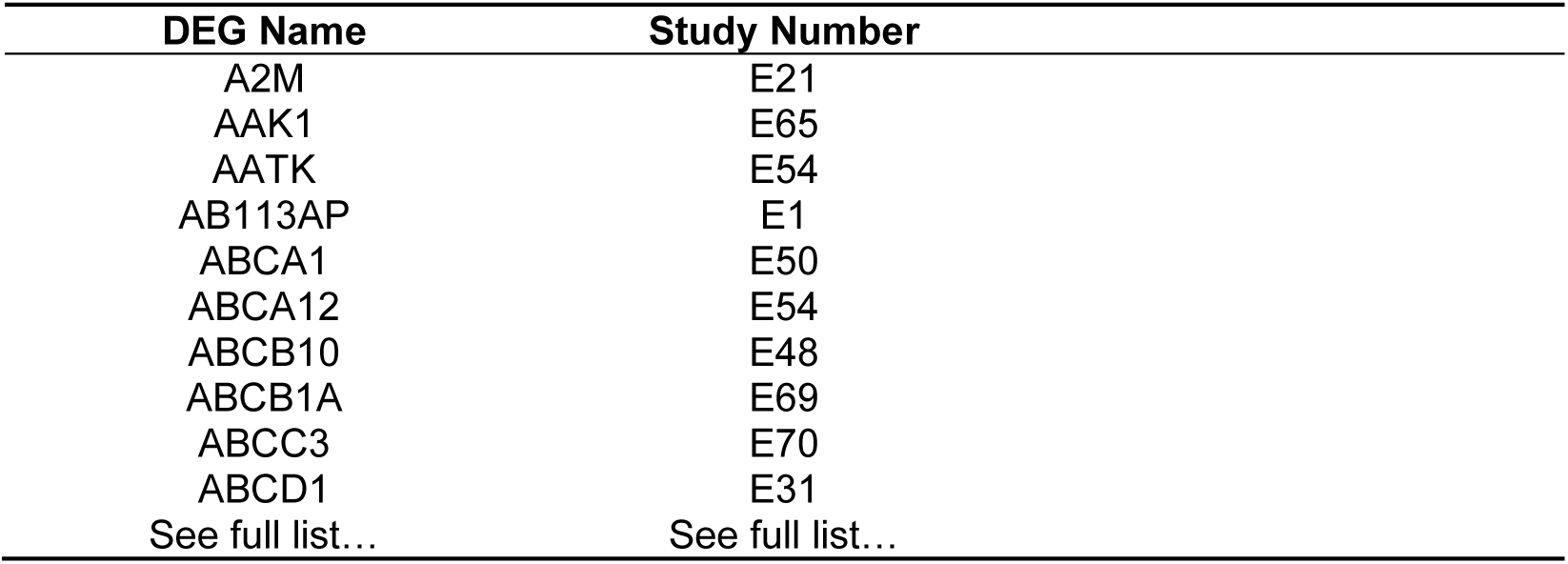
A sample of alphabetized differentially expressed genes (DEG) in E1-E70 studies. See AllE1E70DEG3239Total for the entire list of DEG in a table format after the References and Disclaimer.

| DEG Name | Study Number |
| --- | --- |
| A2M | E21 |
| AAK1 | E65 |
| AATK | E54 |
| AB113AP | E1 |
| ABCA1 | E50 |
| ABCA12 | E54 |
| ABCB10 | E48 |
| ABCB1A | E69 |
| ABCC3 | E70 |
| ABCD1 | E31 |
| See full list... | See full list... |

**Table B2.** KCN genes in E1-E70 studies that encode H^+^-sensitive proteins.

| **KCN Gene** | **Protein** | **Reference** |
| --- | --- | --- |
| *KCNA1* | Kv1.1 | [292] |
| *KCNA3* | Kv1.3 | [293,294] |
| *KCNA5* | Kv1.5 | [295-301] |
| *KCNAB2* | Kvbeta2 | [302,303] |
| *KCND3* | Kv4.3 | [304-306] |
| *KCNE2* | MiRP1 | [85,307-311] |
| *KCNE3* | MiRP2, R83H variant | [253] |
| *KCNH1* | Kv10.1, EAG1 | [254,312] |
| *KCNH2* | Kv11.1, HERG1 | [254,313-325] |
| *KCNH3* | Kv12.2, ELK2 | [254] |
| *KCNH8* | Kv12.1, ELK3 | [254,326] |
| *KCNJ5* | Kir3.4, GIRK4 | [255,256,327] |
| *KCNJ8* | Kir6.1, KATP-1 | [327-333] |
| *KCNJ10* | Kir4.1, BIRK10 | [334-336] |
| *KCNJ10/KCNJ16* | Kir4.1/Kir5.1heterodimer | [72,326,337-343] |
| *KCNJ13* | Kir7.1 | [72,327,344] |
| *KCNK1* | K2P1, TWIK1 | [343,345-350] |
| *KCNK2* | K2P2, TREK1 | [343,348,351-357] |
| *KCNK3* | K2P3, TASK1 | [343,346,348,355,358-361] |
| *KCNK5* | K2P5, TASK2 | [237,343,346,348,362-369] |
| *KCNK6* | K2P6, TWIK2 | [346,370,371] |
| *KCNK15* | K2P15, TASK5 | [372] |
| *KCNMA1* | KCa1.1, BK, Slo1, maxiK | [96,373-388] |
| *KCNMB4* | BKbeta4 | [389-393] |
| *KCNN2* | KCa2.2, SK2 | [394] |
| *KCNN3* | KCa2.3, SK3 | [394] |
| *KCNN4* | KCa3.1, SK4, Gardos | [394-397] |
| *KCNQ1* | Kv7.1, KvLQT1 | [85,237,307-309,311,399-405] |
| *KCNQ2* | Kv7.2 | [402(described data),406,407] |
| *KCNQ3* | Kv7.3 | [406] |
| *KCNQ5* | Kv7.5 | [408] |
| *KCNT1* | KCa4.1, Slo2.2, KNa1.1 | [409-412] |
| *KCNU1* | KCa5.1, Slo3, KCNMA3 | [389,412-418] |

**Figure B1.**
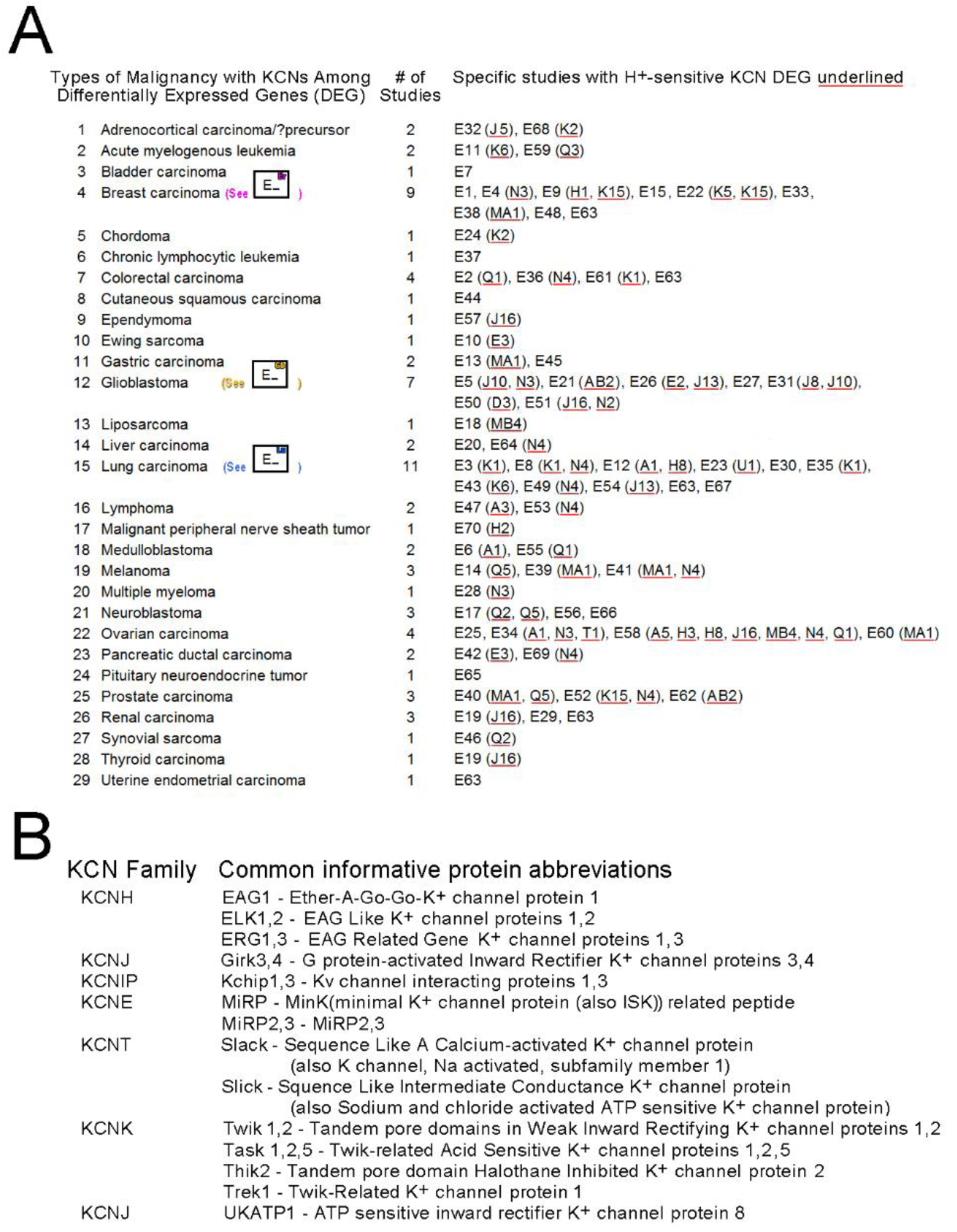
Types of malignancies studied in E1-E70 and informative protein abbreviations for proteins in the KCN family are listed. **(A)** Specific studies are listed for each type of 29 malignancies with their detected KCN DEG. Red underlining indicates the KCN DEG that encode H^+^-sensitive proteins. Boxes that correspond with studies of brain, breast, and lung malignancies are color coded, yellow, pink, and blue, respectively. These graphics are also used in the histograms shown in Figures 3 and 4. (**B**) Some of the common protein abbreviations for KCN family members in Figure 3 indicating functional information not explained in the text are included.

## Text B1. Potential Interactions of KCN Proteins with H^+^-Sensitive G Protein-Coupled Receptors.

Multiple KCN proteins have interactions with four G protein-coupled receptors encoded by *GPR4*, *65*, *68*, and *151* that are among 6 reported to be H^+^-sensitive [419–425]. H^+^-sensitive proteins encoded by GPR genes are found in many types of tumors [423,424,426,427]. Functional complementation between proteins encoded by *GPR4* and *KCNK5*/TASK2 [427,428,429] and related expression profiles of proteins encoded by *GPR31*, G*PR151*, *KCNK3*/TASK1, and *KCNK9*/TASK3 have been found in tumors [425]. STRING interactions (Gene Cards, 10/17/2024) of proteins encoded by *GPR4*, *65*, *68*, and*151* as a group included proteins encoded by 9 KCNs, *KCNJ8*, *K1, K2, K4, K5*, *K10*, *K16*, *K17*, and *K18*. Among these, 8 of the 70 studies (E3, 8, 22, 24, 31,35, 61, 68) detected *KCNJ8*, *K1, K2, K5* as DEG. Also, although *GPR39*’s encoded protein is not reported to be H^+^-sensitive, it interacts with the four H^+^-sensitive proteins encoded by *GPR4*, *65*, *68*, and *132*. *GPR39* is a DEG found in 2 of the 70 studies, E14 [194] and E60 [177], that also detected *KCNQ5* and *KCNMA1*, respectively. The ligand of the protein encoded by *GPR39* is encoded by *GPNMB* [430] and *GPNMB* was a DEG in 3 studies, E26 [213], E29 [158], and E49 [167]. *GPR35*’s protein (also not H^+^-sensitive) interacts with *GPR4*’s encoded H^+^-sensitive receptor protein. *GPR35* is a DEG in two studies that found *KCNG2* and *KCNE3*, in E20 [211] and E42 [218], respectively. The ligand for *GPR35*’s receptor protein is kynurenic acid [431]. *GPR84*’s protein (also not H^+^-sensitive) interacts with *GPR65*’s encoded H^+^-sensitive receptor protein. *GPR84* is a DEG from two studies that found *KCNK6* in E11 [174] and *KCNK12* and *KCNU1* in E23 [202]. The ligands for *GPR84*’s receptor protein are medium-chain fatty acids [432]. Thirteen of the14 studies (out of 70) with potential indirect GPR protein sensitivity to H^+^ also detected KCN DEG that encode H^+^-sensitive proteins. Potentially, KCN DEG encoded proteins in 13 (18.6%) of the E1-E70 studies have indirect involvement with H^+^-sensitive G-protein-coupled receptor proteins via STRING interactions (Gene Cards, 10/17/2024) or as described above according to the cited literature. They may function collaboratively with GPR encoded H^+^-sensitive proteins in tumors.

## Disclaimer

The statements and opinions are those of the author. The author disclaims responsibility for any injury to people or property resulting from any ideas, methods, instructions, products, or publications cited in the content.

---

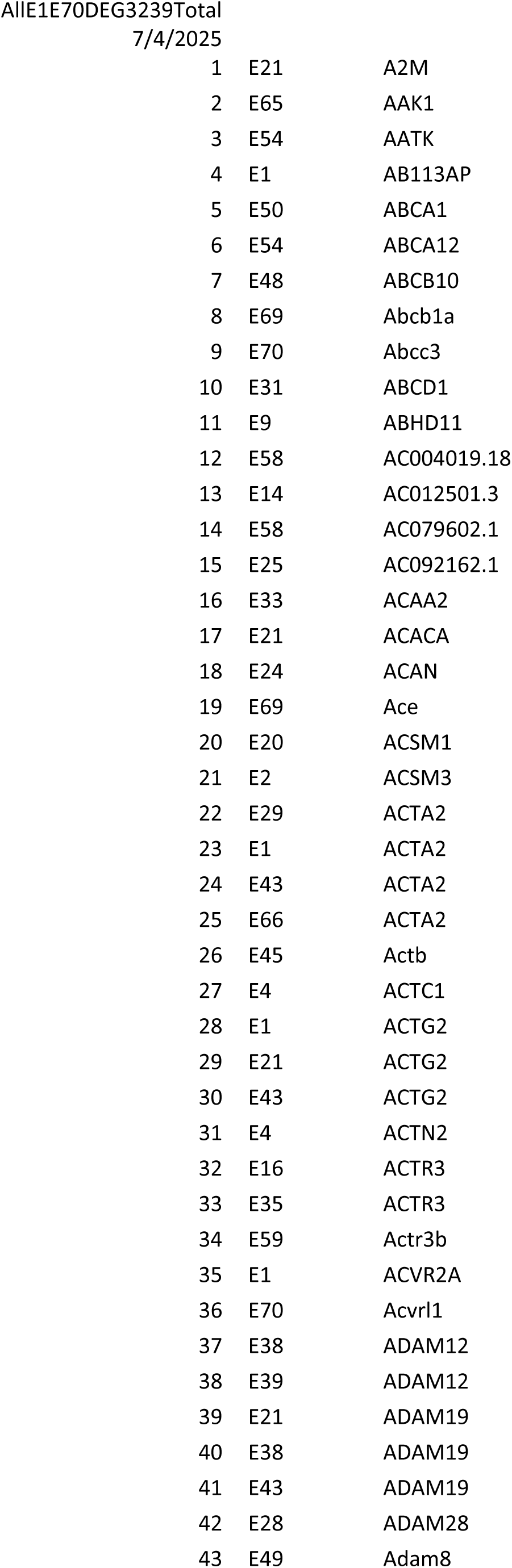
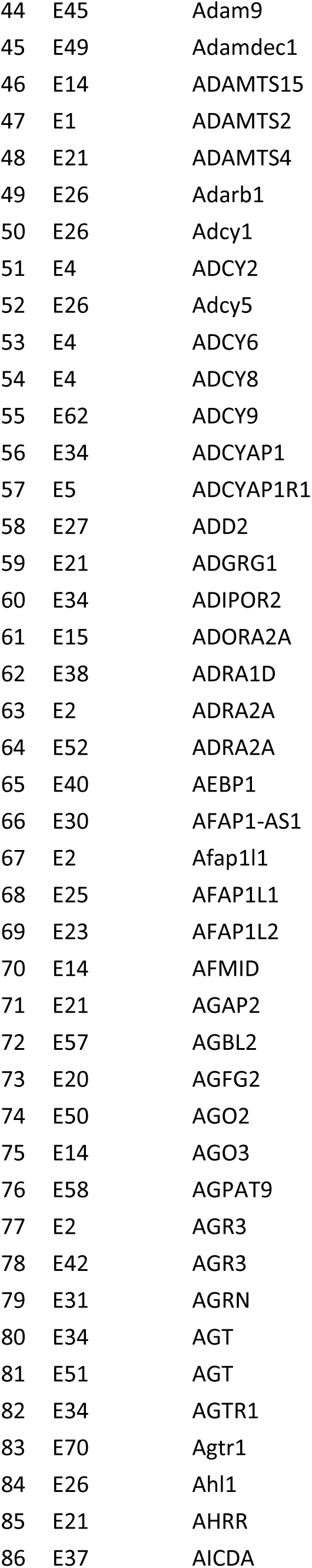
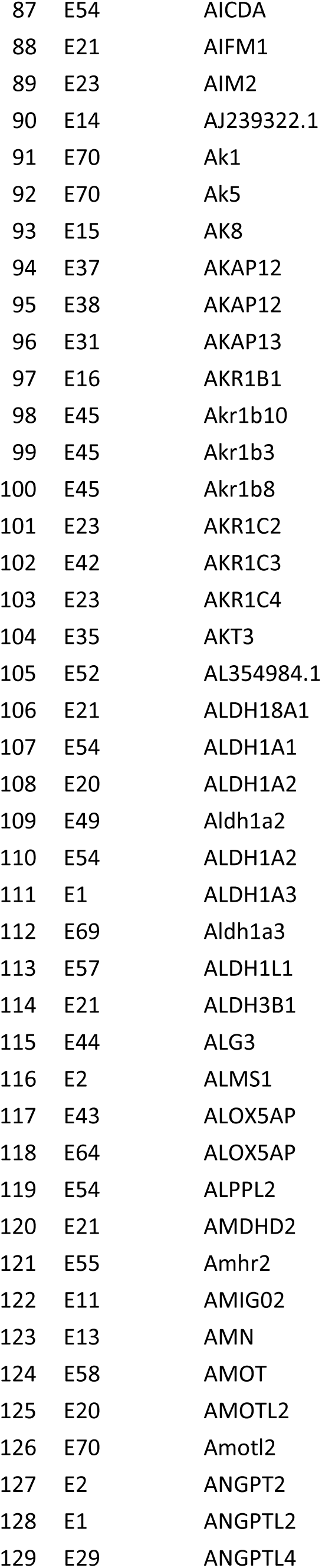
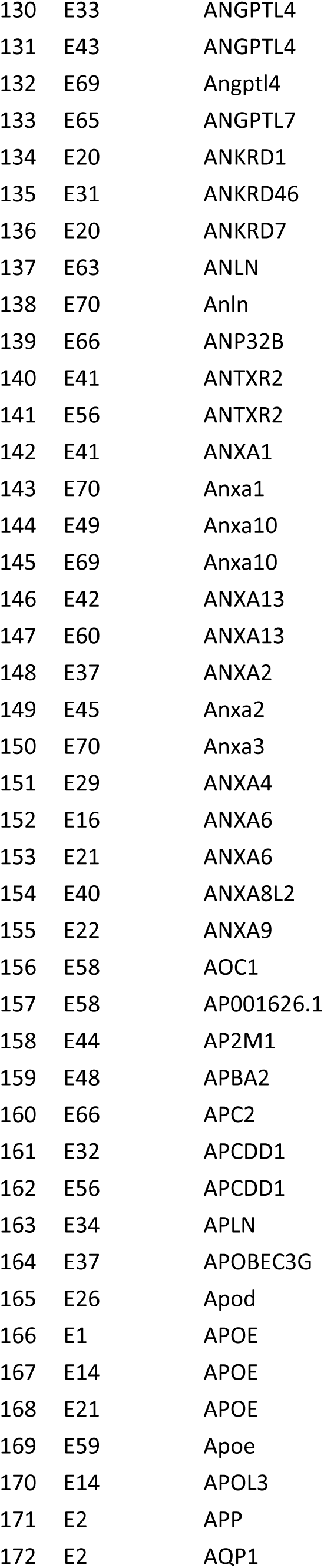
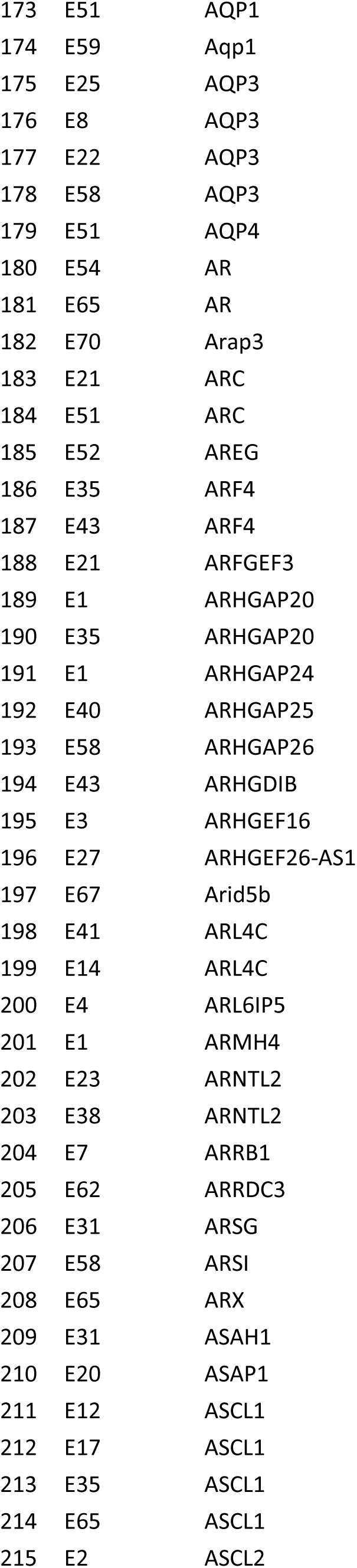
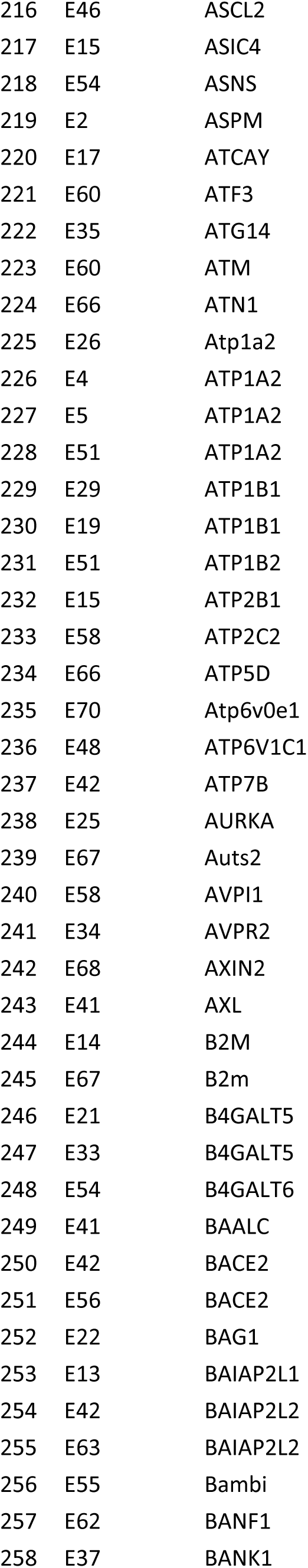
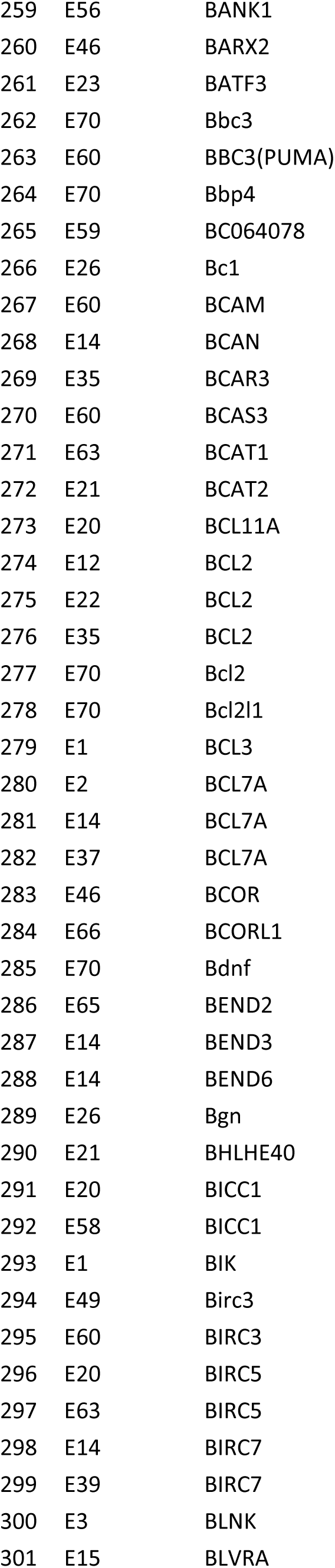
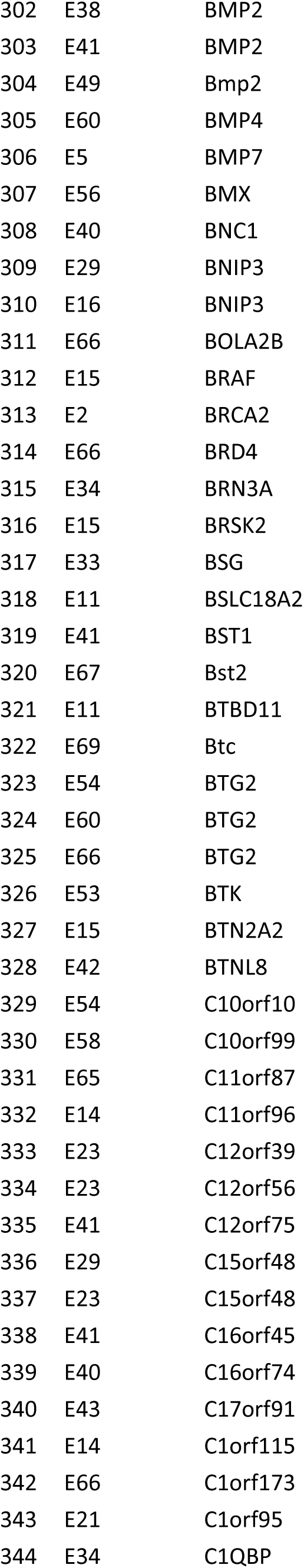
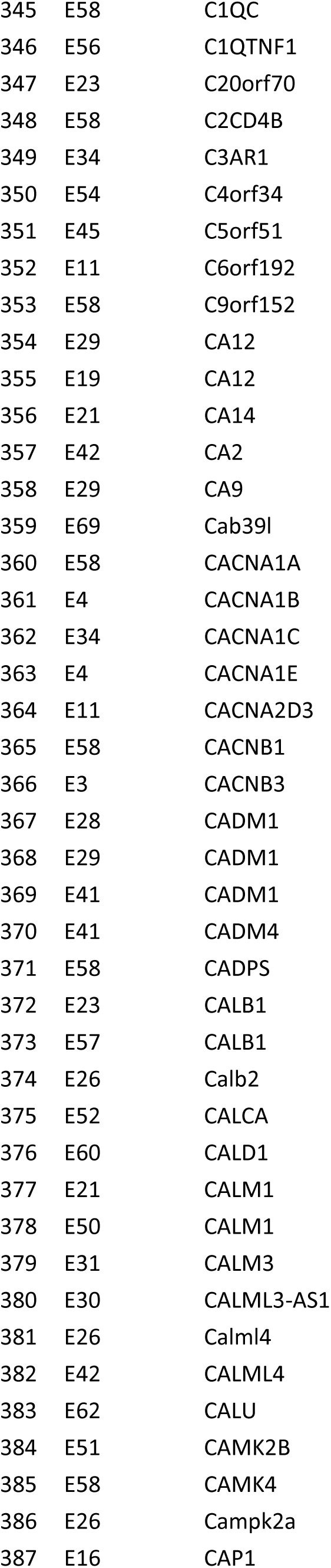
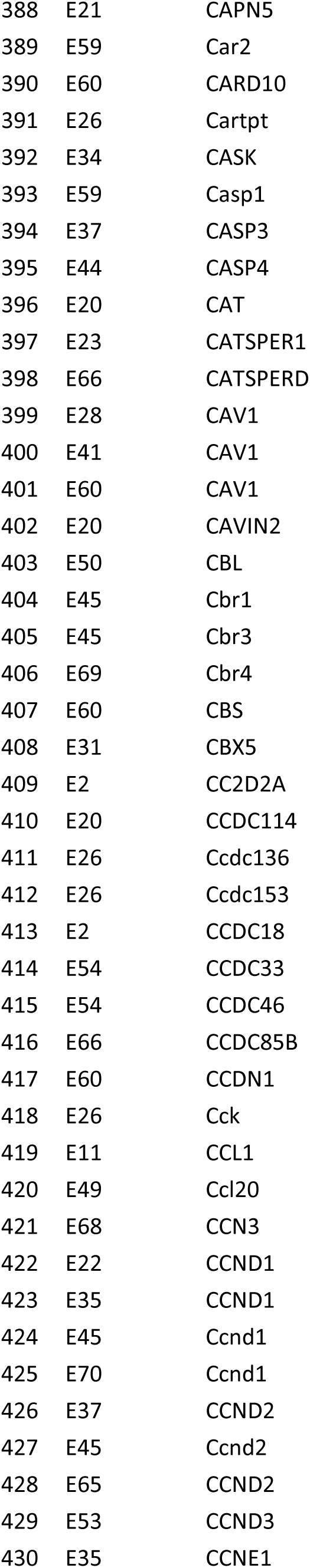
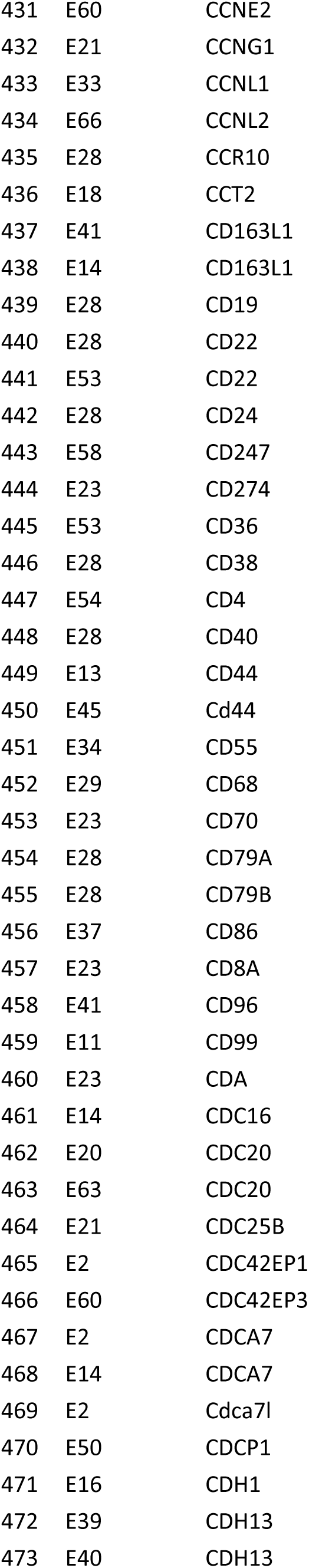
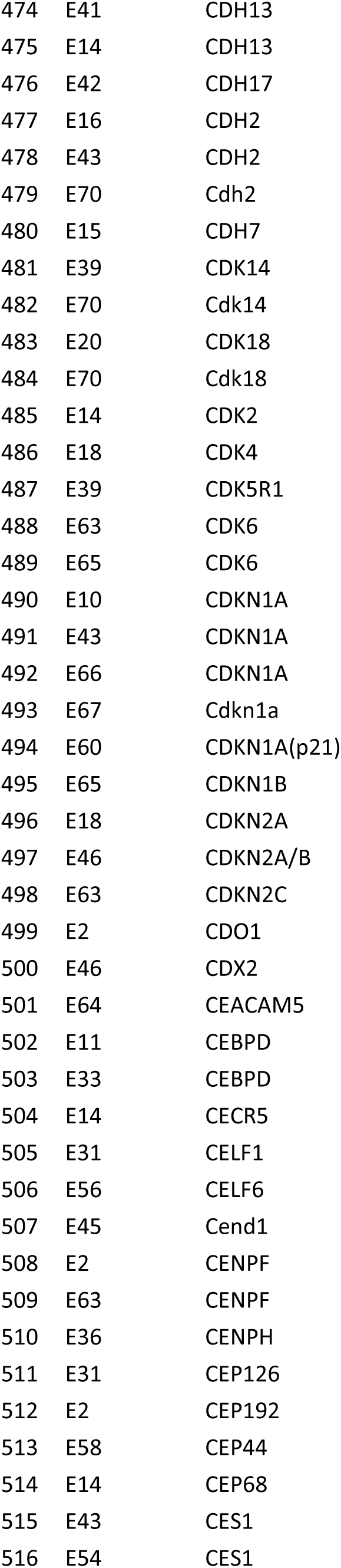
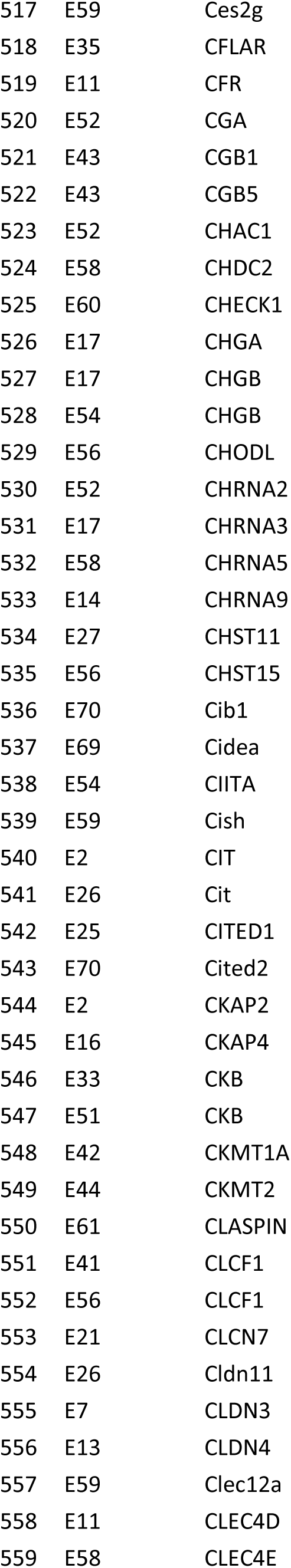
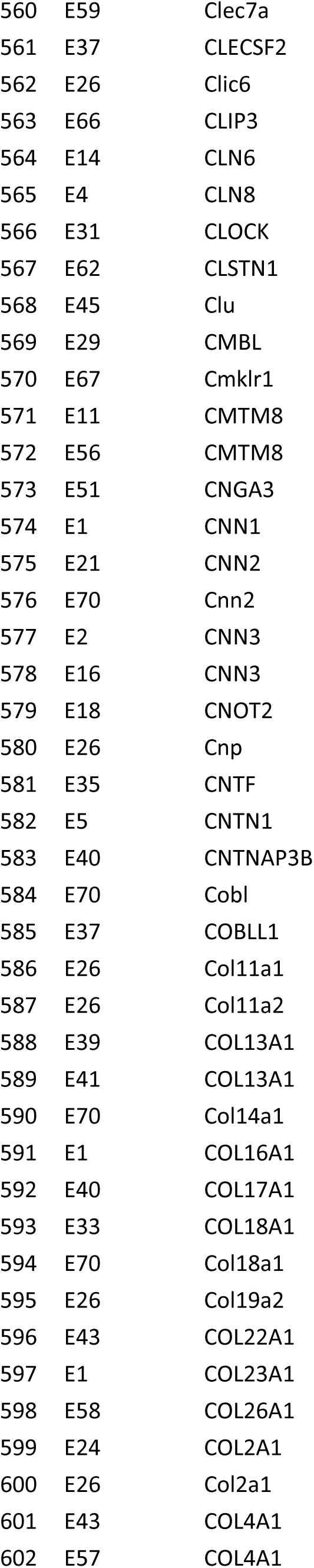
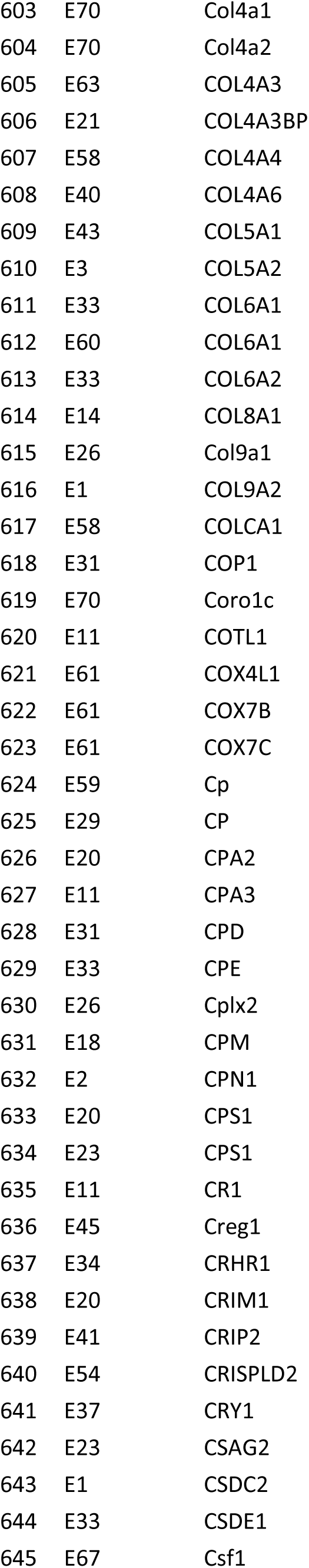
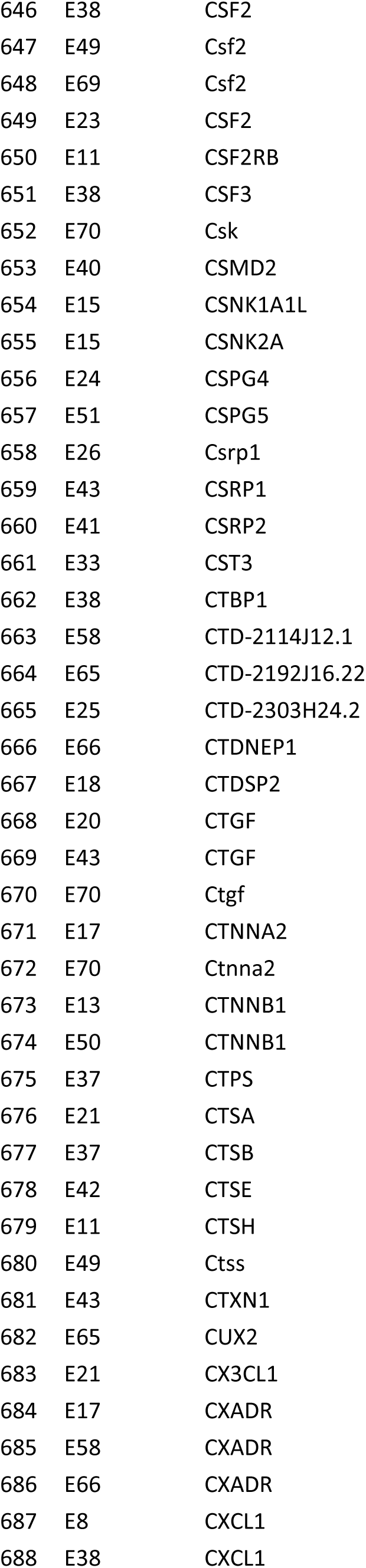
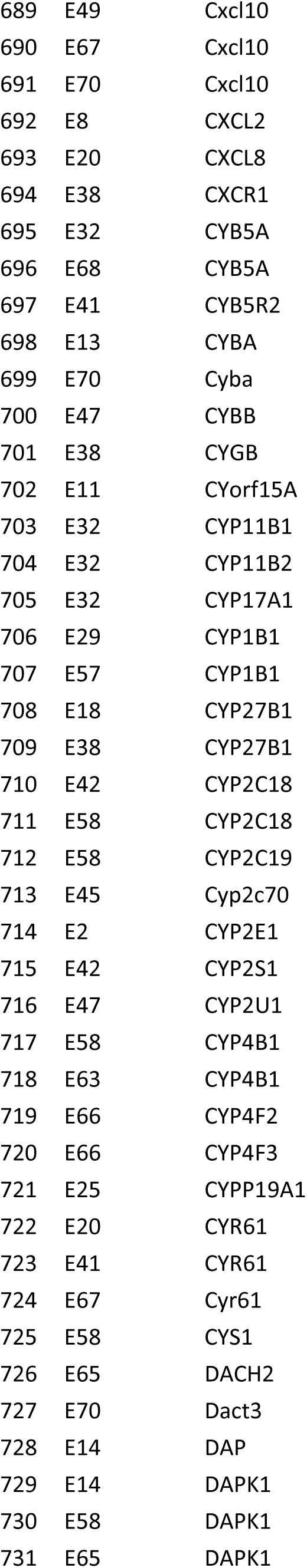
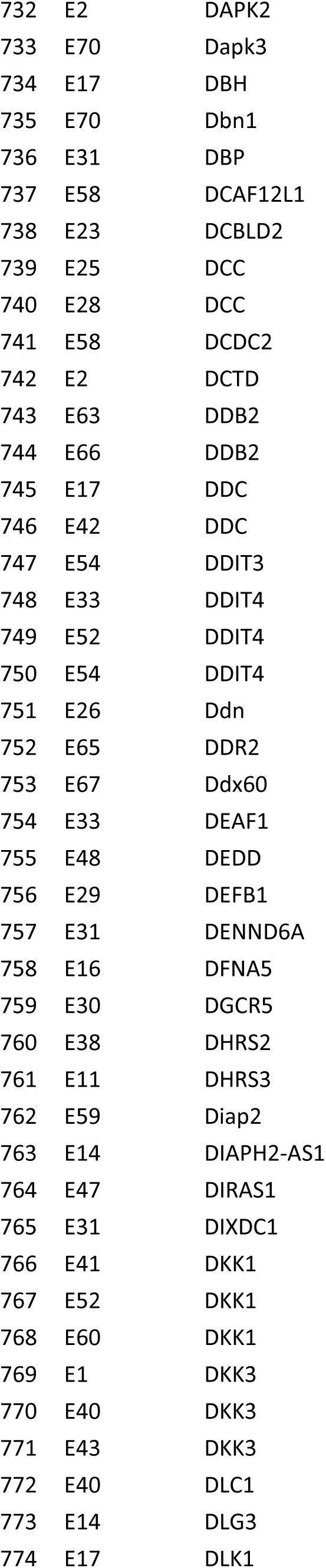
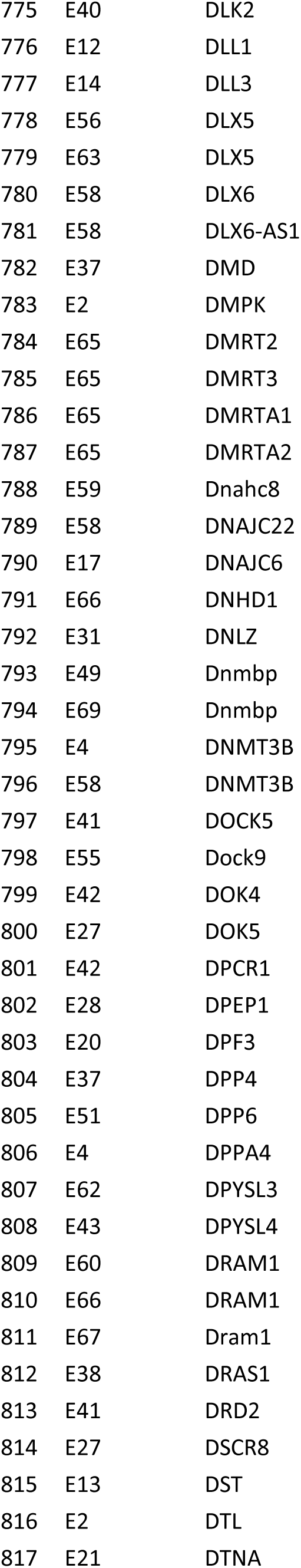
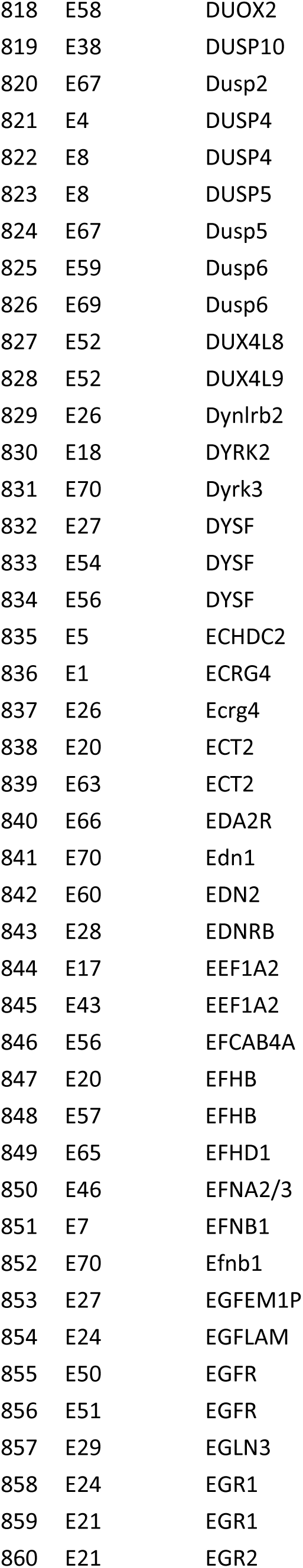
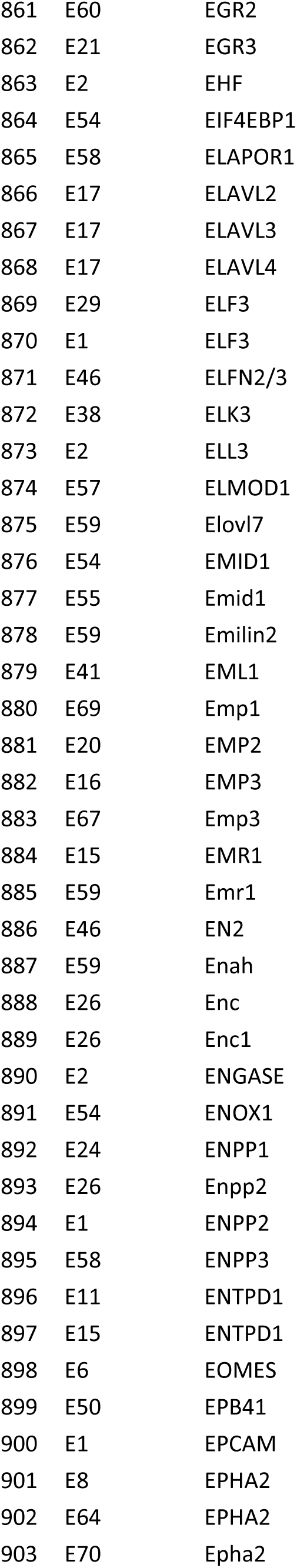
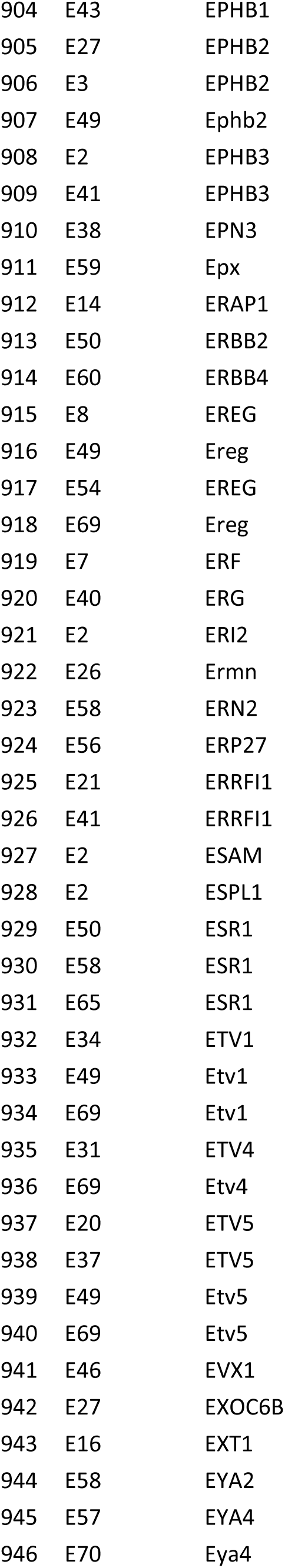
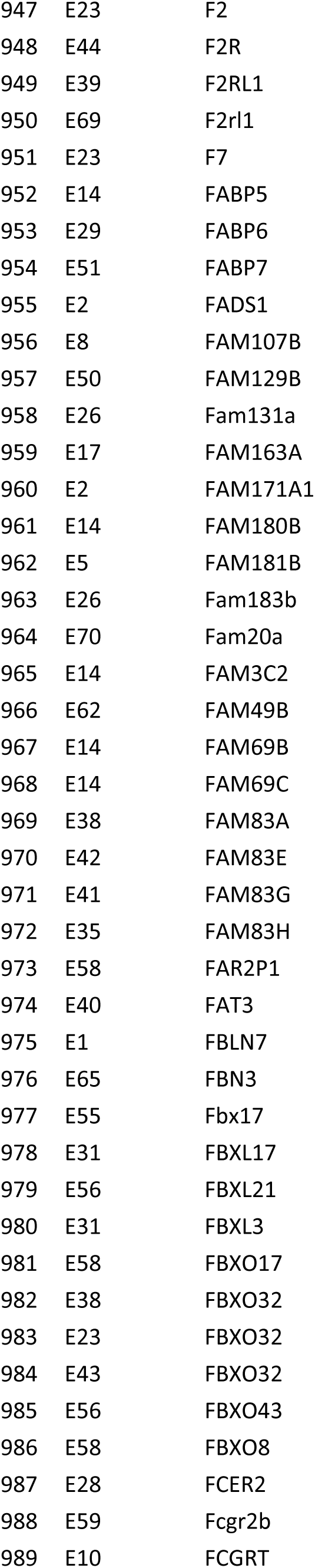
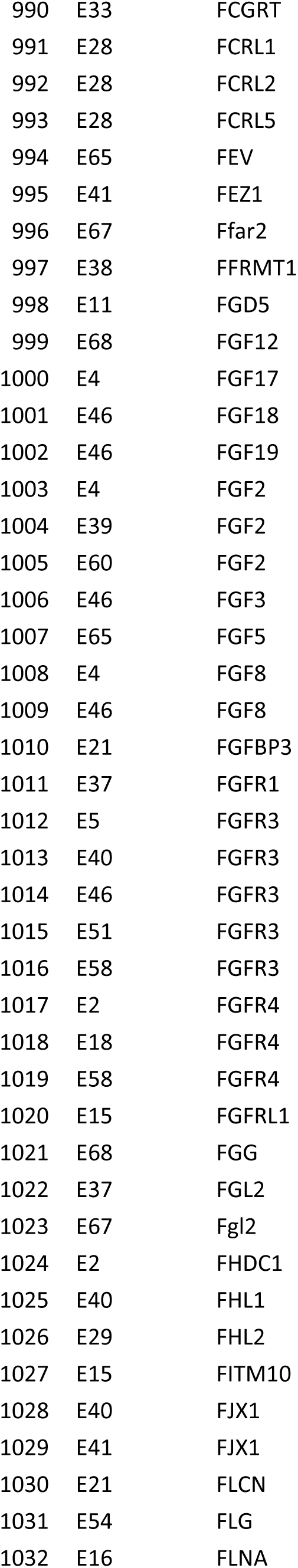
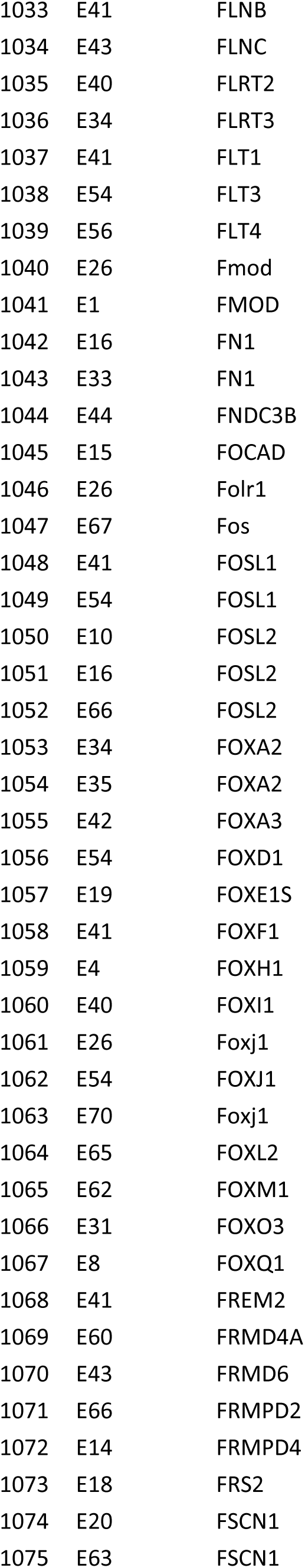
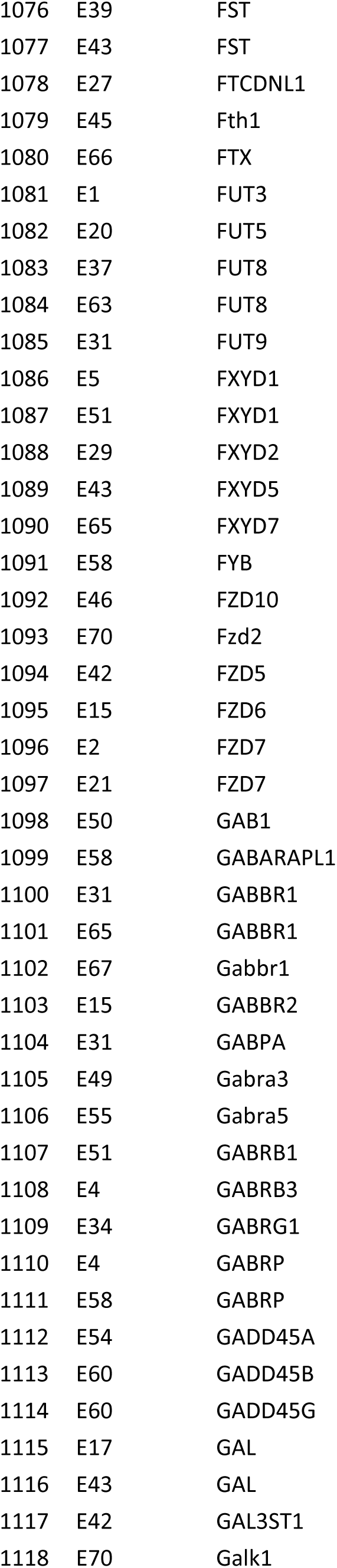
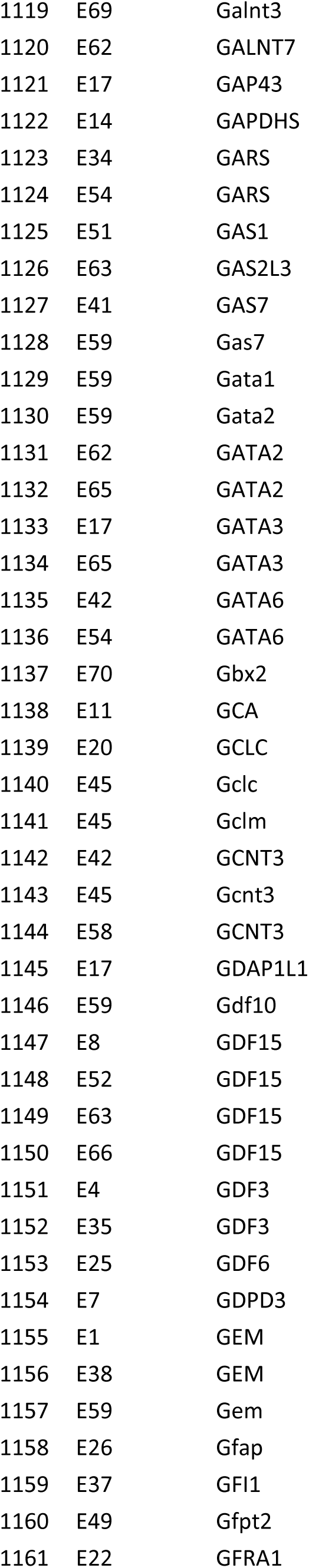
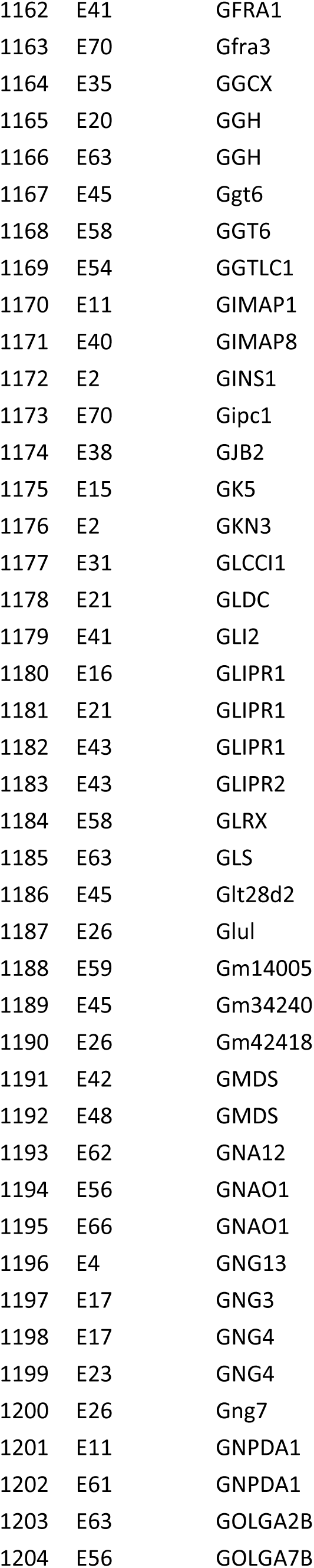
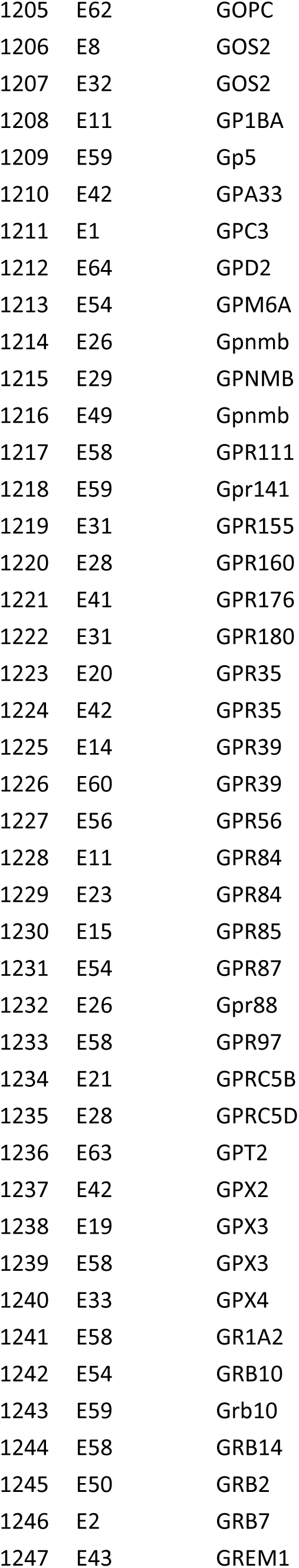
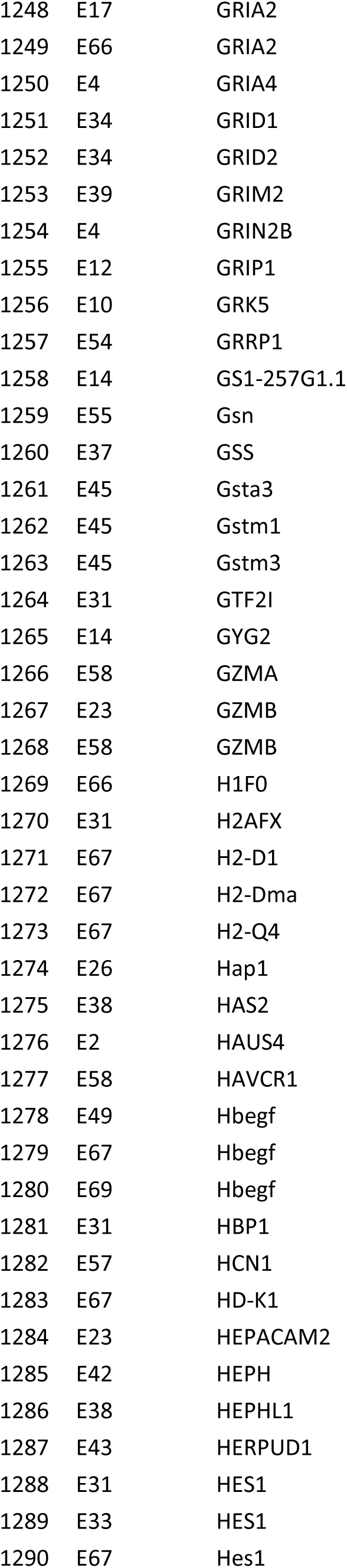
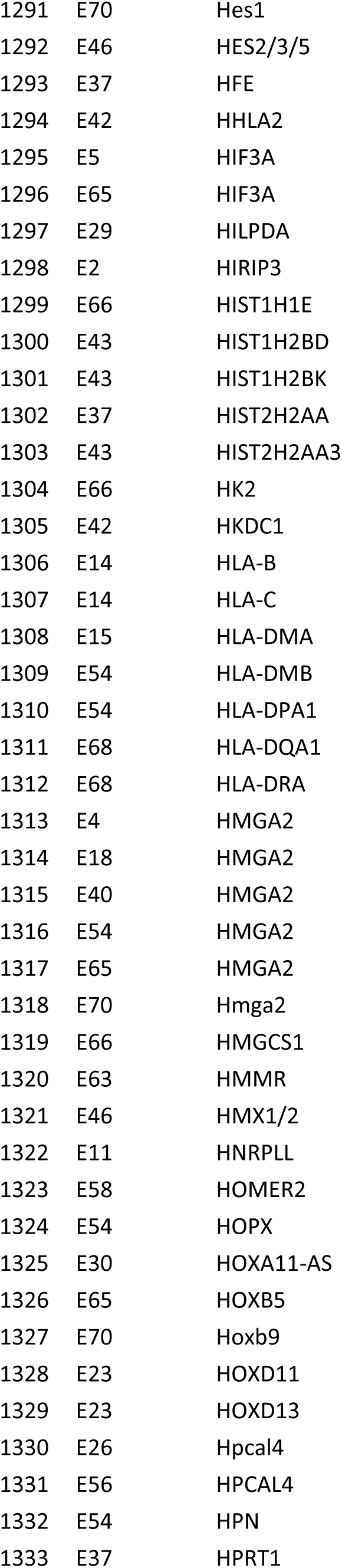
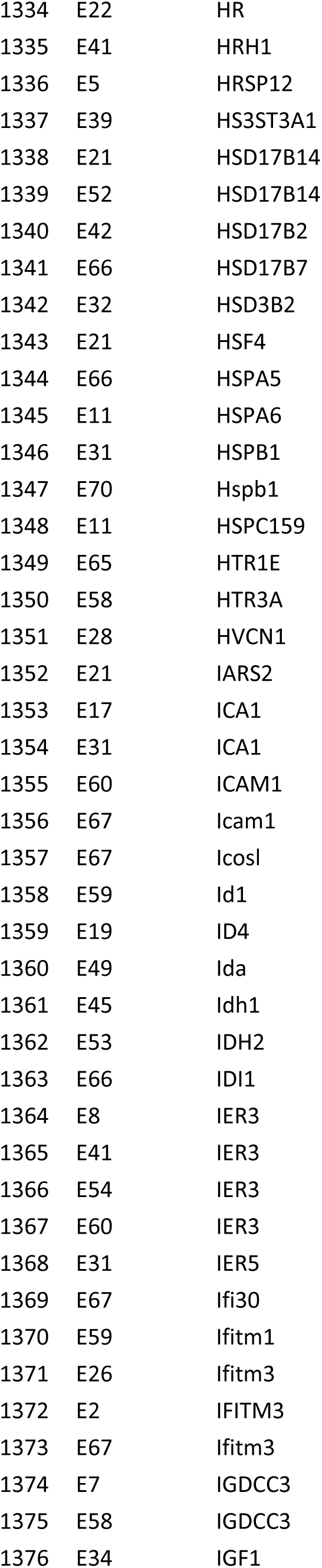
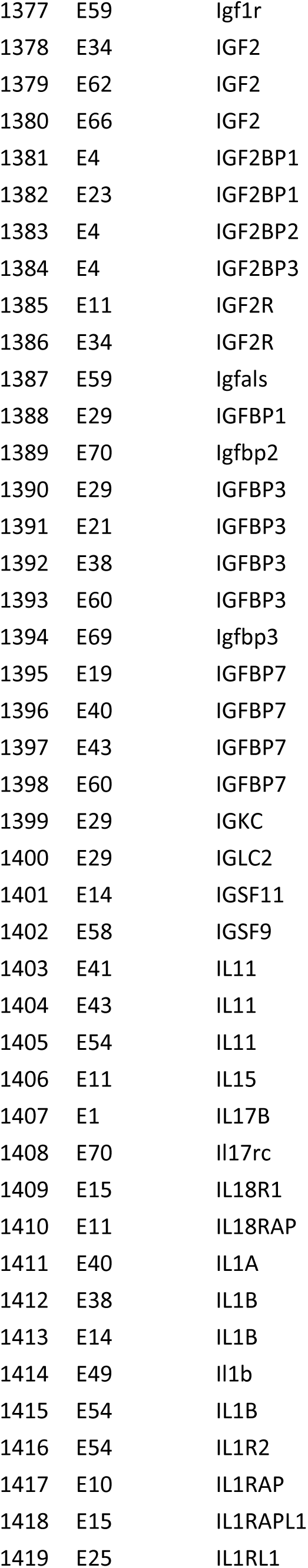
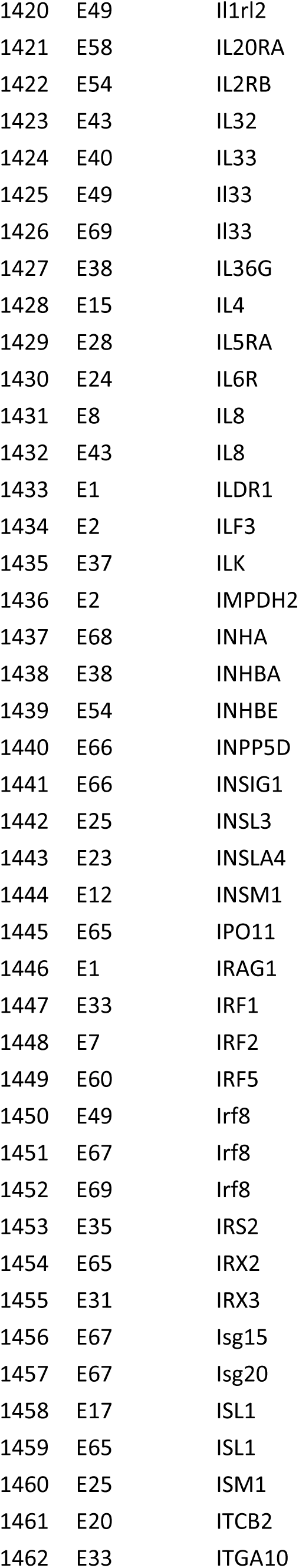
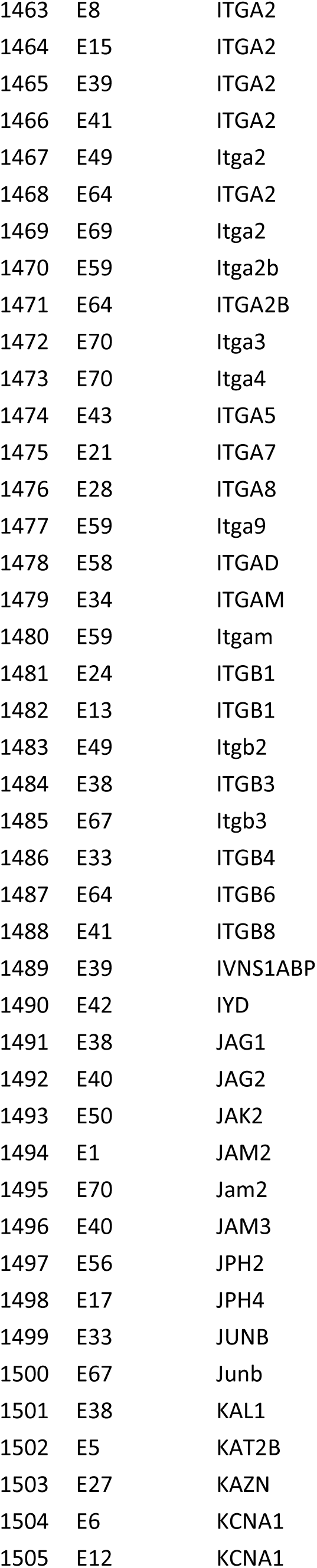
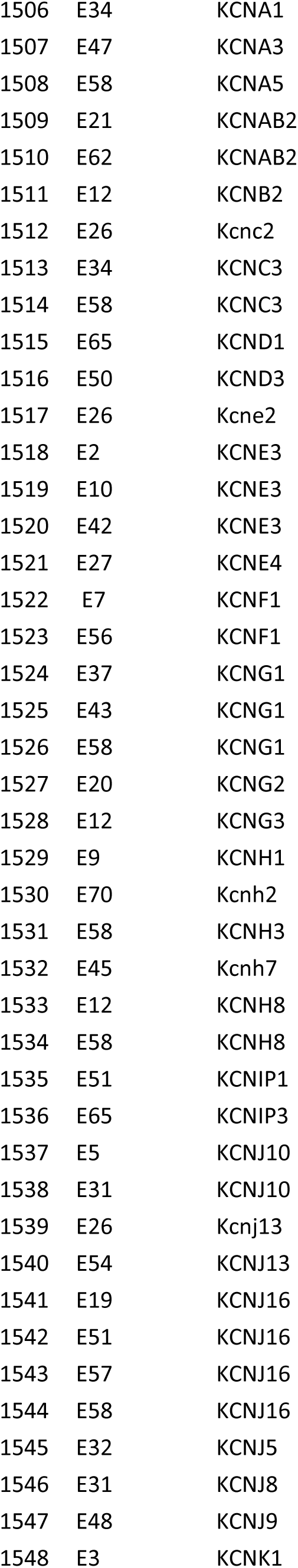
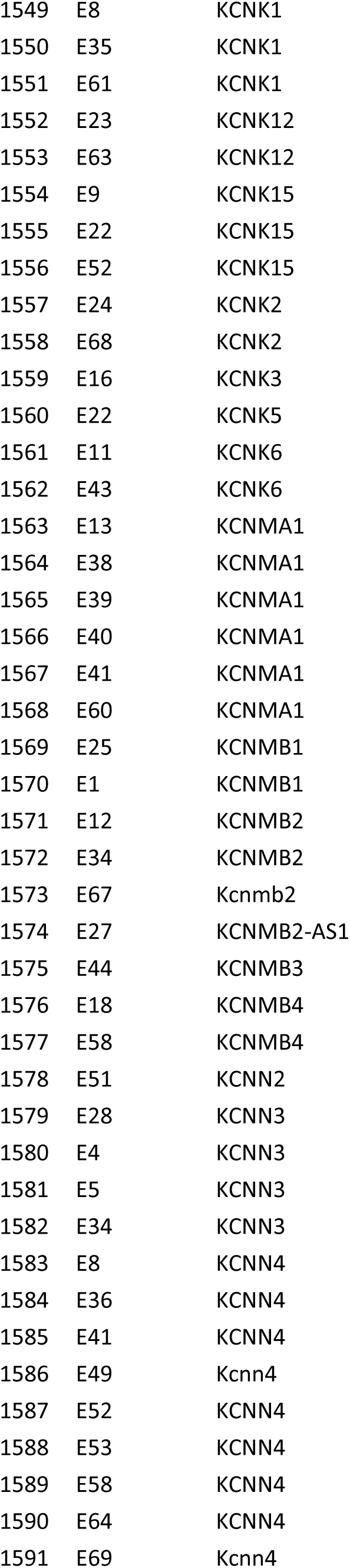
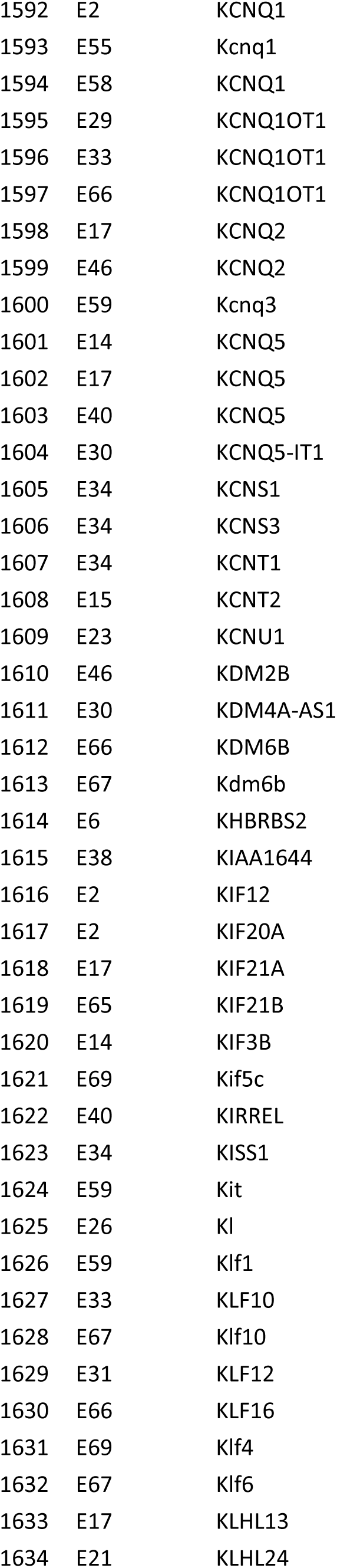
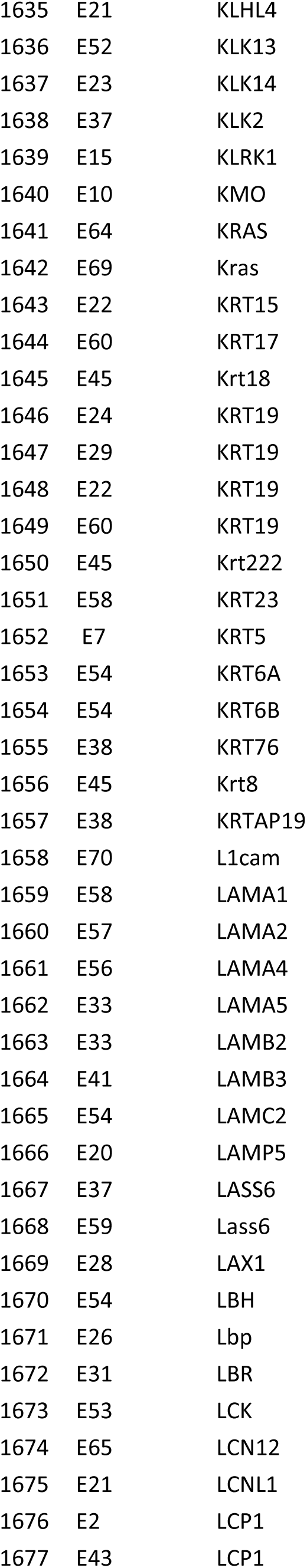
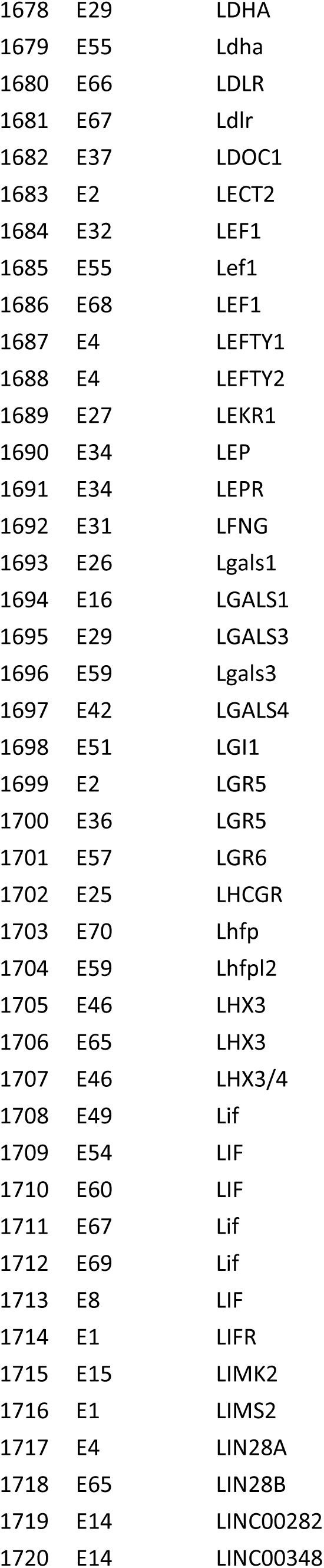
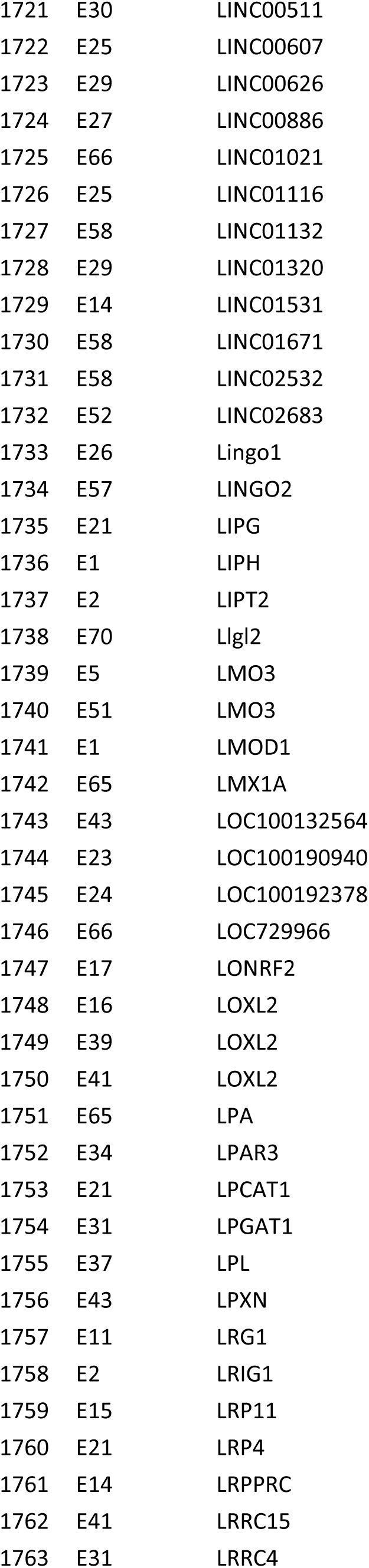
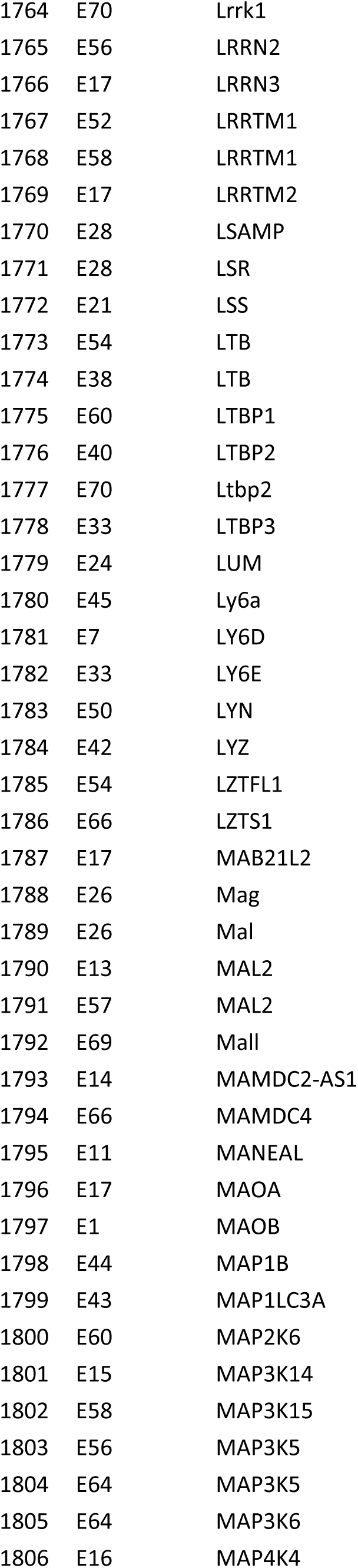
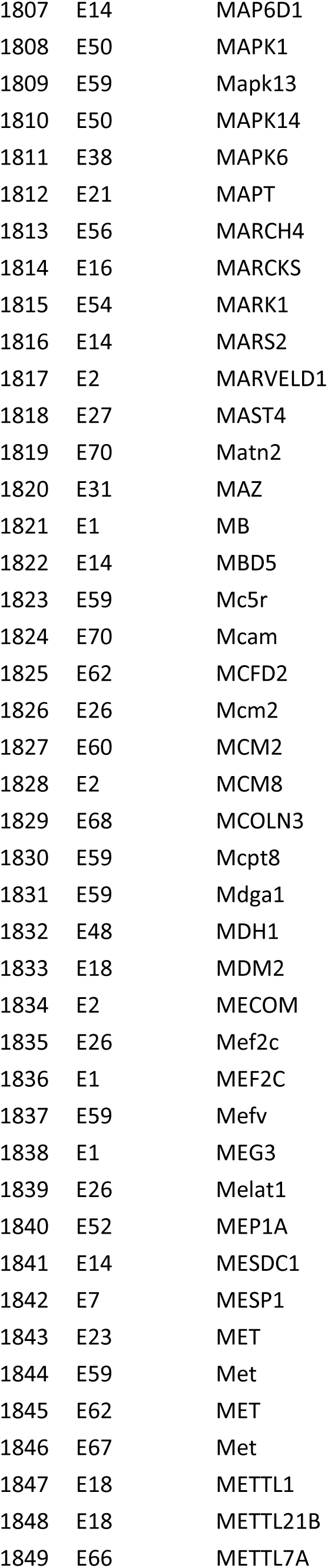
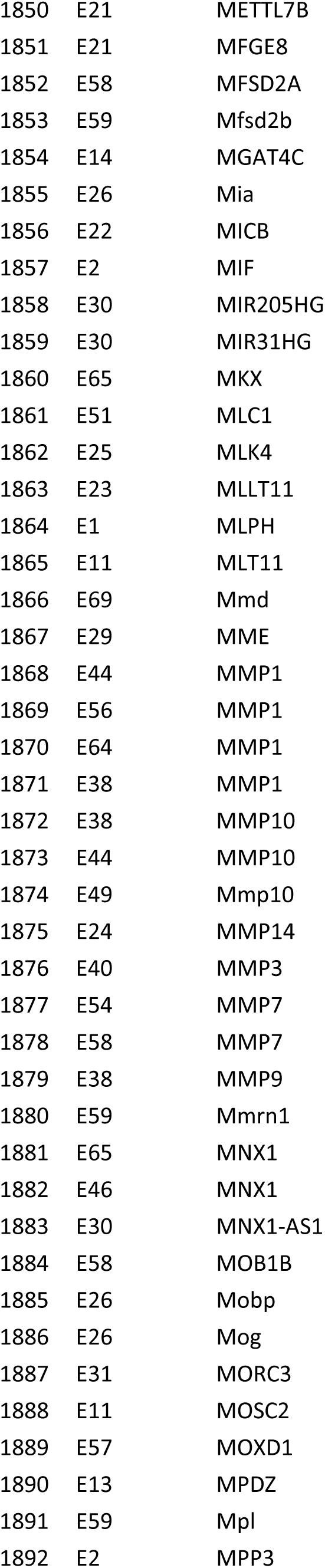
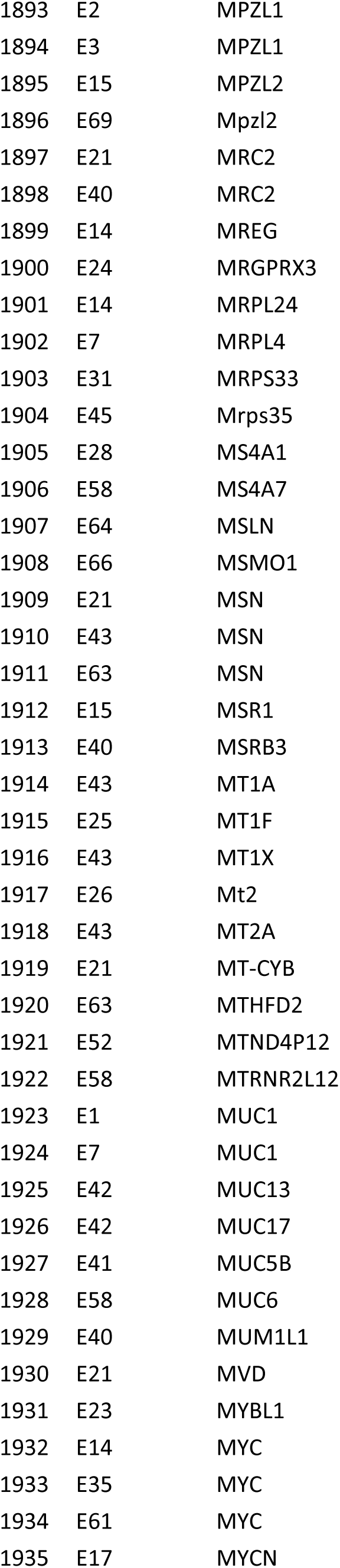
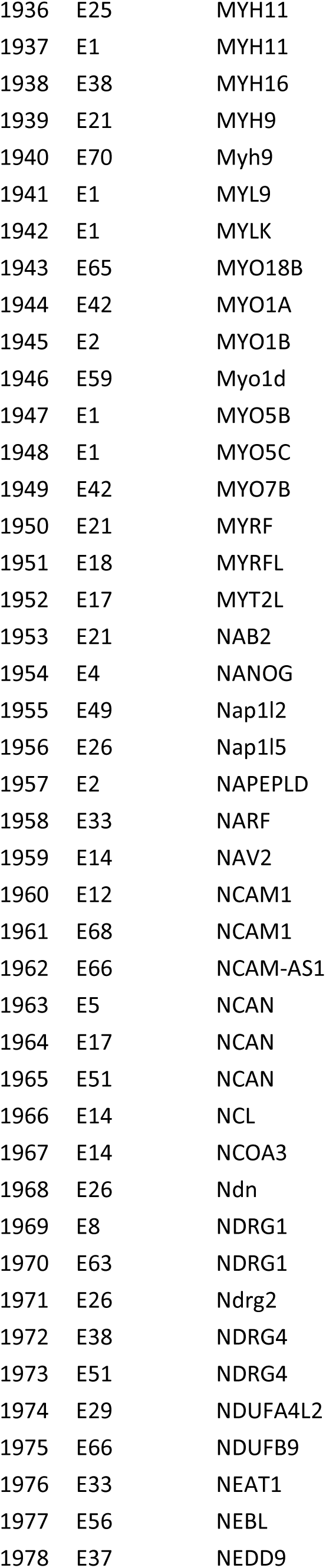
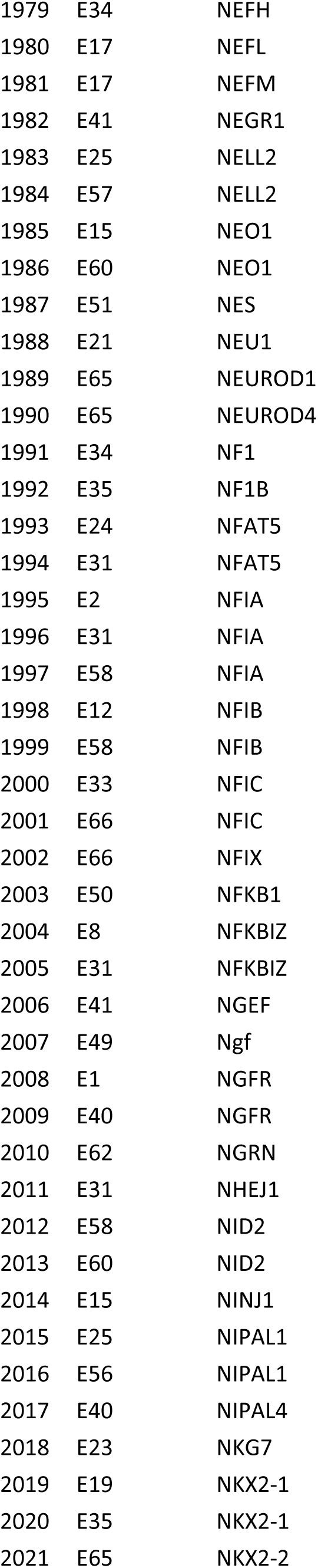
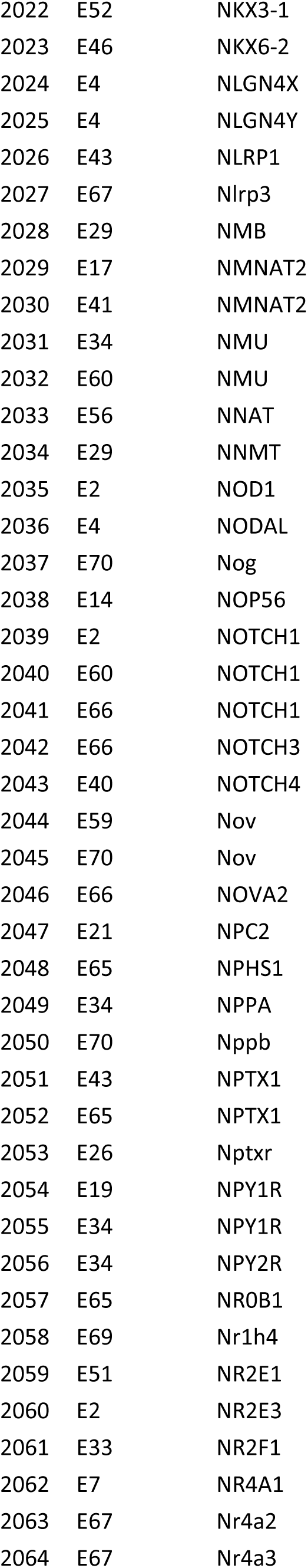
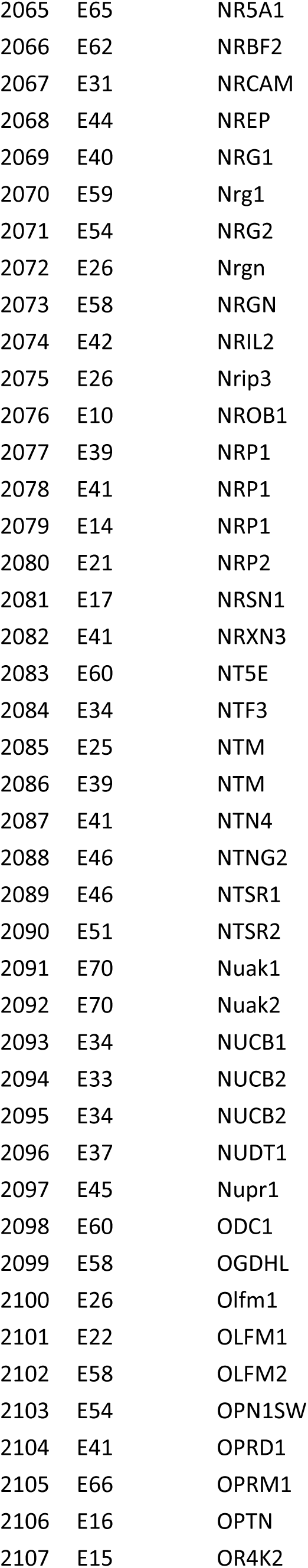
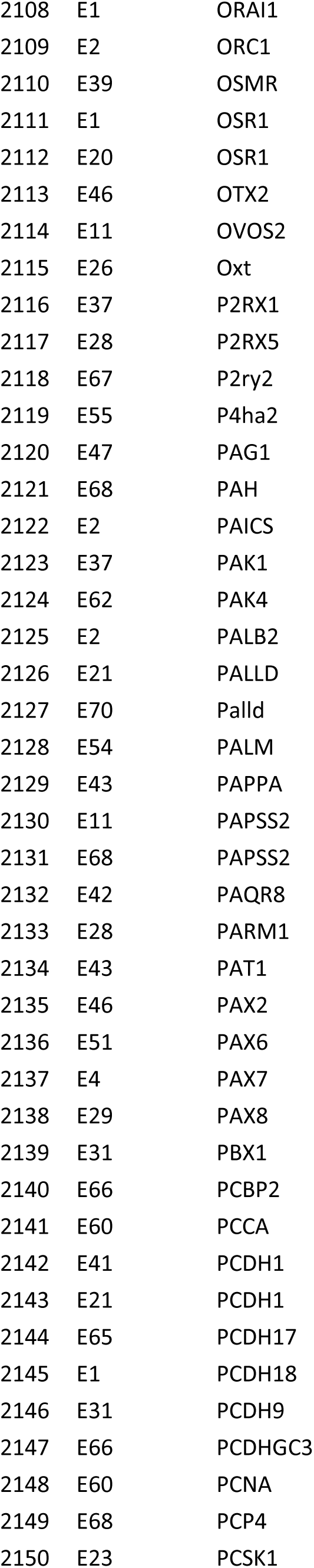
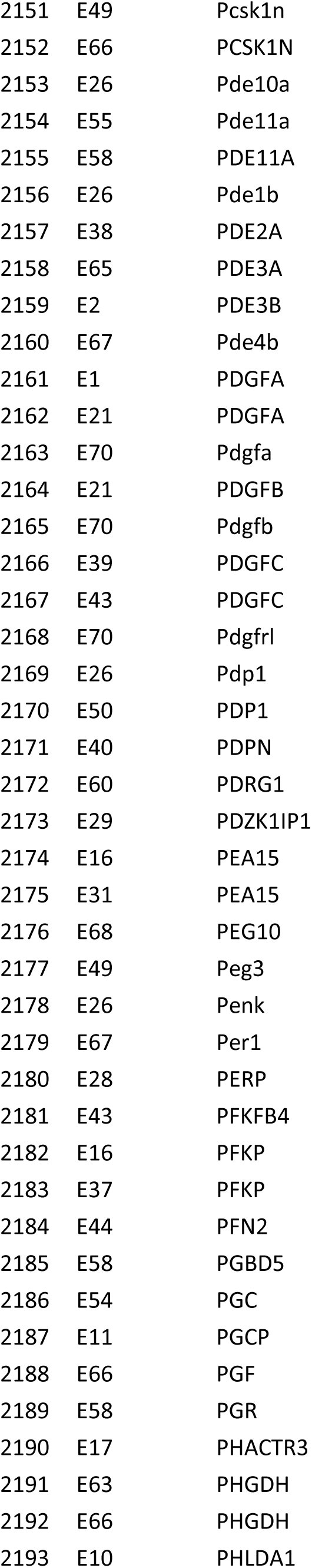
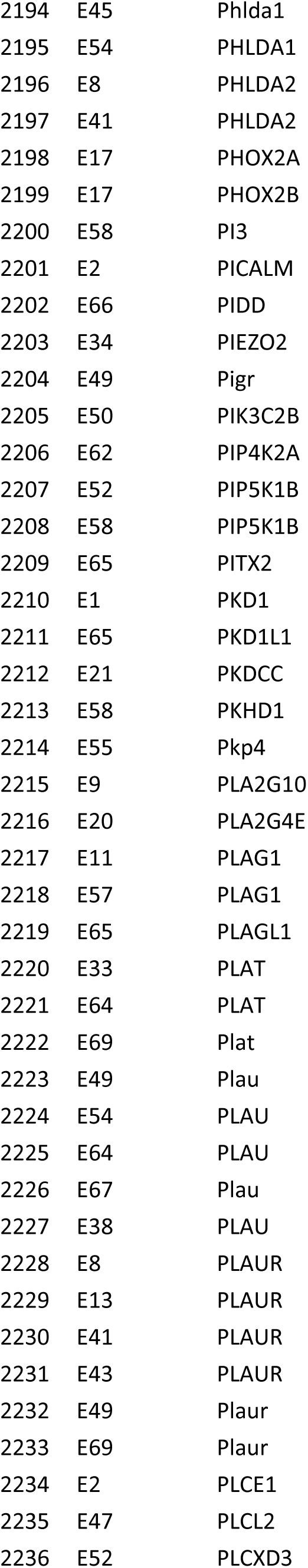
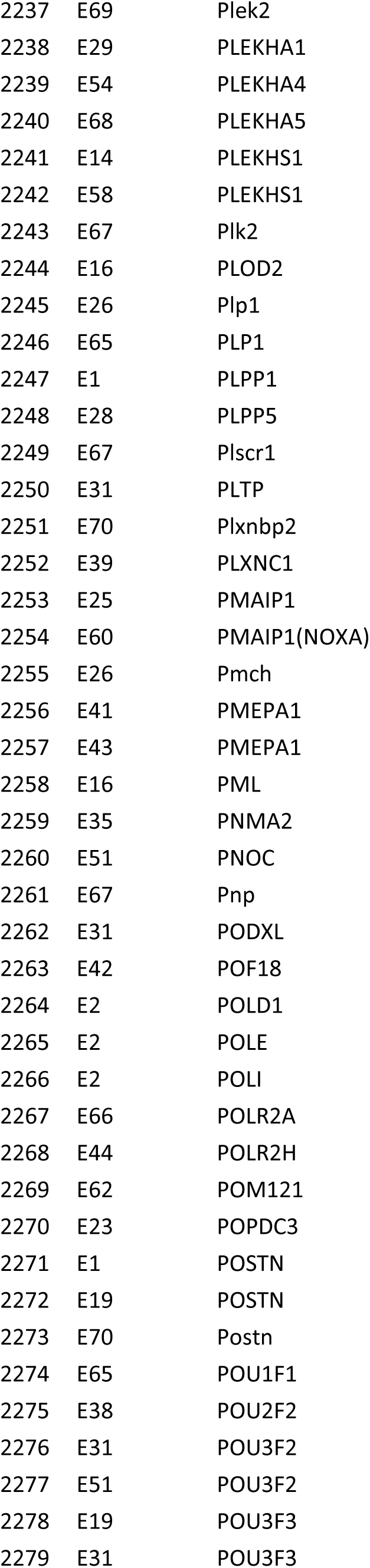
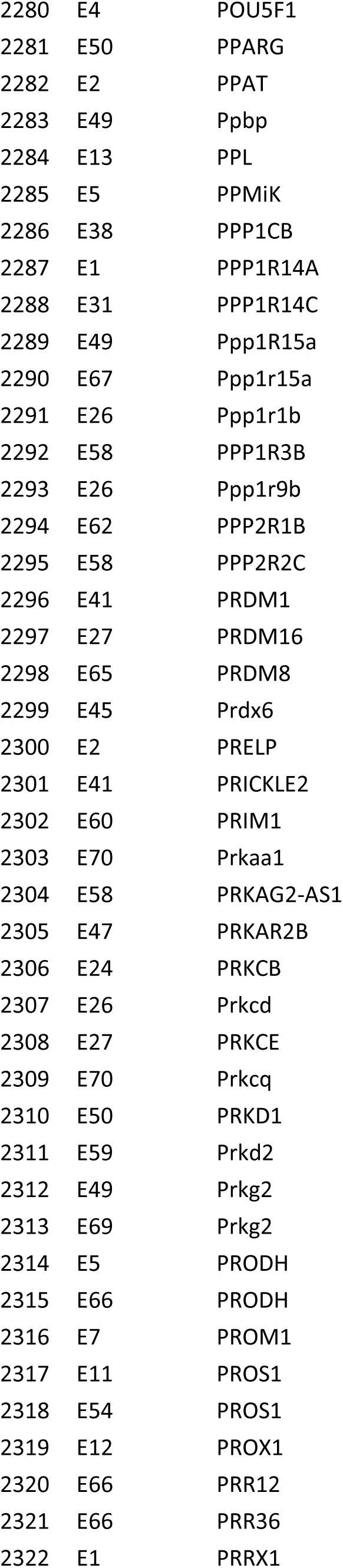
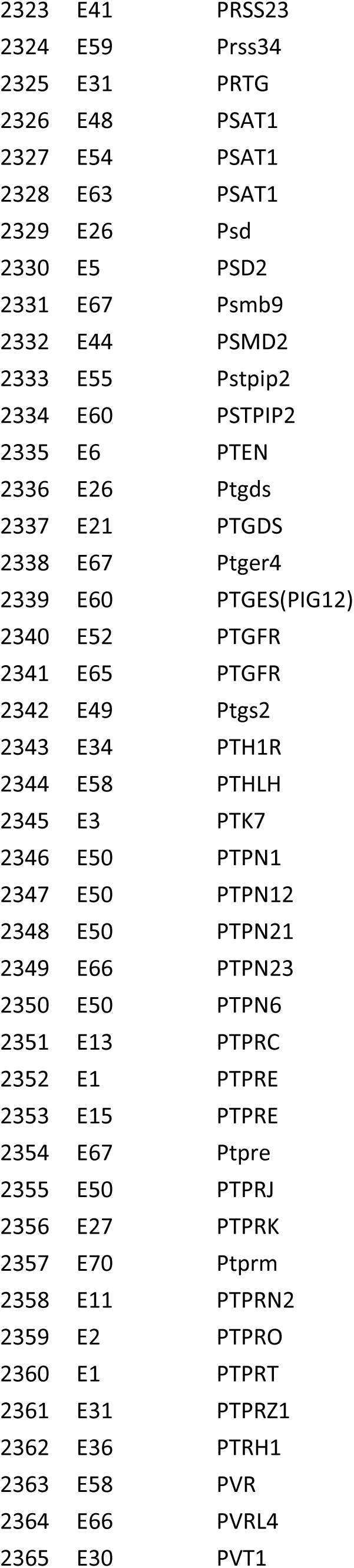
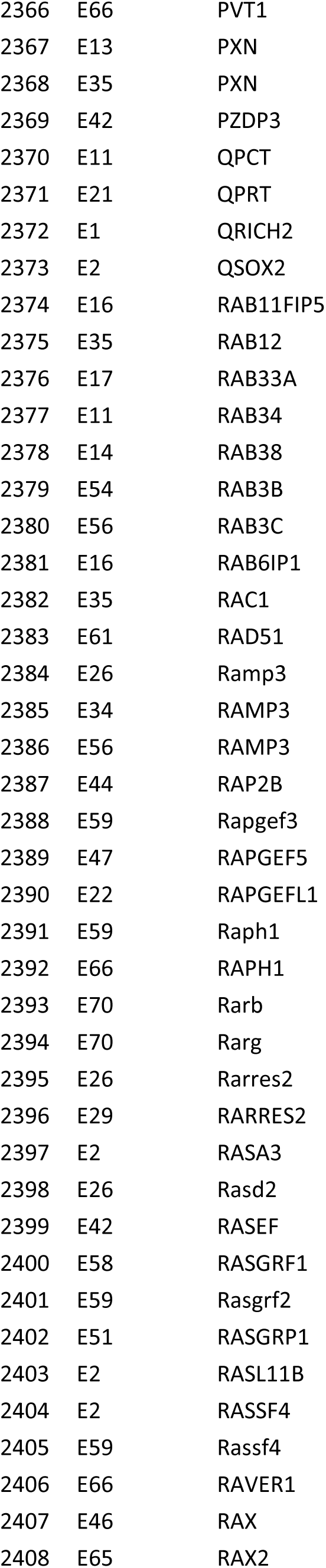
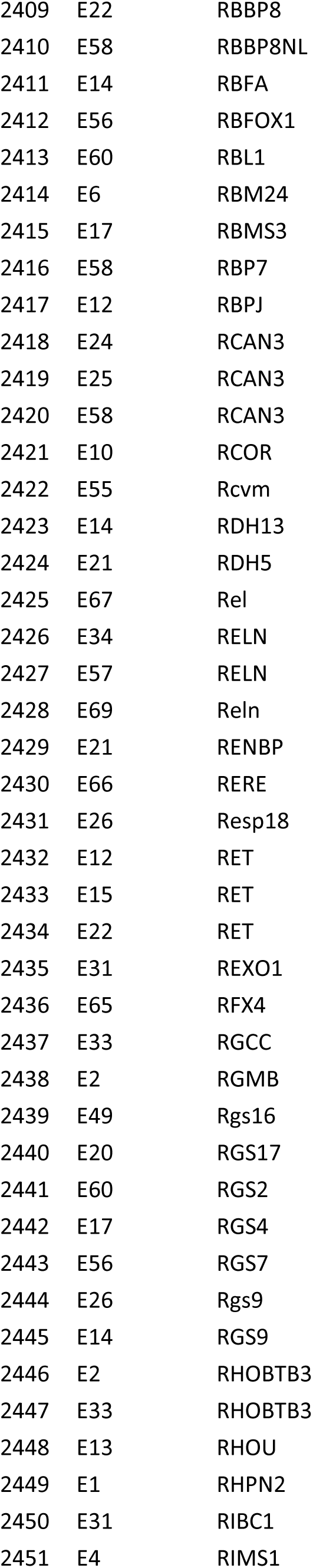
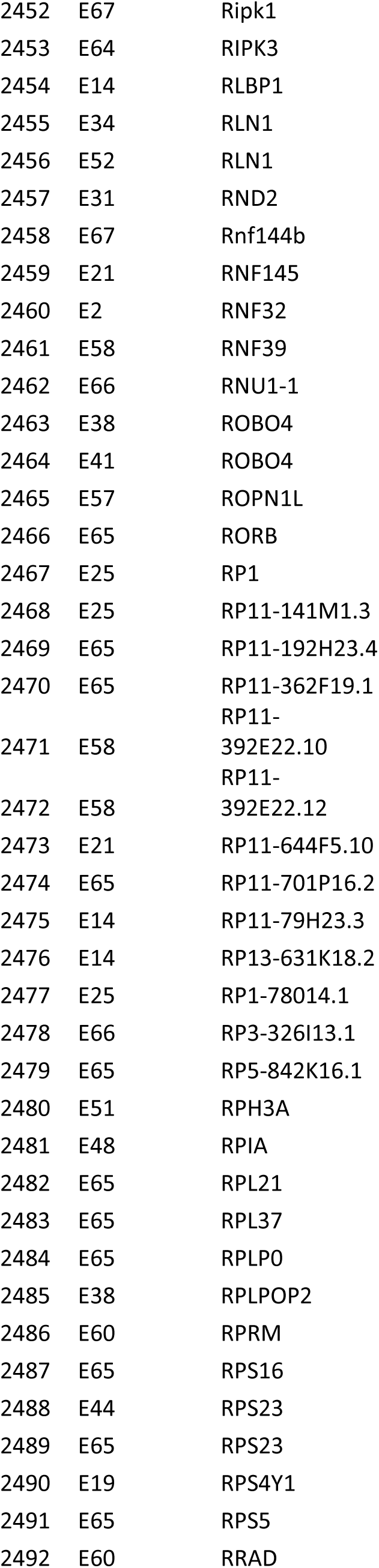
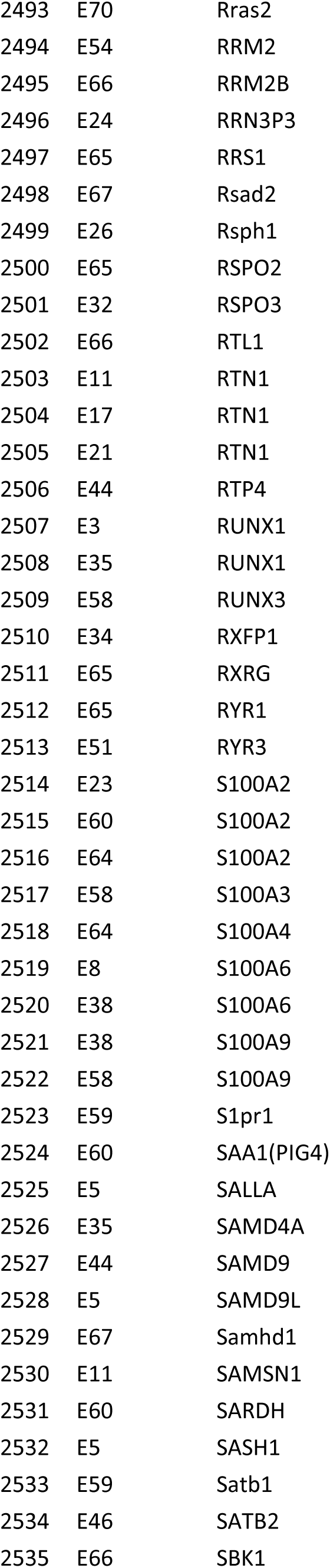
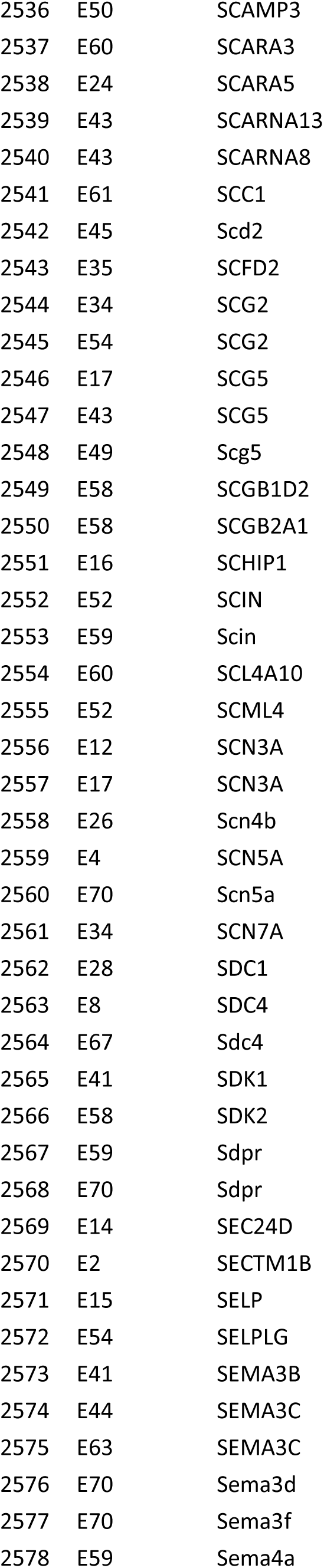
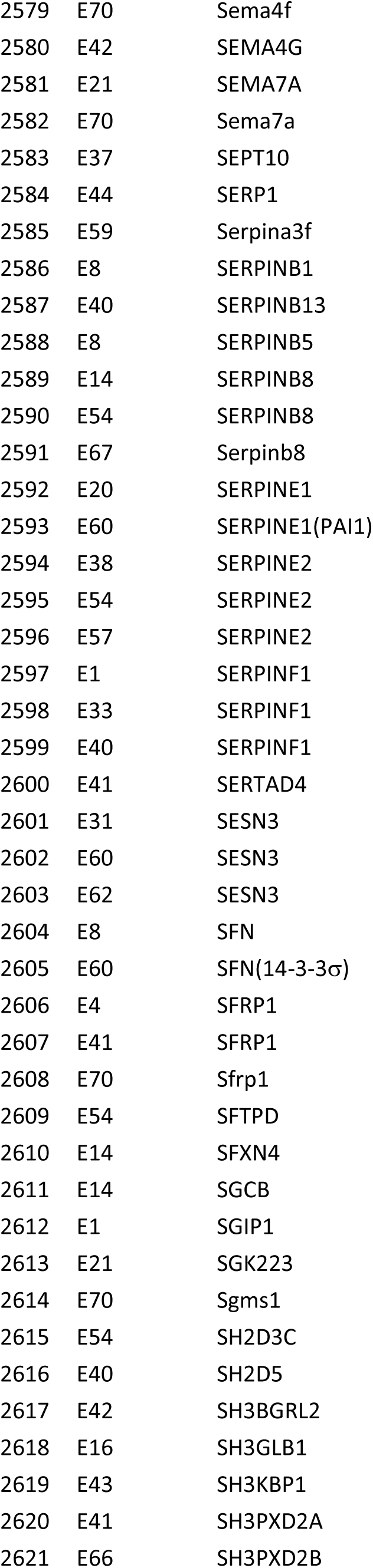
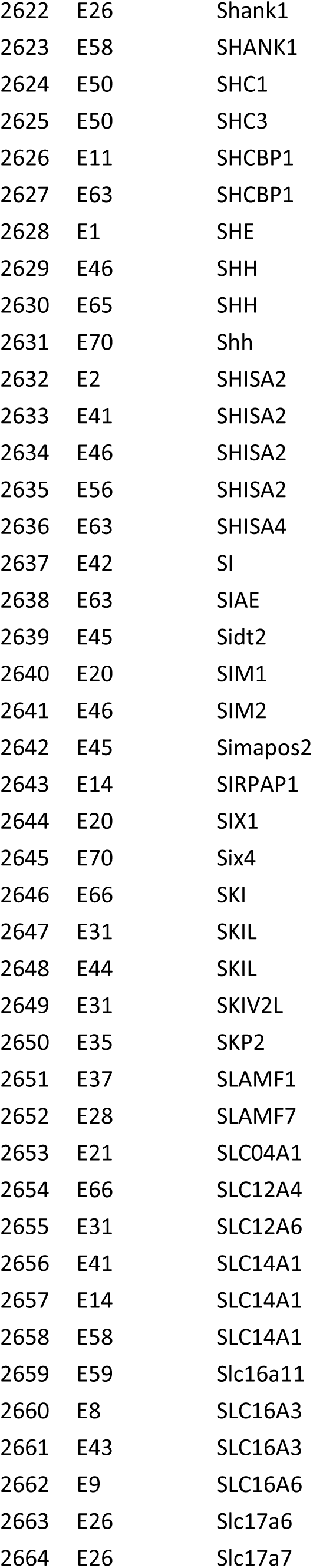
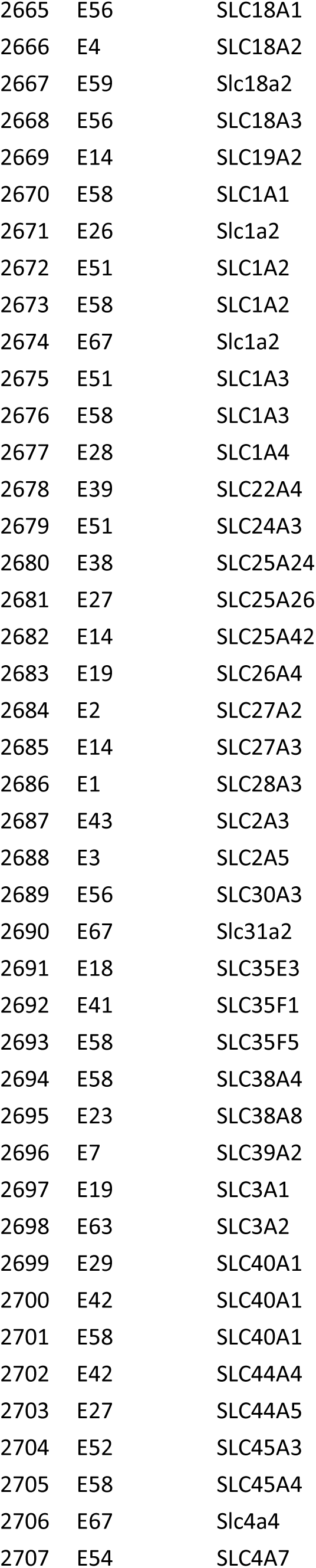
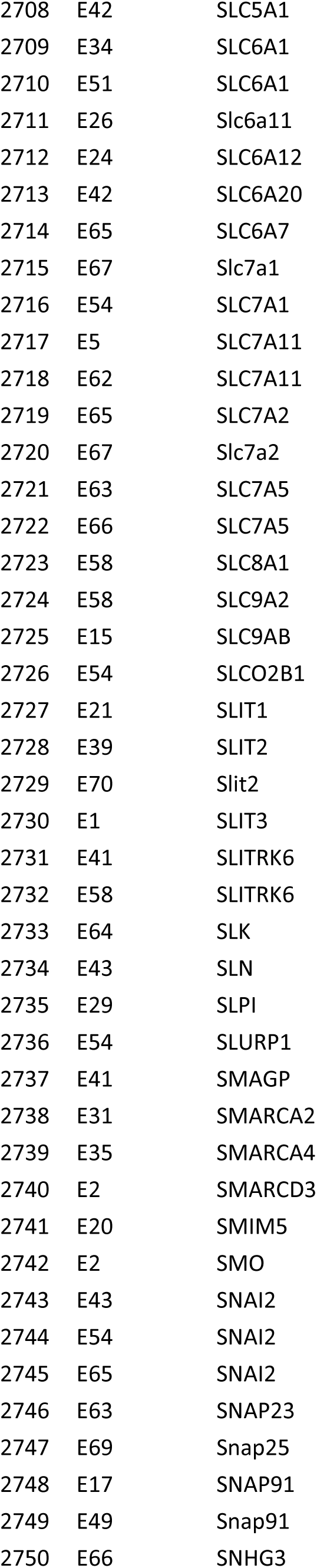
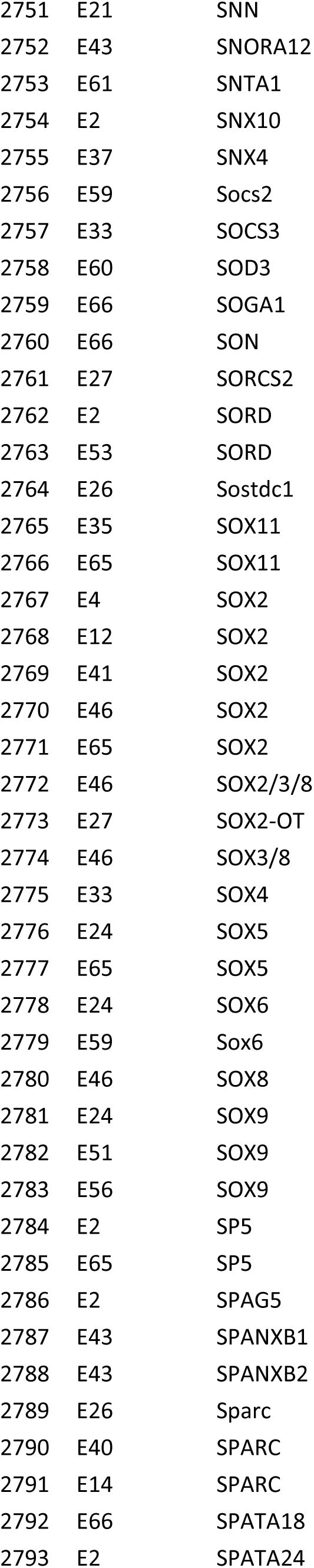
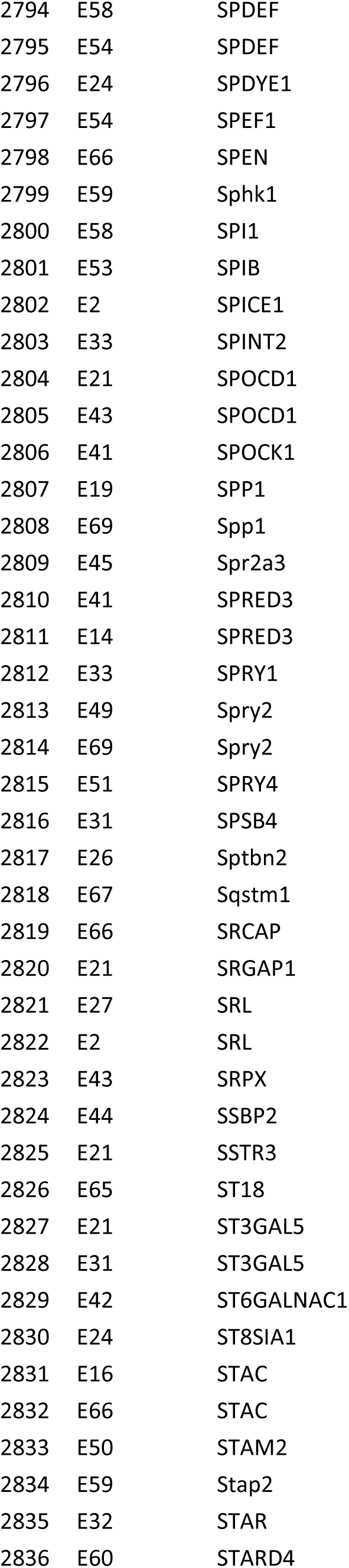
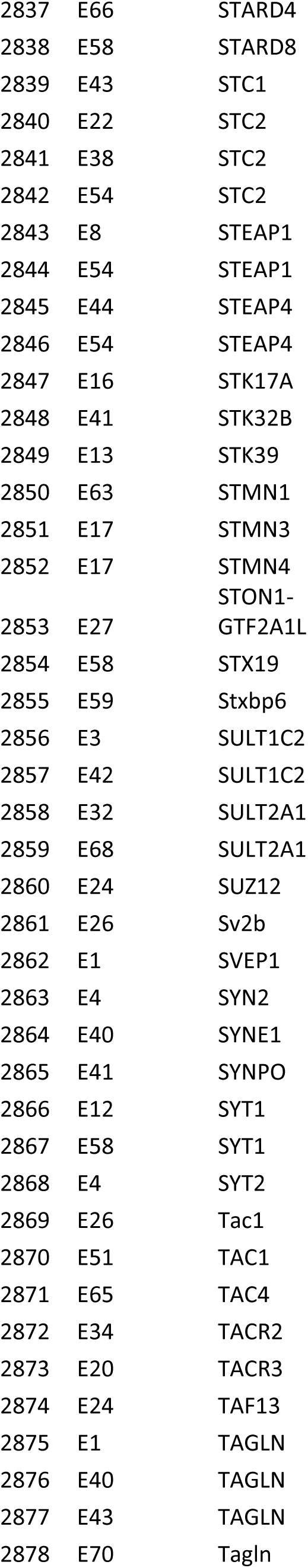
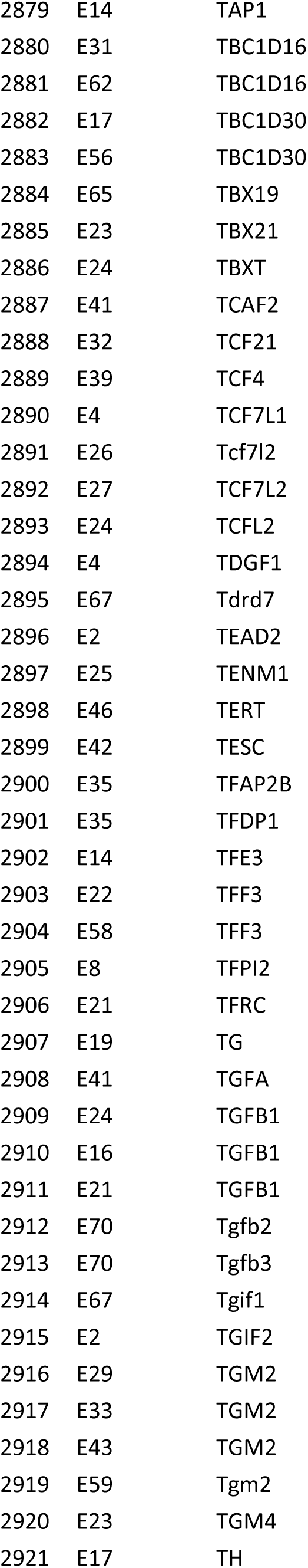
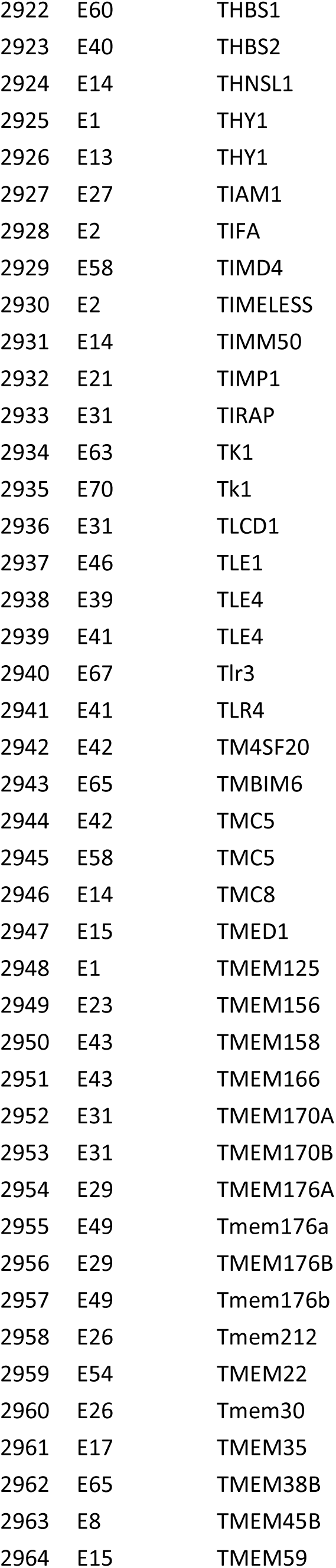
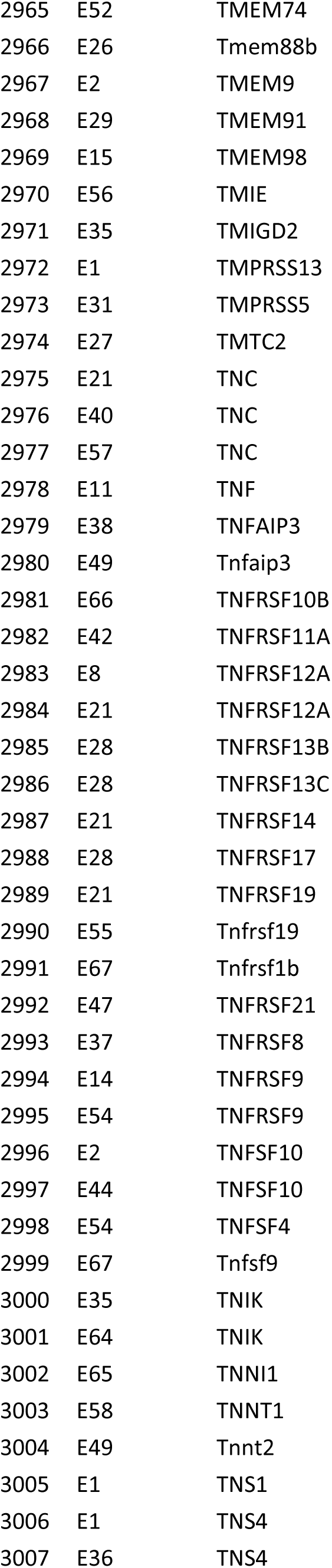
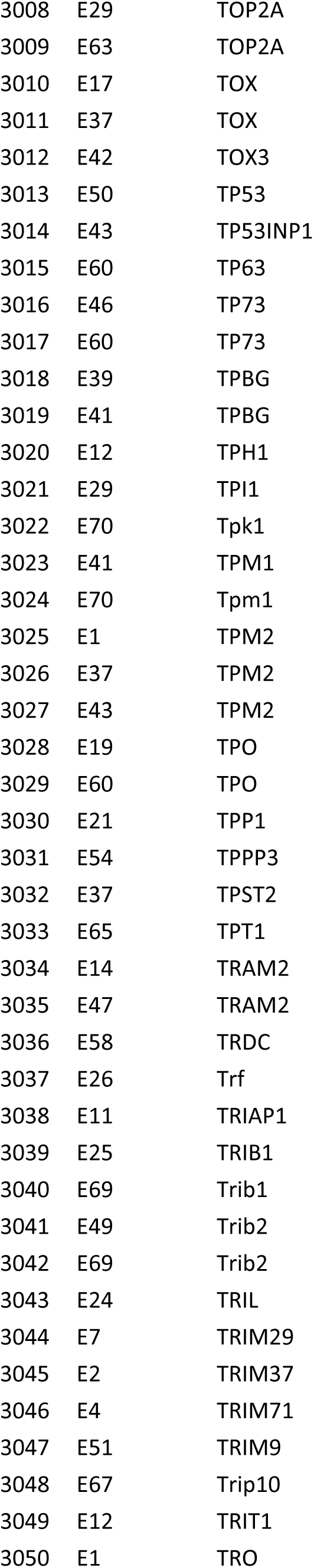
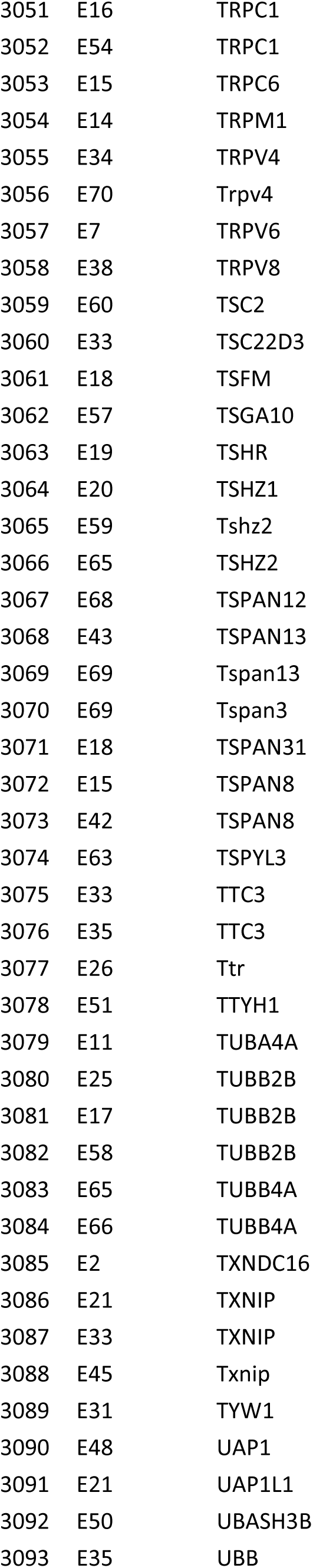
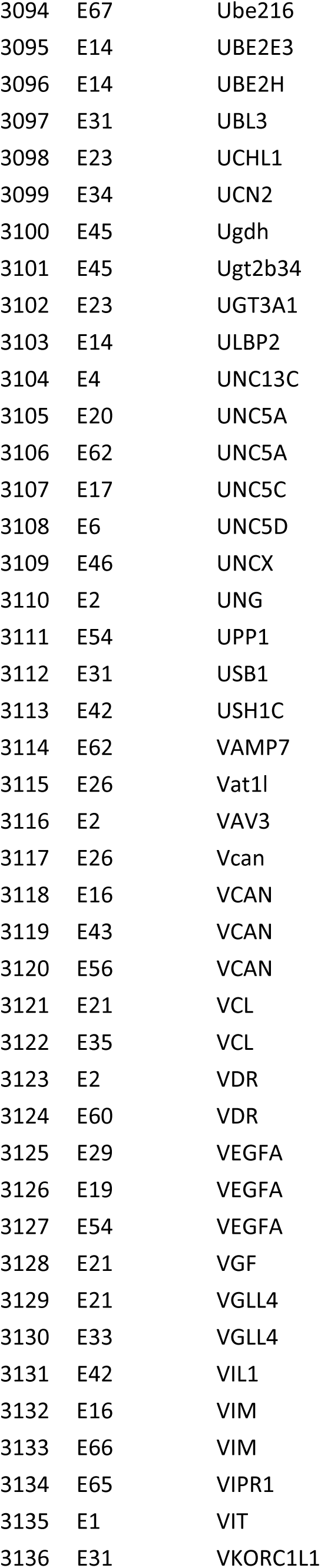
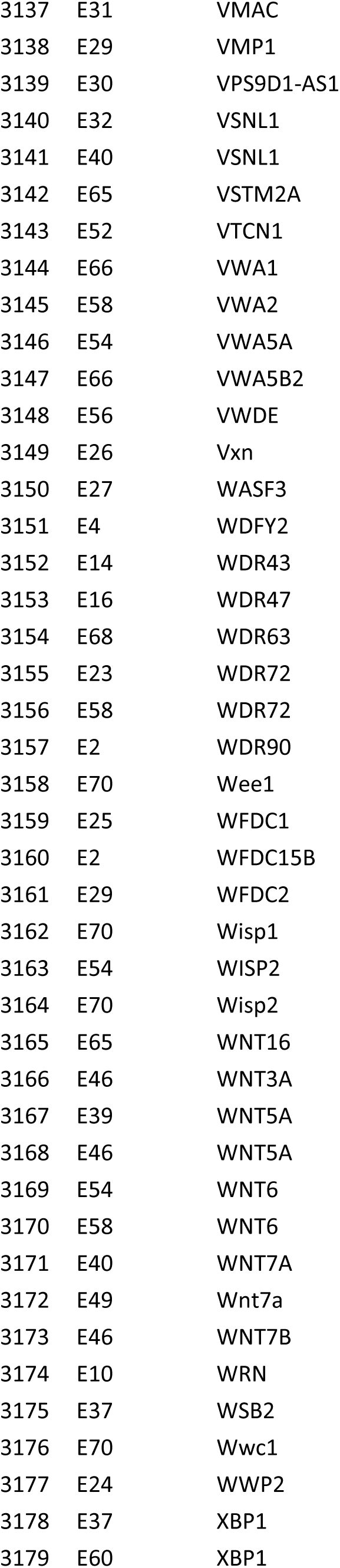
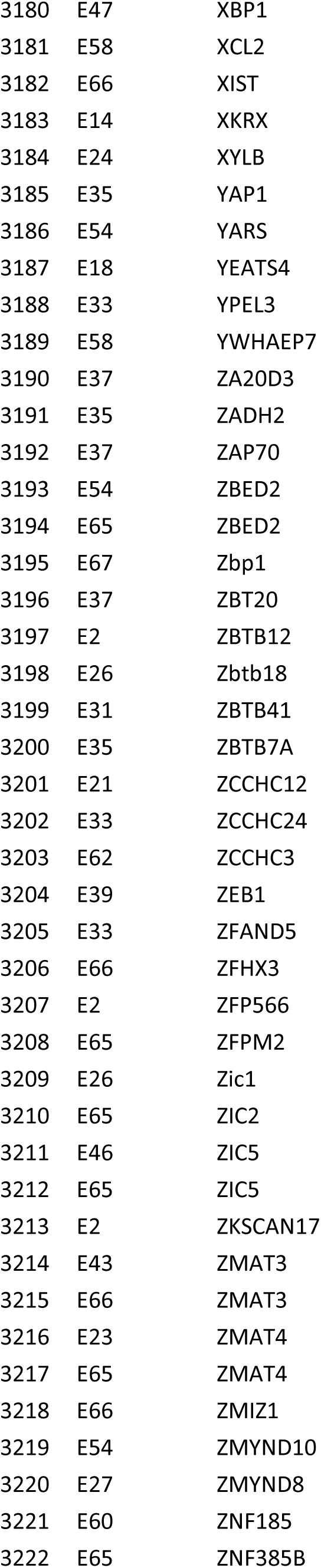
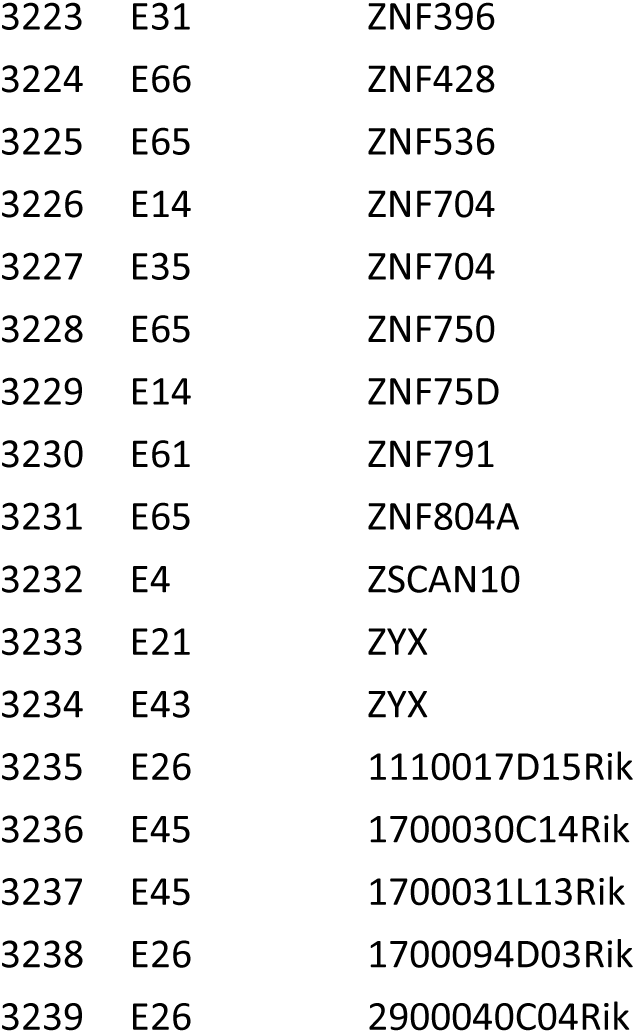

